# Horizontal gene transfer as an indispensable driver for Neocallimastigomycota evolution into a distinct gut-dwelling fungal lineage

**DOI:** 10.1101/487215

**Authors:** Chelsea L. Murphy, Noha H. Youssef, Radwa A. Hanafy, MB Couger, Jason E. Stajich, Y. Wang, Kristina Baker, Sumit S. Dagar, Gareth W. Griffith, Ibrahim F. Farag, TM Callaghan, Mostafa S. Elshahed

**Author notes:** Corresponding author: Mailing address: Oklahoma State University, Department of Microbiology and Molecular Genetics, 1110 S Innovation Way, Stillwater, OK 74074. Both authors contributed equally to this work (Chelsea L. Murphy, Noha H. Youssef).

## Abstract

Survival and growth of the anaerobic gut fungi (AGF, Neocallimastigomycota) in the herbivorous gut necessitate the possession of multiple abilities absent in other fungal lineages. We hypothesized that horizontal gene transfer (HGT) was instrumental in forging the evolution of AGF into a phylogenetically distinct gut-dwelling fungal lineage. Patterns of HGT were evaluated in the transcriptomes of 27 AGF strains, 22 of which were isolated and sequenced in this study, and 4 AGF genomes broadly covering the breadth of AGF diversity. We identified 283 distinct incidents of HGT in AGF transcriptomes, with subsequent gene duplication resulting in an HGT frequency of 2.1-3.6% in AGF genomes. The majority of HGT events were AGF specific (91.5%) and wide (70.7%), indicating their occurrence at early stages of AGF evolution. The acquired genes allowed AGF to expand their substrate utilization range, provided new venues for electron disposal, augmented their biosynthetic capabilities, and facilitated their adaptation to anaerobiosis. The majority of donors were anaerobic fermentative bacteria prevalent in the herbivorous gut. This work strongly indicates that HGT indispensably forged the evolution of AGF as a distinct fungal phylum and provides a unique example of the role of HGT in shaping the evolution of a high rank taxonomic eukaryotic lineage.

**Importance:** The anaerobic gut fungi (AGF) represent a distinct basal phylum lineage (Neocallimastigomycota) commonly encountered in the rumen and alimentary tracts of herbivores. Survival and growth of anaerobic gut fungi in these anaerobic, eutrophic, and prokaryotes dominated habitats necessitates the acquisition of several traits absent in other fungal lineages. This manuscript assesses the role of horizontal gene transfer as a relatively fast mechanism for trait acquisition by the Neocallimastigomycota post sequestration in the herbivorous gut. Analysis of twenty-seven transcriptomes that represent the broad Neocallimastigomycota diversity identified 283 distinct HGT events, with subsequent gene duplication resulting in an HGT frequency of 2.1-3.6% in AGF genomes. These HGT events have allowed AGF to survive in the herbivorous gut by expanding their substrate utilization range, augmenting their biosynthetic pathway, providing new routes for electron disposal by expanding fermentative capacities, and facilitating their adaptation to anaerobiosis. HGT in the AGF is also shown to be mainly a cross-kingdom affair, with the majority of donors belonging to the bacteria. This work represents a unique example of the role of HGT in shaping the evolution of a high rank taxonomic eukaryotic lineage.

## Introduction

Horizontal gene transfer (HGT) is defined as the acquisition, integration, and retention of foreign genetic material into a recipient organism (1). HGT represents a relatively rapid process for trait acquisition; as opposed to gene creation either from preexisting genes (via duplication, fission, fusion, or exon shuffling) or through *de-novo* gene birth from non-coding sequences (2-6). In prokaryotes, the occurrence, patterns, frequency, and impact of HGT on the genomic architecture (7), metabolic abilities (8, 9), physiological preferences (10, 11), and ecological fitness (12) has been widely investigated, and the process is now regarded as a major driver of genome evolution in bacteria and archaea (13, 14). Although eukaryotes are perceived to evolve principally through modifying existing genetic information, analysis of HGT events in eukaryotic genomes has been eliciting increasing interest and scrutiny. In spite of additional barriers that need to be overcome in eukaryotes, e.g. crossing the nuclear membrane, germ line sequestration in sexual multicellular eukaryotes, and epigenetic nucleic acids modifications mechanisms (5, 15), it is now widely accepted that HGT contributes significantly to eukaryotic genome evolution (16, 17). HGT events have convincingly been documented in multiple phylogenetically disparate eukaryotes ranging from the Excavata (18-21), SAR supergroup (22-25), Algae (26), Plants (27), and Opisthokonta (28-31). Reported HGT frequency in eukaryotic genomes ranges from a handful of genes, e.g. (32), to up to 9.6% in Bdelloid rotifers (30).

The kingdom Fungi represents a phylogenetically coherent clade that evolved ≈ 900-1481 Mya from a unicellular flagellated ancestor (33-35). To date, multiple efforts have been reported on the detection and quantification of HGT in fungi. A survey of 60 fungal genomes reported HGT frequencies of 0-0.38% (29), and similar low values were observed in the genomes of five early-diverging pathogenic Microsporidia and Cryptomycota (36). The osmotrophic lifestyle of fungi (37) has typically been regarded as less conducive to HGT compared to the phagocytic lifestyle of several microeukaryotes with relatively higher HGT frequency (38).

The anaerobic gut fungi (AGF, Neocallimastigomycota) represent a phylogenetically distinct basal fungal lineage. The AGF appear to exhibit a restricted distribution pattern, being encountered in the gut of ruminant and non-ruminant herbivorous (39). In the herbivorous gut, the life cycle of the AGF (Figure S1) involves the discharge of motile flagellated zoospores from sporangia in response to animal feeding, the chemotaxis and attachment of zoospores to ingested plant material, spore encystment, and the subsequent production of rhizoidal growth that penetrates and digests plant biomass through the production of a wide array of cellulolytic and lignocellulolytic enzymes.

Survival, colonization, and successful propagation of AGF in the herbivorous gut necessitate the acquisition of multiple unique physiological characteristics and metabolic abilities absent in other fungal lineages. These include, but are not limited to, development of a robust plant biomass degradation machinery, adaptation to anaerobiosis, and exclusive dependence on fermentation for energy generation and recycling of electron carriers (40, 41). Therefore, we hypothesized that sequestration into the herbivorous gut was conducive to the broad adoption of HGT as a relatively faster adaptive evolutionary strategy for niche adaptation by the AGF (Figure S1). Further, since no part of the AGF life cycle occurs outside the animal host and no reservoir of AGF outside the herbivorous gut has been identified (39), then acquisition would mainly occur from donors that are prevalent in the herbivorous gut (Figure S1). Apart from earlier observations on the putative bacterial origin of a few catabolic genes in two AGF isolates (42, 43), and preliminary BLAST-based queries of a few genomes (41, 44), little is currently known on the patterns, determinants, and frequency of HGT in the Neocallimastigomycota. To address this hypothesis, we systematically evaluated the patterns of HGT acquisition in the transcriptomes of 27 AGF strains and 4 AGF genomes broadly covering the breadth of AGF genus-level diversity. Our results document the high level of HGT in AGF in contrast to HGT paucity across the fungal kingdom. The identity of genes transferred, distribution pattern of events across AGF genera, phylogenetic affiliation of donors, and the expansion of acquired genetic material in AGF genomes highlight the role played by HGT in forging the evolution and diversification of the Neocallimastigomycota as a phylogenetically, metabolically, and ecologically distinct lineage in the fungal kingdom.

## Materials and Methods

### Organisms

Type strains of the Neocallimastigomycota are unavailable through culture collections due to their strict anaerobic and fastidious nature, as well as the frequent occurrence of senescence in AGF strains (45). As such, obtaining a broad representation of the Neocallimastigomycota necessitated the isolation of representatives of various AGF genera *de novo*. Samples were obtained from the feces, rumen, or digesta of domesticated and wild herbivores around the city of Stillwater, OK and Val Verde County, Texas (Table 1). Samples were immediately transferred to the laboratory and the isolation procedures usually commenced within 24 hours of collection. A second round of isolation was occasionally conducted on samples stored at −20° C for several weeks (Table 1).

**Table 1:**
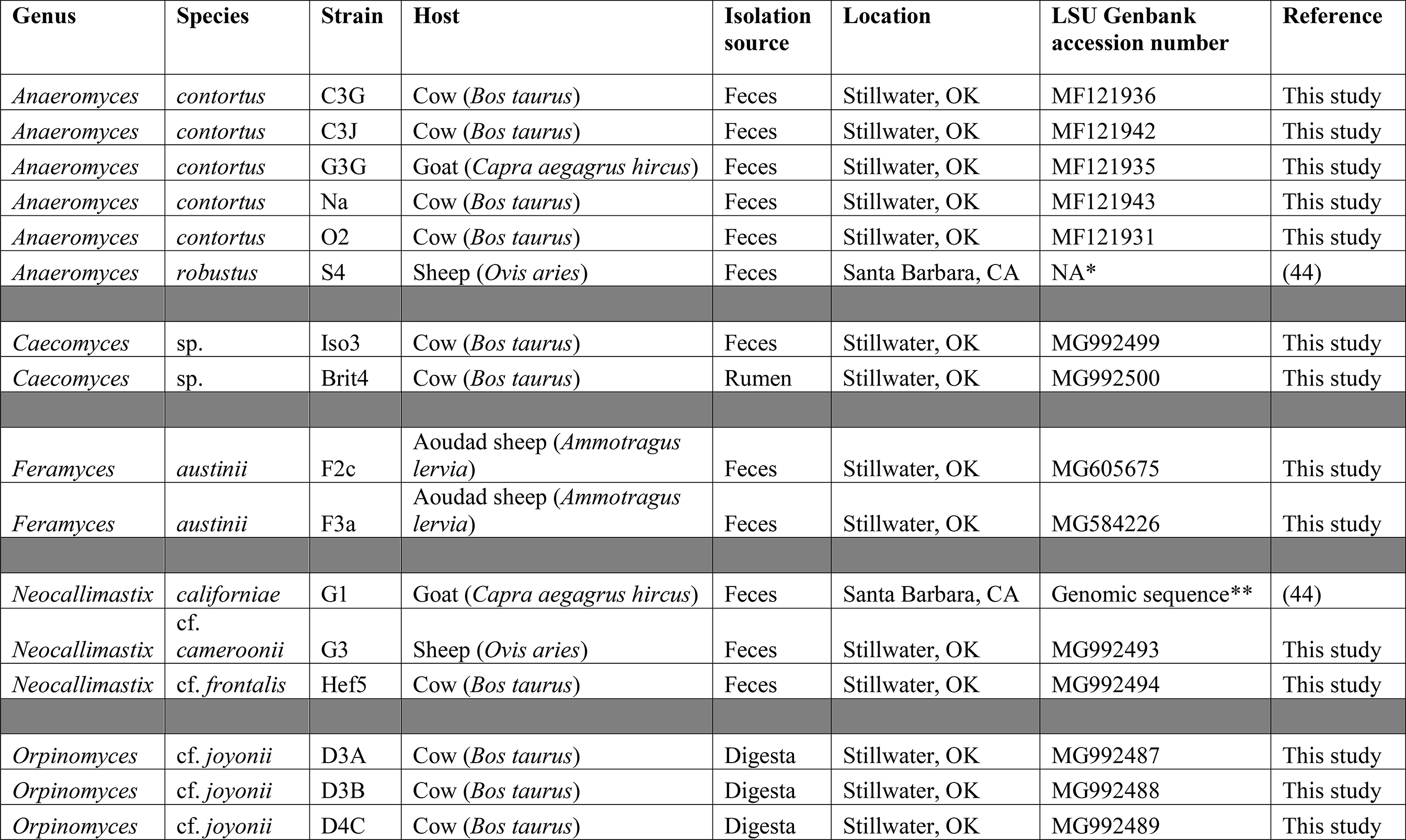
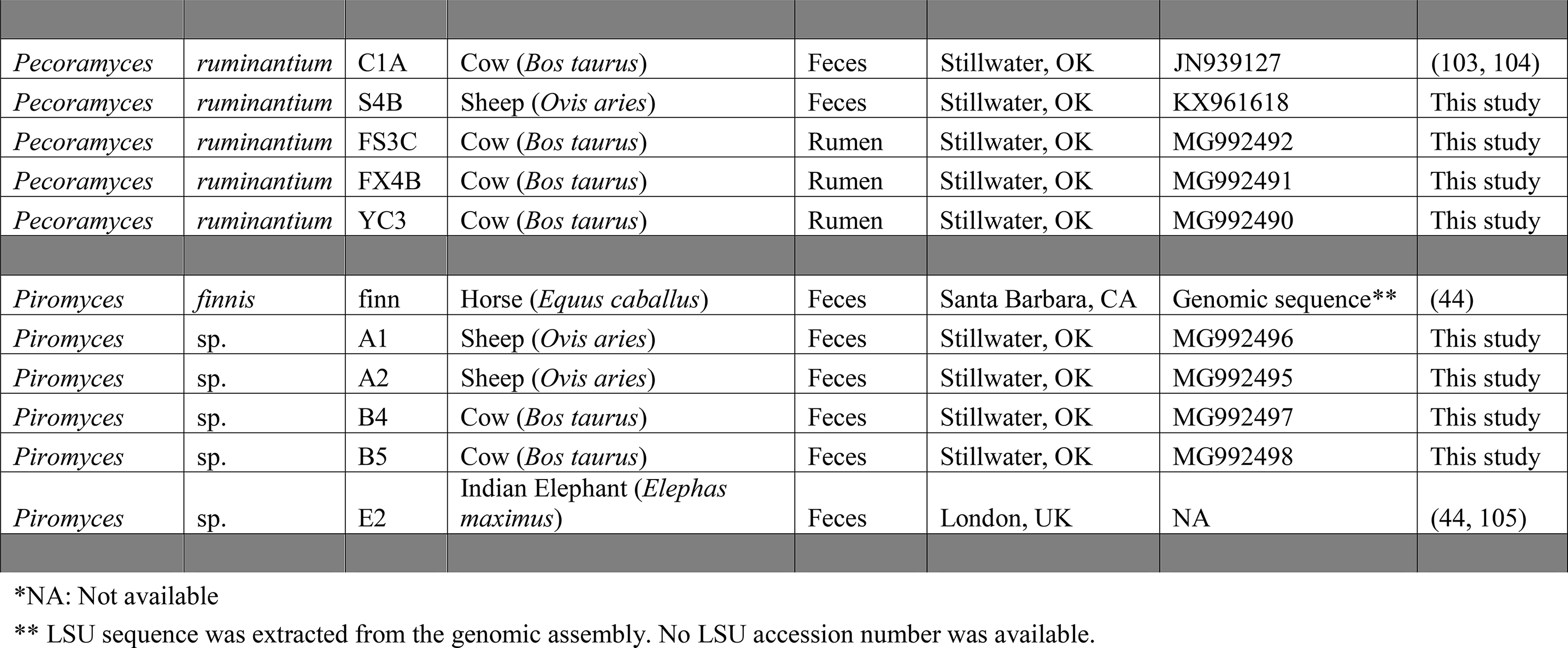
Neocallimastigomycota strains analyzed in this study.

Isolation was performed using a rumen fluid medium reduced by cysteine-sulfide, supplemented with a mixture of kanamycin, penicillin, streptomycin, and chloramphenicol (50 μg/mL, 50 μg/mL, 20 μg/mL, and 50 μg/mL, respectively), and dispensed under a stream of 100% CO_2_ (41, 46). All media were prepared according to the Hungate technique (47), as modified by Balch and Wolfe (48). Cellulose (0.5%), or a mixture of switchgrass (0.5%) and cellobiose (0.5%) were used as carbon sources. Samples were serially diluted and incubated at 39°C for 24-48 h. Colonies were obtained from dilutions showing visible signs of fungal growth using the roll tube technique (49). Colonies obtained were inoculated into liquid media, and a second round of isolation and colony picking was conducted to ensure culture purity. Microscopic examination of thallus growth pattern, rhizoid morphology, and zoospore flagellation, as well LSU rRNA gene D1-D2 domain amplification and sequencing were employed to determine the genus level affiliation of all isolates (46). The cultures were routinely sub-cultured on rumen fluid medium supplemented with antibiotics (to guard against accidental bacterial contamination) and stored on agar media as described previously (41, 50).

### Sequencing and assembly

Transcriptomic sequencing was conducted for twenty-two AGF strains. Sequencing multiple taxa provides stronger evidence for the occurrence of HGT in a target lineage (51), and allows for the identification of phylum-wide versus genus- and species-specific HGT events. Transcriptomic, rather than genomic, sequencing was chosen for AGF-wide HGT identification efforts since enrichment for polyadenylated (poly(A)) transcripts prior to RNA-seq provides a built-in safeguard against possible prokaryotic contamination, an issue that often plagued eukaryotic genome-based HGT detection efforts (52, 53), as well as to demonstrate that HGT genes identified are transcribed in AGF. Further, sequencing and assembly of a large number of Neocallimastigomycota genomes is challenging due to the extremely high AT content in intergenic regions and the extensive proliferation of microsatellite repeats, often necessitating employing multiple sequencing technologies for successful genomic assembly (41, 44).

RNA extraction was conducted as described previously (54). Briefly, fungal biomass was obtained by vacuum filtration and grounded with a pestle under liquid nitrogen. RNA was extracted using Epicentre MasterPure Yeast RNA Purification kit (Epicentre, Madison, WI, USA) and stored in RNase-free TE buffer. Transcriptomic sequencing using Illumina HiSeq2500 2X150bp paired end technology was conducted using the services of a commercial provider (Novogene Corporation, Beijing, China).

RNA-Seq reads were assembled by the de novo transcriptomic assembly program Trinity (55) using previously established protocols (56). All settings were implemented according to the recommended protocol for fungal genomes, with the exception of the absence of the “–jaccard_clip” flag due to the low gene density of anaerobic fungal genomes. The assembly process was conducted on the Oklahoma State University High Performance Computing Cluster as well as the XSEDE HPC Bridges at the Pittsburg Super Computing Center. Quantitative levels for all assembled transcripts were determined using Bowtie2 (57). The program Kallisto was used for quantification and normalization of the gene expression of the transcriptomes (58). All final peptide models predicted were annotated using the Trinotate platform with a combination of homology-based search using BLAST+, domain identification using hmmscan and the Pfam 30.0 database 19 (59), and cellular localization with SignalP 4.0 (60). The twenty-two transcriptomes sequenced in this effort, as well as previously published transcriptomic datasets from *Pecoramyces ruminantium* (41), *Piromyces finnis*, *Piromyces* sp. E2, *Anaeromyces robustus*, and *Neocallimastix californiae* (44) were examined. In each dataset, redundant transcripts were grouped into clusters using CD-HIT-EST with identity parameter of 95% (-c 0.95). The obtained non-redundant transcripts from each analyzed transcriptome were subsequently used for peptide and coding sequence prediction using the TransDecoder with a minimum peptide length of 100 amino acids (http://transdecoder.github.io). Assessment of transcriptome coverage per strain was conducted using BUSCO (61).

### HGT identification

A combination of BLAST similarity searches, comparative similarity index (HGT index, *h*_*U*_), and phylogenetic analyses were conducted to identify HGT events in the analyzed transcriptomic datasets (Figure 1). We define an HGT event as the acquisition of a foreign gene/Pfam by AGF from a single lineage/donor. All predicted peptides were queried against Uniprot databases (downloaded May 2017) each containing both reviewed (Swiss-Prot) and unreviewed (TrEMBL) sequences. The databases encompassed nine different phylogenetic groups; Bacteria, Archaea, Viridiplantae, Opisthokonta-Chaonoflagellida, Opisthokonta-Fungi (without Neocallimastigomycota representatives), Opisthokonta-Metazoa, Opisthokonta-Nucleariidae and Fonticula group, all other Opisthokonta, and all other non-Opisthokonta-non-Viridiplantae Eukaryota. For each peptide sequence, the bit score threshold and HGT index *h*_*U*_ (calculated as the difference between the bit-scores of the best non-fungal and the best Dikarya fungal matches) were determined. Peptide sequences that satisfied the criteria of having a BLASTP bit-score against a non-fungal database that was >100 (i.e. 2^-100^ chance of random observation) and an HGT index *h*_*U*_ that was ≥30 were considered HGT candidates and subjected to additional phylogenetic analysis. We chose to work with bit-score rather than the raw scores since the bit-score measures sequence similarity independent of query sequence length and database size. This is essential when comparing hits from databases with different sizes (for example, the Bacteria database contained 83 million sequences while the Choanoflagellida database contained 21 thousand sequences). We chose an *h*_*U*_ value of ≥30 (a difference of bit-score of at least 30 between the best non-fungal hit and the best fungal hit to an AGF sequence) previously suggested and validated (62, 63) as the best tradeoff between sensitivity and specificity. Since the bit-score is a logarithmic value that describes sequence similarity, a bit-score > 30 ensure that the sequence aligned much better to the non-fungal hit than it did to the fungal hit.

**Figure 1.**
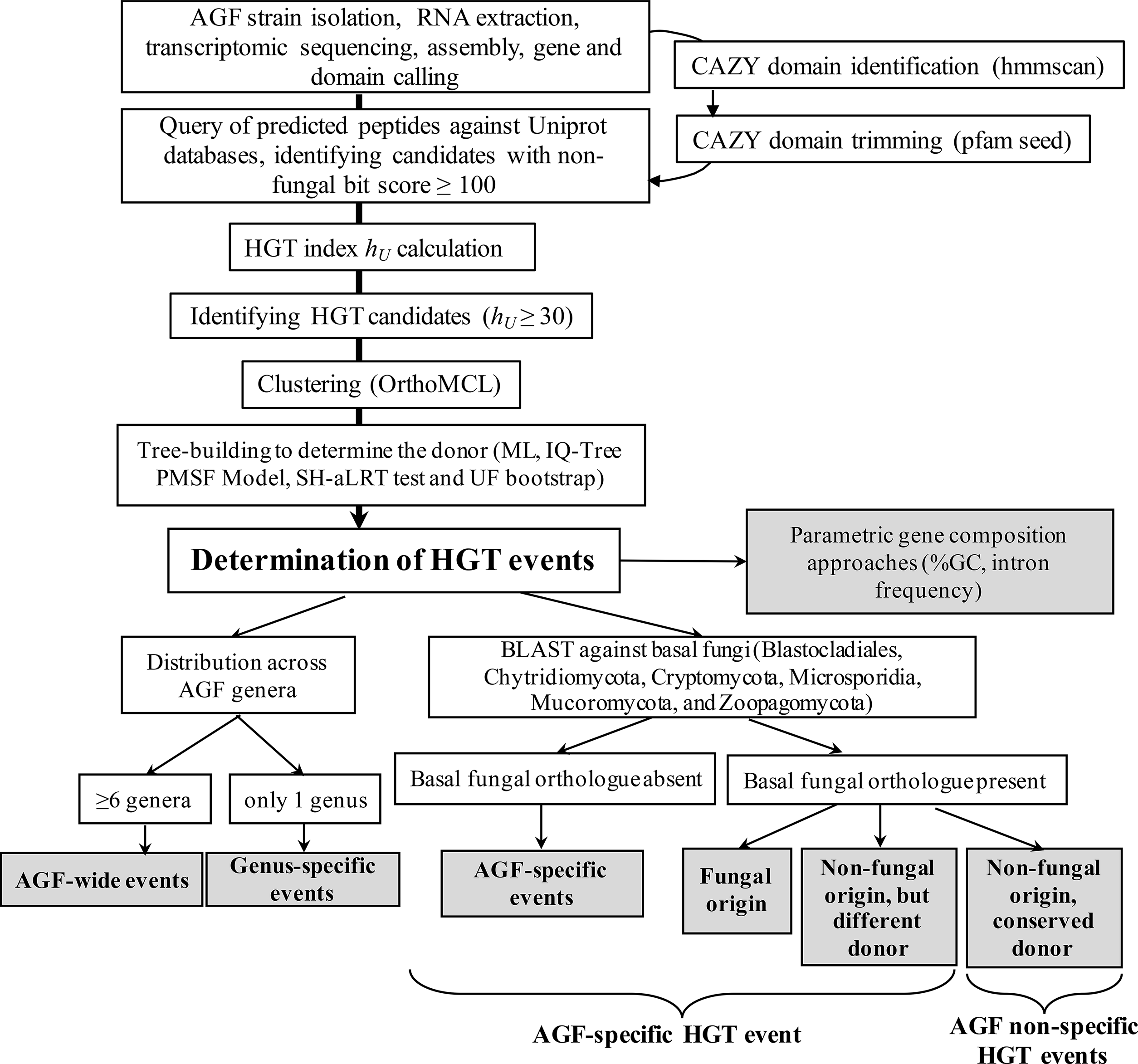
Workflow diagram describing the procedure employed for identification HGT events in Neocallimastigomycota datasets analyzed in this study.

The identified HGT candidates were modified by removing all CAZyme-encoding sequences (due to their multi-modular nature, see below) and further clustered into orthologues using OrthoMCL (64). Orthologues obtained were subjected to detailed phylogenetic analysis to confirm HGT occurrence as well as to determine the potential donor. Each Orthologue was queried against the nr database using web Blastp (65) under two different settings: once against the full nr database and once against the Fungi (taxonomy ID: 4751) excluding the Neocallimastigomycetes (Taxonomy ID: 451455). The first 250 hits obtained using these two Blastp searches with an e-value below e^-10^ were downloaded and combined in one fasta file. Datasets were reduced by removing duplicate sequences as well as redundant sequences from one organism. AGF and reference sequences were aligned using the standalone Clustal Omega (66). Alignments were viewed and manually curated in Mega (67). The alignments were used to generate guide trees in FastTree under the LG model (68). Guide trees were in turn used as input to IQ-tree (69) to generate maximum likelihood trees under the posterior mean site frequency method (PMSF), shown before to ameliorate long-branch attraction artifacts (70). Both the (-alrt 1000) option for performing the Shimodaira–Hasegawa approximate likelihood ratio test (SH-aLRT), as well as the (-bb 1000) option for ultrafast bootstrap (UFB) (71) were added to the IQ-tree command line. This resulted in the generation of phylogenetic trees with two support values (SH-aLRT and UFB) on each branch. Candidates that showed a nested phylogenetic affiliation that was incongruent to organismal phylogeny with strong SH-aLRT and UFB supports were deemed horizontally transferred.

### Identification of HGT events in carbohydrate active enzymes (CAZymes) transcripts

In AGF genomes, carbohydrate active enzymes (CAZymes) are often encoded by large multi-module genes with multiple adjacent CAZyme or non-CAZyme domains (41, 44). A single gene can hence harbor multiple CAZyme pfams of different (fungal or non-fungal) origins (41, 44). As such, our initial efforts for HGT assessment in CAZyme-encoding transcripts using an entire gene/ transcript strategy yielded inaccurate results since similarity searches only identified pfams with the lowest e-value or highest number of copies, while overlooking additional CAZyme pfams in the transcript (Figure S2). To circumvent the multi-modular nature of AGF CAZyme transripts, we opted for the identification of CAZyme HGT events on trimmed domains, rather than entire transcript. CAZyme-containing transcripts (Glycoside hydrolases (GHs), Polysaccharide lyases (PLs), and Carbohydrate Esterases (CEs)) were first identified by searching the entire transcriptomic datasets against the dbCAN hidden markov models V5 (72) (downloaded from the dbCAN web server in September 2016) using the command hmmscan in standalone HMMER. For each CAZy family identified, predicted peptides across all transcriptomic datasets were grouped in one fasta file that was then amended with the corresponding Pfam seed sequences (downloaded from the Pfam website (http://pfam.xfam.org/) in March 2017). Sequences were aligned using the standalone Clustal Omega (66) to their corresponding Pfam seeds. Using the Pfam seed sequences as a guide for the start and end of the domain, aligned sequences were then truncated in Jalview (73). Truncated transcripts with an identified CAZy domain were again compared to the pfam database (74) using hmmscan (75) to ensure correct assignment to CAZy families and accurate domain trimming. These truncated peptide sequences were then analyzed to pinpoint incidents of HGT using the approach described above.

### Neocallimastigomycota-specific versus non-specific HGT events

To determine whether an identified HGT event (i.e. foreign gene acquisition from a specific donor) is specific to the phylum Neocallimastigomycota; the occurrence of orthologues (30% identity, >100 amino acids alignment) of the identified HGT genes in basal fungi, i.e. members of Blastocladiales, Chytridiomycota, Cryptomycota, Microsporidia, Mucormycota, and Zoopagomycota, as well as the putative phylogenetic affiliation of these orthologues, when encountered, were assessed. HGT events were judged to be Neocallimastigomycota-specific if: 1. orthologues were absent in all basal fungal genomes, 2. orthologues were identified in basal fungal genomes, but these orthologues were of clear fungal origin, or 3. orthologues were identified in basal fungal genomes and showed a non-fungal phylogenetic affiliation, but such affiliation was different from that observed in the Neocallimastigomycota. On the other hand, events were judged to be non-specific to the Neocallimastigomycota if phylogenetic analysis of basal fungal orthologues indicated a non-fungal origin with a donor affiliation similar to that observed in the Neocallimastigomycota (Figure 1).

### Mapping HGT events to available AGF genomes

HGT events identified in AGF datasets examined (both CAZy and non-CAZy events) were mapped onto currently available AGF genome assemblies (41, 44) (Genbank accession numbers ASRE00000000.1, MCOG00000000.1, MCFG00000000.1, MCFH00000000.1). The duplication and expansion patterns, as well as GC content, and intron distribution were assessed in all identified genes. Averages were compared to AGF genome average using Student t-test to identify possible deviations in such characteristics as often observed with HGT genes (76). To avoid any bias the differences in the number of genes compared might have on the results, we also compared the GC content, codon usage, and intron distribution averages for the identified genes to a subset of an equal number of randomly chosen genes from AGF genomes. We used the MEME Suite’s fasta-subsample function (http://meme-suite.org/doc/fasta-subsample.html) to randomly select an equal number of genes from the AGF genomes.

### Validation of HGT-identification pipeline using previously published datasets

As a control, the frequency of HGT occurrence in the genomes of a filamentous ascomycete (*Colletotrichum graminicola,* GenBank Assembly accession number GCA_000149035.1), and a microsporidian (*Encephalitozoon hellem,* GenBank Assembly accession number GCA_000277815.3) were determined using our pipeline (Table S1); and the results were compared to previously published results (36, 77).

### Guarding against false positive HGT events due to contamination

Multiple safeguards were taken to ensure that the frequency and incidence of HGT reported here are not due to bacterial contamination of AGF transcripts. These included: 1. Application of antiobiotics in all culturing procedures as described above. 2. Utilization of transcriptomes rather than genomes selects for eukaryotic polyadenylated (poly(A)) transcripts prior to RNA-seq as a built-in safeguard against possible prokaryotic contamination. 3. Mapping HGT transcripts identified to genomes generated in prior studies and confirming the occurrence of introns in the majority of HGT genes identified. 4. Applying a threshold where only transcripts identified in >50% of transcriptomic assemblies from a specific genus are included and 5. The exclusion of HGT events showing suspiciously high (>90%) sequence identity to donor sequences.

In addition, recent studies have demonstrated that GenBank-deposited reference genomes (52) and transcriptomes (78) of multicellular organisms are often plagued by prokaryotic contamination. The occurrence of prokaryotic contamination in reference donors’ genomes/transcriptomes could lead to false positive HGT identification, or incorrect HGT assignments. To guard against any false positive HGT event identification due to possible contamination in reference datasets, sequence data from potential donor reference organisms were queried using blast, and their congruence with organismal phylogeny was considered a prerequisite for inclusion of an HGT event.

### Data accession

Sequences of individual transcripts identified as horizontally transferred are deposited in GenBank under the accession number MH043627-MH043936, and MH044722-MH044724. The whole transcriptome shotgun sequences were deposited in GenBank under the BioProject PRJNA489922, and Biosample accession numbers SAMN09994575-SAMN09994596. Transcriptomic assemblies were deposited in the SRA under project accession number SRP161496. Alignments, as well as Newick tree files for all HGT genes are provided as Supplementary datasets. Trees of HGT events discussed in the results and discussion sections are presented in the supplementary document (S5-S45).

## Results

### Isolates

The transcriptomes of 22 different isolates were sequenced. These isolates belonged to six out of the nine currently described AGF genera: *Anaeromyces* (n=5), *Caecomyces* (n=2), *Neocallimastix* (n=2), *Orpinomyces* (n=3), *Pecoramyces* (n=4), *Piromyces* (n=4), as well as the recently proposed genus *Feramyces* (n=2) (79) (Table 1, Supplementary Fig. 3). Out of the three AGF genera not included in this analysis, two are currently represented by a single strain that was either lost (genus *Oontomyces* (80)), or appears to exhibit an extremely limited geographic and animal host distribution (genus *Buwchfawromyces* (81)). The third unrepresented genus (*Cyllamyces*) has recently been suggested to be phylogenetically synonymous with *Caecomyces* (82). As such, the current collection is a broad representation of currently described AGF genera.

### Sequencing

Transcriptomic sequencing yielded 15.2-110.8 million reads (average, 40.87) that were assembled into 31,021-178,809 total transcripts, 17,539-132,141 distinct transcripts (clustering at 95%), and 16,500-70,061 predicted peptides (average 31,611) (Table S2). Assessment of transcriptome coverage using BUSCO (61) yielded high completion values (82.76-97.24%) for all assemblies (Table S1). For strains with a sequenced genome, genome coverage (percentage of genes in a strain’s genome for which a transcript was identified) ranged between 70.9-91.4% (Table S2).

### HGT events

A total of 12,786 orthologues with a non-fungal bit score > 100, and an HGT index > 30 were identified. After removing orthologues occurring only in a single strain or in less than 50% of isolates belonging to the same genus, 2147 events were further evaluated. Phylogenetic analysis could not confirm the HGT nature (e.g. single long branch that could either be attributed to HGT or gene loss in all other fungi, unstable phylogeny, and/or low bootstrap) of 1846 orthologues and so were subsequently removed. Of the remaining 291 orthologues, 8 had suspiciously high (>90%) first hit amino acid identity. Although the relatively recent divergence and/or acquisition time could explain this high level of similarity, we opted to remove these orthologues as a safeguard against possible bacterial contamination of the transcriptomes. Ultimately, a total of 283 distinct HGT events that satisfied the criteria described above for HGT were identified (Table S3). The average number of events per genus was 223±13 and ranged between 210 in the genus *Orpinomyces* to 242 in the genus *Pecoramyces* pantranscriptomes (Fig. 2A). The majority of HGT acquisition events identified (259, 91.52%) appear to be Neocallimastigomycota-specific, i.e. identified only in genomes belonging to the Neocallimastigomycota, but not in other basal fungal genomes (Table S4), strongly suggesting that such acquisitions occurred post, or concurrent with, the evolution of Neocallimastigomycota as a distinct fungal lineage. As well, the majority of these identified genes were Neocallimastigomycota-wide, being identified in strains belonging to at least six out of the seven examined genera (200 events, 70.7%), suggesting the acquisition of such genes prior to genus level diversification within the Neocallimastigomycota. Only 33 events (11.7%) were genus-specific, with the remainder (50 events, 17.7%) being identified in the transcriptomes of 3-5 genera (Table S4, Figure S4, and Fig. 2b).

**Figure 2.**
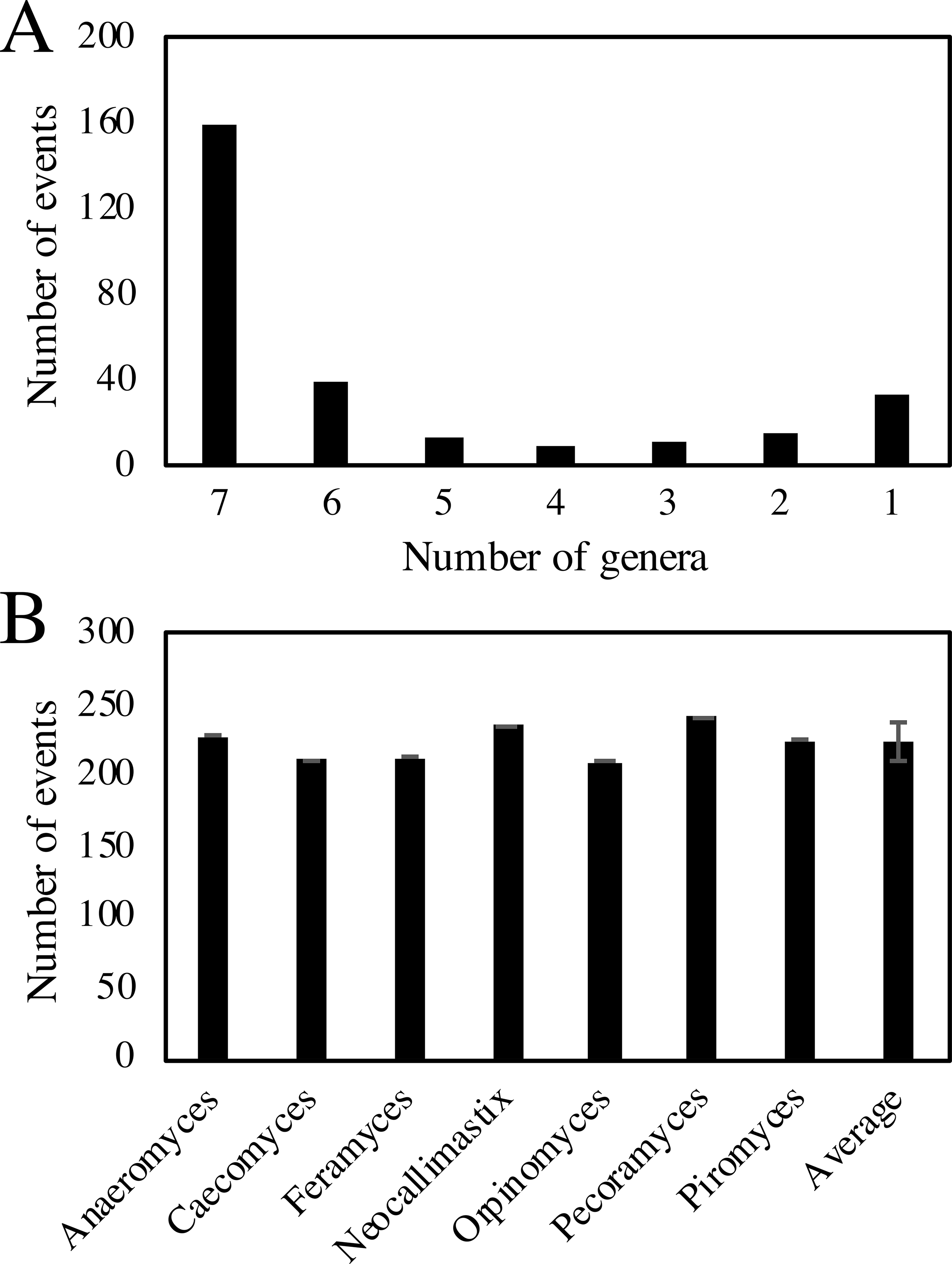
(A) Distribution pattern of HGT events in AGF transcriptomes demonstrating that the majority of events were Neocallimastigomycota-wide i.e. identified in all seven AGF genera examined. (B) Total Number of HGT events identified per AGF genus.

The absolute majority (89%) of events were successfully mapped to at least one of the four AGF genomes (Table S5), with a fraction (7/31) of the unmapped transcripts being specific to a genus with no genome representative (*Feramyces*, *Caecomyces*). Compared to a random subset of 283 genes in each of the sequenced genomes, horizontally transferred genes in AGF genomes exhibited significantly (P<0.0001) fewer introns (1.1±031 vs 3.32±0.83), as well as higher GC content (31±4.5 vs 27.7±5.5) (Table S5). Further, HGT genes/pfams often displayed high levels of gene/ pfam duplication and expansion within the genome (Table S5), resulting in an HGT frequency of 2.13% in *Pecoramyces ruminantium* (348 HGT genes out of 16,347 total genes), 3.07% in *Piromyces finnis* (352 HGT genes out of 11,477 total genes), 3.27% in *Anaeromyces robustus* (423 HGT genes out of 12,939 total genes), and 3.60% in *Neocallimastix californiae* (753 HGT genes out of 20,939 total genes).

### Donors

A bacterial origin was identified for the majority of HGT events (84.8%), with four bacterial phyla (Firmicutes, Proteobacteria, Bacteroidetes, and Spirochaetes) identified as donors for 177 events (62.5% of total, 73.8% of bacterial events) (Fig. 3A). Specifically, the contribution of members of the Firmicutes (125 events) was paramount, the majority of which were most closely affiliated with members of the order Clostridiales (106 events). In addition, minor contributions from a wide range of bacterial phyla were also identified (Fig. 3A). The majority of the putative donor taxa are strict/ facultative anaerobes, and many of which are also known to be major inhabitants of the herbivorous gut and often possess polysaccharide-degradation capabilities (83, 84). Archaeal contributions to HGT were extremely rare (6 events). On the other hand, multiple (34) events with eukaryotic donors were identified. In few instances, a clear non-fungal origin was identified for a specific event, but the precise inference of the donor based on phylogenetic analysis was not feasible (Table S4).

**Figure 3.**
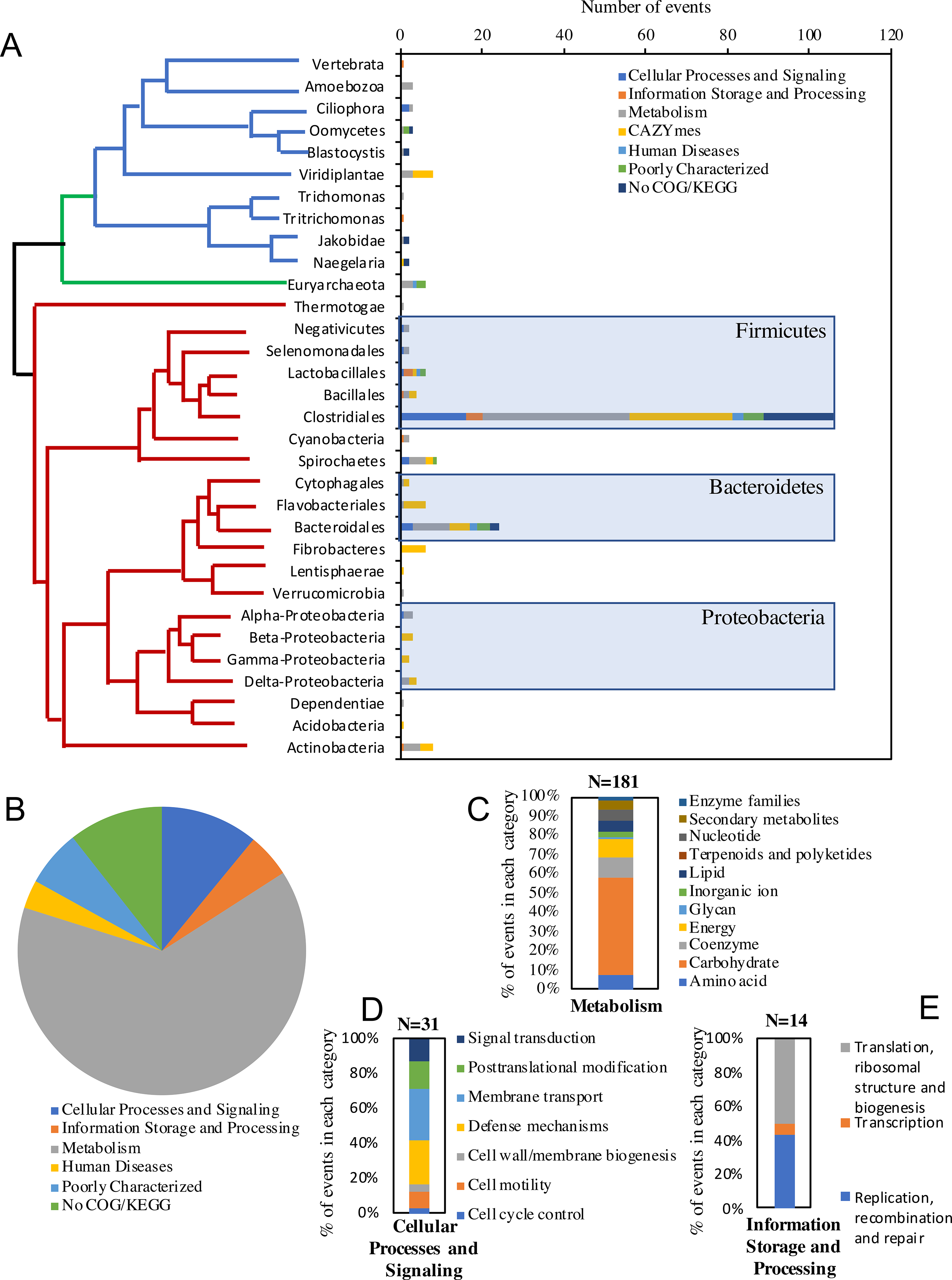
Identity of HGT donors and their contribution to the various functional classes. The X-axis shows the absolute number of events belonging to each of the functional classes shown in the legend. The tree is intended to show the relationship between the donors’ taxa and is not drawn to scale. Bacterial donors are shown with red branches depicting the phylum-level, with the exception of Firmicutes and Bacteroidetes donors, where the order-level is shown, and Proteobacteria, where the class-level is shown. Archaeal donors are shown with green branches and all belonged to the Methanobacteriales order of Euryarchaeota. Eukaryotic donors are shown with blue branches. Only the 230 events from a definitive-taxon donor are shown in the figure. The other 53 events were clearly nested within a non-fungal clade, but a definitive donor taxon could not be ascertained. Functional classification of the HGT events, determined by searching the Conserved Domain server (106) against the COG database are shown in B. For events with no COG classification, a search against the KEGG orthology database (107) was performed. For the major COG/KEGG categories (metabolism, cellular processes and signaling, and Information storage and processing), sub-classifications are shown in C, D, and E, respectively.

### Metabolic characterization

Functional annotation of HGT genes/pfams indicated that the majority (63.96%) of events encode metabolic functions such as extracellular polysaccharide degradation and central metabolic processes. Bacterial donors were slightly overrepresented in metabolic HGT events (87.3% of the metabolism-related events, compared to 84.8% of the total events). Genes involved in cellular processes and signaling represent the second most represented HGT events (10.95%), while genes involved in information storage and processing only made up 4.95% of the HGT events identified (Figs 3b-e). Below we present a detailed description of the putative abilities and functions enabled by HGT transfer events.

#### Central catabolic abilities

Multiple HGT events encoding various central catabolic processes were identified in AGF transcriptomes and successfully mapped to the genomes (Fig. 4, Table S4, Figs S5-S16). A group of events appear to encode enzymes that allow AGF to channel specific substrates into central metabolic pathways. For example, genes encoding enzymes of the Leloir pathway for galactose conversion to glucose-1-phosphate (galactose-1-epimerase, galactokinase (Fig. 5A), and galactose-1-phosphate uridylyltransferase) were identified, in addition to genes encoding ribokinase, as well as xylose isomerase and xylulokinase for ribose and xylose channeling into the pentose phosphate pathway. As well, genes encoding deoxyribose-phosphate aldolase (DeoC) enabling the utilization of purines as carbon and energy sources were also horizontally acquired in AGF. Further, several of the glycolysis/gluconeogenesis genes, e.g. phosphoenolpyruvate synthase, as well as phosphoglycerate mutase were also of bacterial origin. Fungal homologues of these glycolysis/gluconeogenesis genes were not identified in the AGF transcriptomes, suggesting the occurrence of xenologous replacement HGT events.

**Figure 4.**
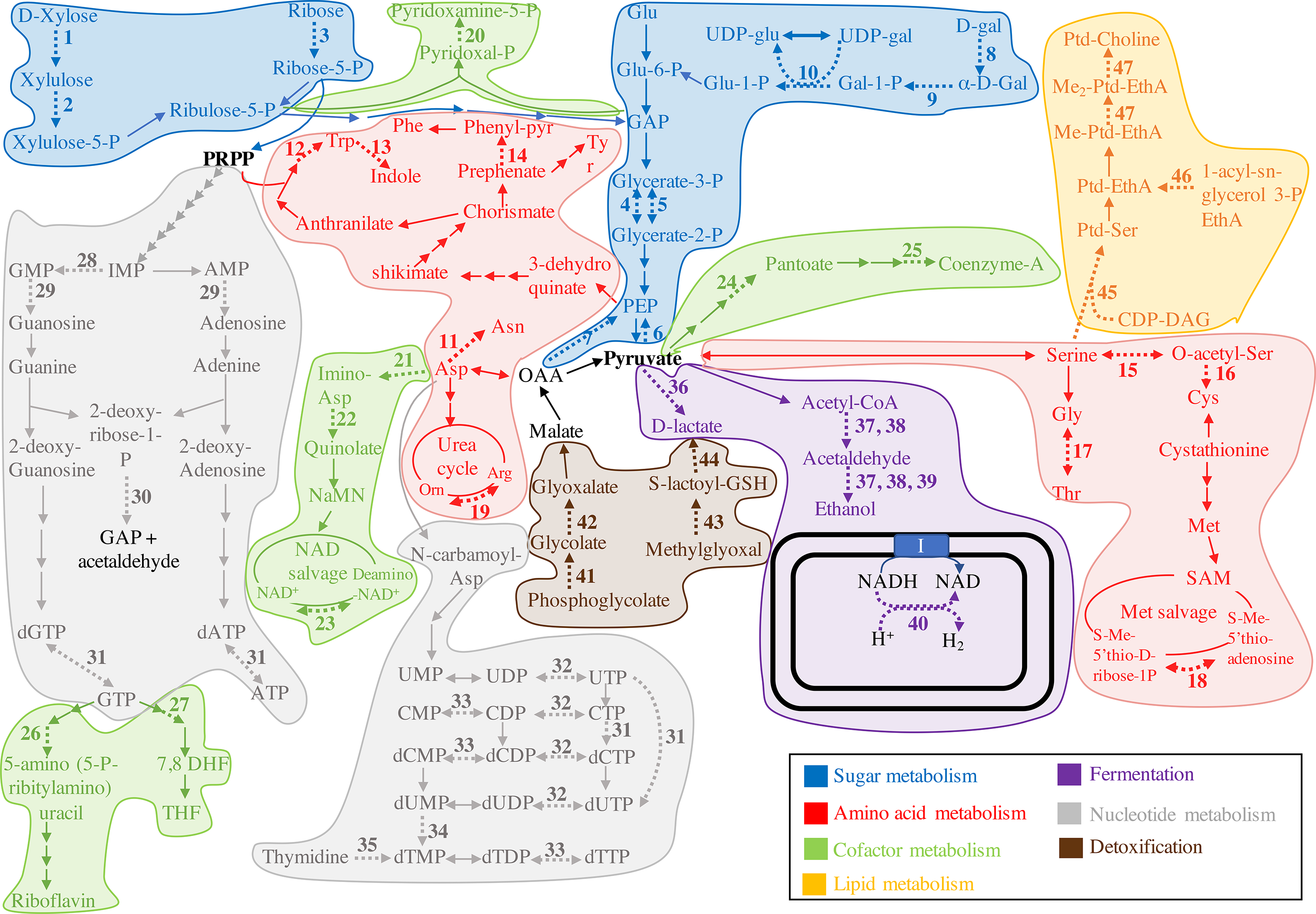
HGT impact on AGF central metabolic abilities. Pathways for sugar metabolism are highlighted in blue, pathways for amino acid metabolism are highlighted in red, pathways for cofactor metabolism are highlighted in green, pathways for nucleotide metabolism are highlighted in grey, pathways for lipid metabolism are highlighted in orange, fermentation pathways are highlighted in purple, while pathways for detoxification are highlighted in brown. The double black lines depict the hydrogenosomal outer and inner membrane. Arrows corresponding to enzymes encoded by horizontally transferred transcripts are shown with thicker dotted lines and are given numbers 1 through 48 as follows. Sugar metabolism (1-11): 1. Xylose isomerase, 2. Xylulokinase, 3. Ribokinase, 4. 2,3-bisphosphoglycerate-independent phosphoglycerate mutase, 5. 2,3-bisphosphoglycerate-dependent phosphoglycerate mutase, 6. Phosphoenolpyruvate synthase, 7. Phosphoenolpyruvate carboxykinase (GTP), 8. Aldose-1-epimerase, 9. Galactokinase, 10. Galactose-1-phosphate uridyltransferase. Amino acid metabolism (11-19): 11. Aspartate-ammonia ligase, 12. Tryptophan synthase (TrpB), 13. Tryptophanase, 14. Monofunctional prephenate dehydratase, 15. Serine-O-acetyltransferase, 16. Cysteine synthase, 17. Low-specificity threonine aldolase, 18. 5′-methylthioadenosine nucleosidase/5′-methylthioadenosine phosphorylase (MTA phosphorylase), 19. Arginase. Cofactor metabolism (20-27): 20. Pyridoxamine 5′-phosphate oxidase, 21. L-aspartate oxidase (NadB), 22. Quinolate synthase (NadA), 23. NH(3)-dependent NAD(+) synthetase (NadE), 24. 2-dehydropantoate 2-reductase, 25. dephosphoCoA kinase, 26. Dihydrofolate reductase (DHFR) family, 27. Dihydropteroate synthase. Nucleotide metabolism (28-35): 28. GMP reductase, 29. Trifunctional nucleotide phosphoesterase, 30. deoxyribose-phosphate aldolase (DeoC), 31. Oxygen-sensitive ribonucleoside-triphosphate reductase class III (NrdD), 32. nucleoside/nucleotide kinase family protein, 33. Cytidylate kinase-like family, 34. thymidylate synthase, 35. thymidine kinase. Pyruvate metabolism (fermentation pathways) (36-40): 36. D-lactate dehydrogenase, 37. bifunctional aldehyde/alcohol dehydrogenase family of Fe-alcohol dehydrogenase, 38. Butanol dehydrogenase family of Fe-alcohol dehydrogenase, 39. Zn-type alcohol dehydrogenase, 40. Fe-only hydrogenase. Detoxification reactions (41-44): 41. Phosphoglycolate phosphatase, 42. Glyoxal reductase, 43. Glyoxalase I, 44. Glyoxalase II. Lipid metabolism (45-47): 45. CDP-diacylglycerol--serine O-phosphatidyltransferase, 46. lysophospholipid acyltransferase LPEAT, 47. methylene-fatty-acyl-phospholipid synthase. Abbreviations: CDP-DAG, CDP-diacylglycerol; 7,8 DHF, 7,8 dihydrofolate; EthA, ethanolamine; Gal, galactose; GAP, glyceraldehyde-3-P; Glu, glucose; GSH, glutathione; I, complex I NADH dehydrogenase; NaMN, Nicotinate D-ribonucleotide; Orn, ornithine; PEP, phosphoenol pyruvate; Phenyl-pyr, phenylpyruvate; PRPP, phosphoribosyl-pyrophosphate; Ptd, phosphatidyl; SAM; S-adenosylmethionine; THF, tetrahydrofolate.

**Figure 5.**
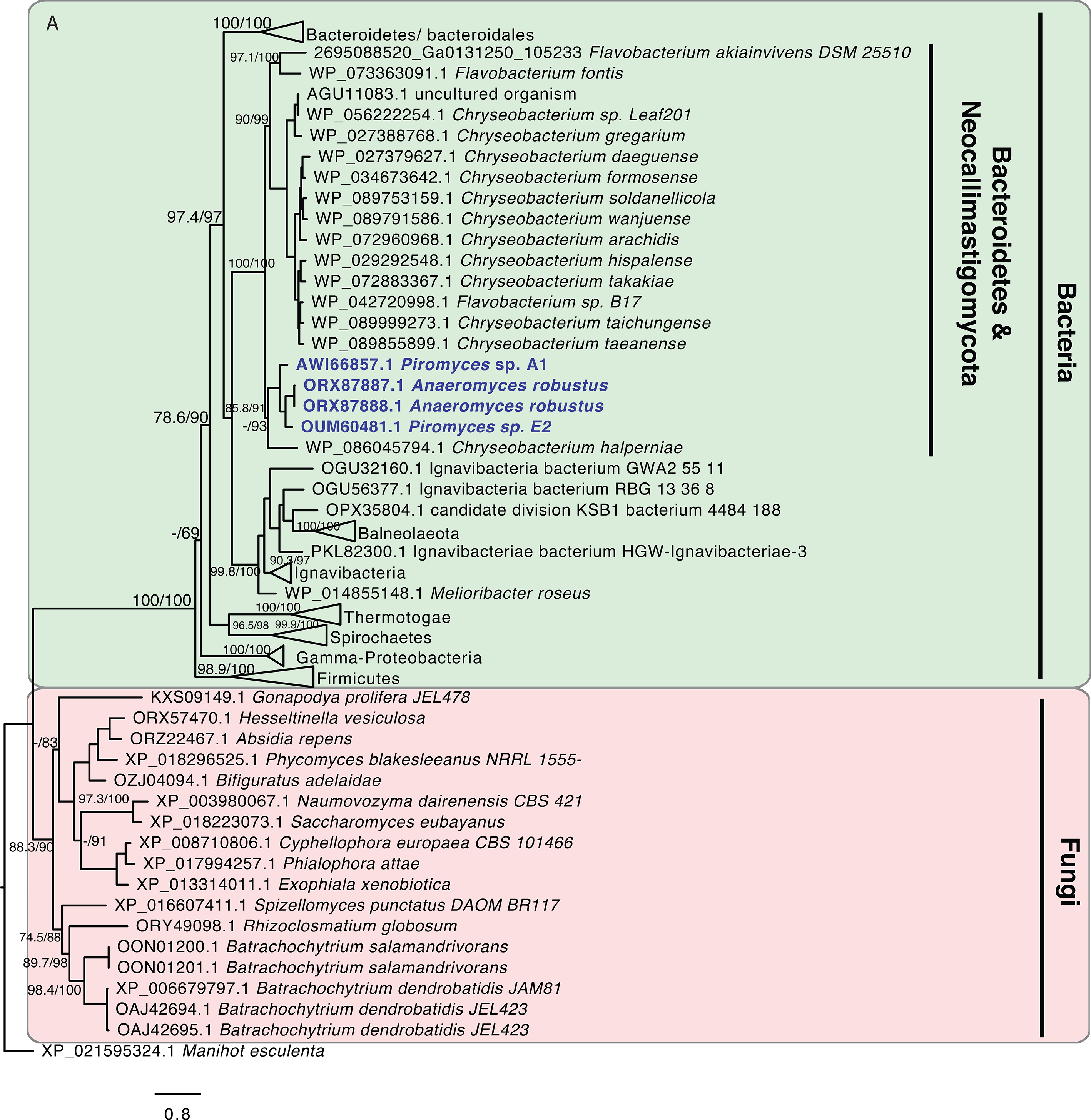
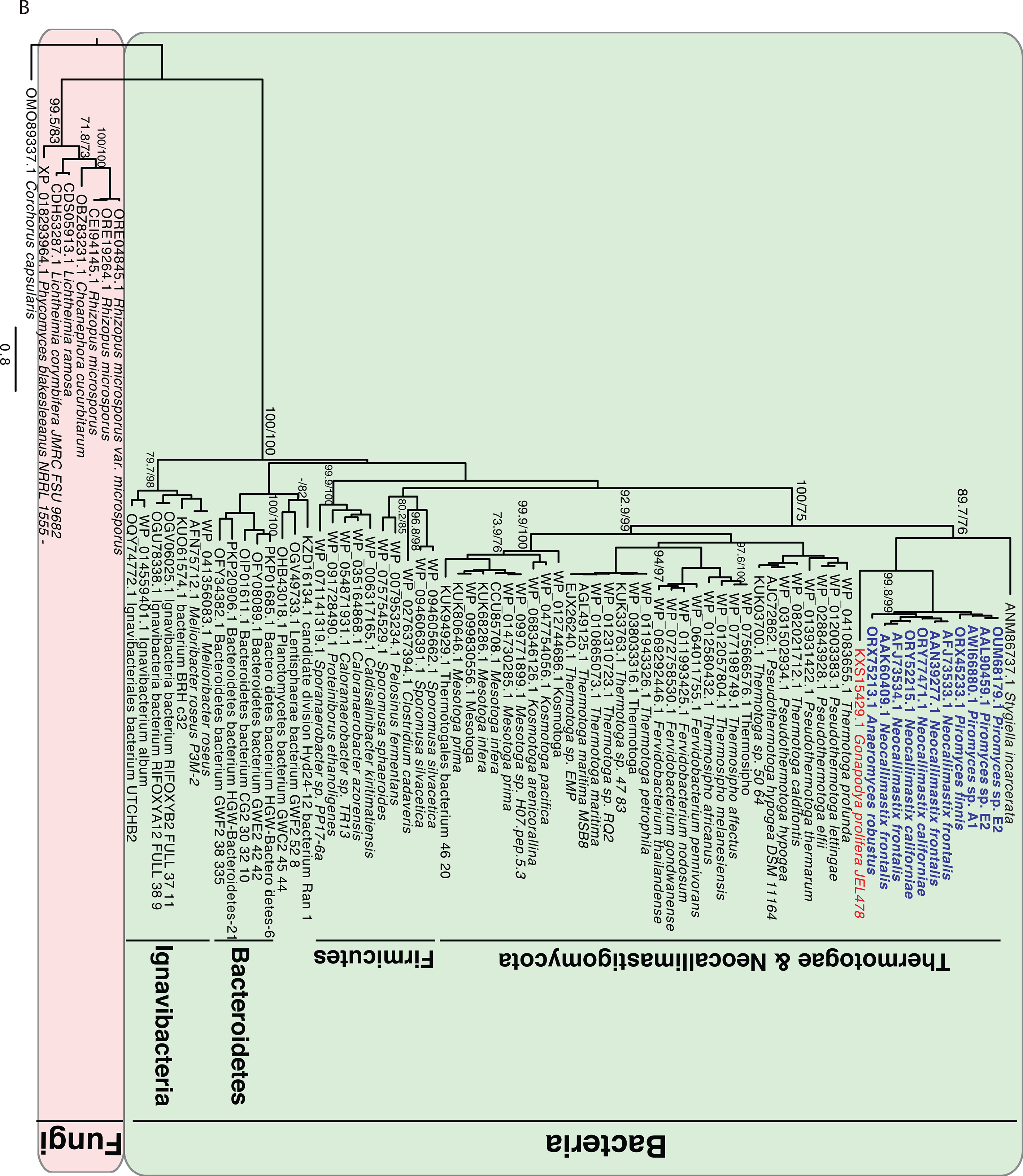
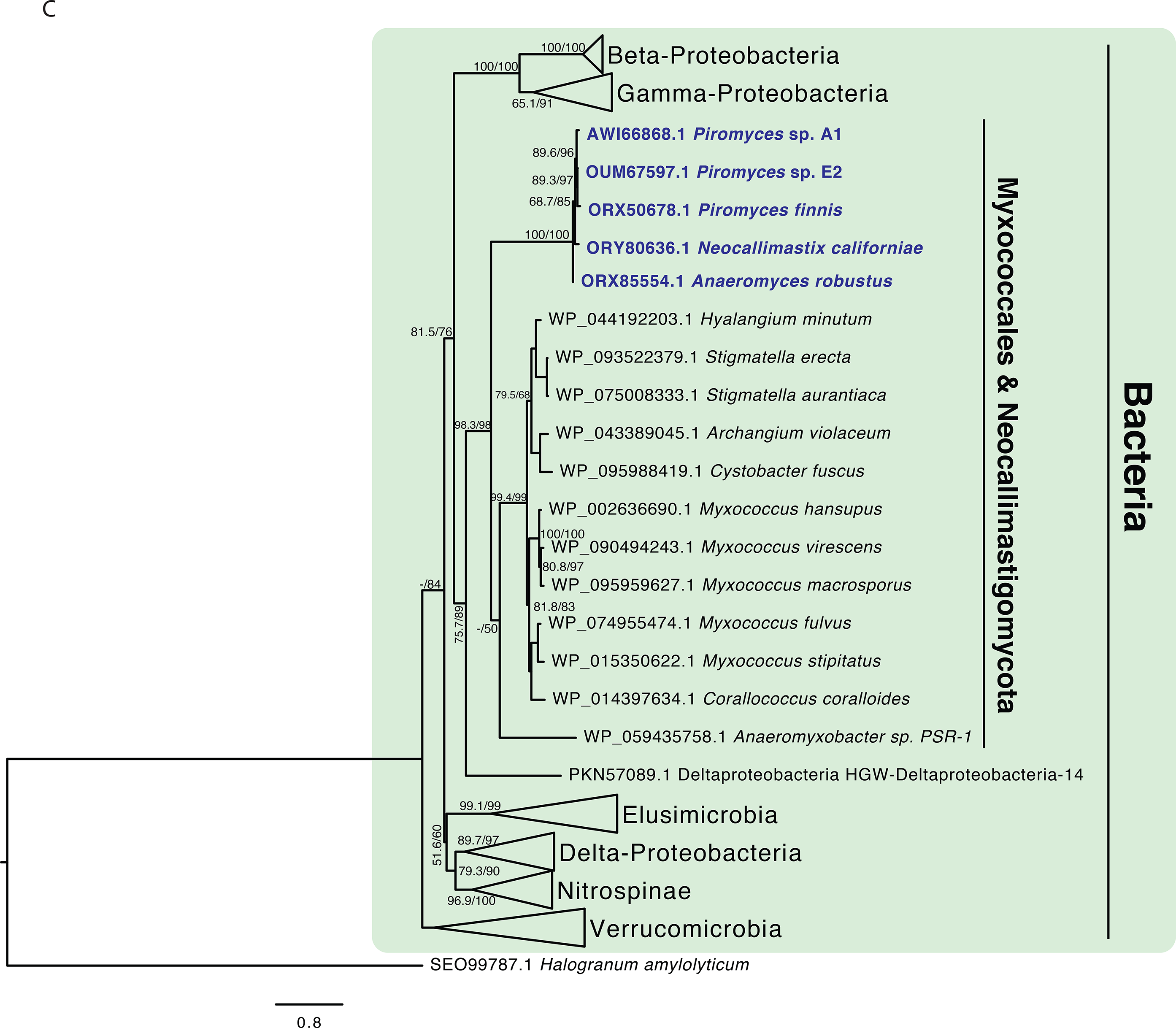
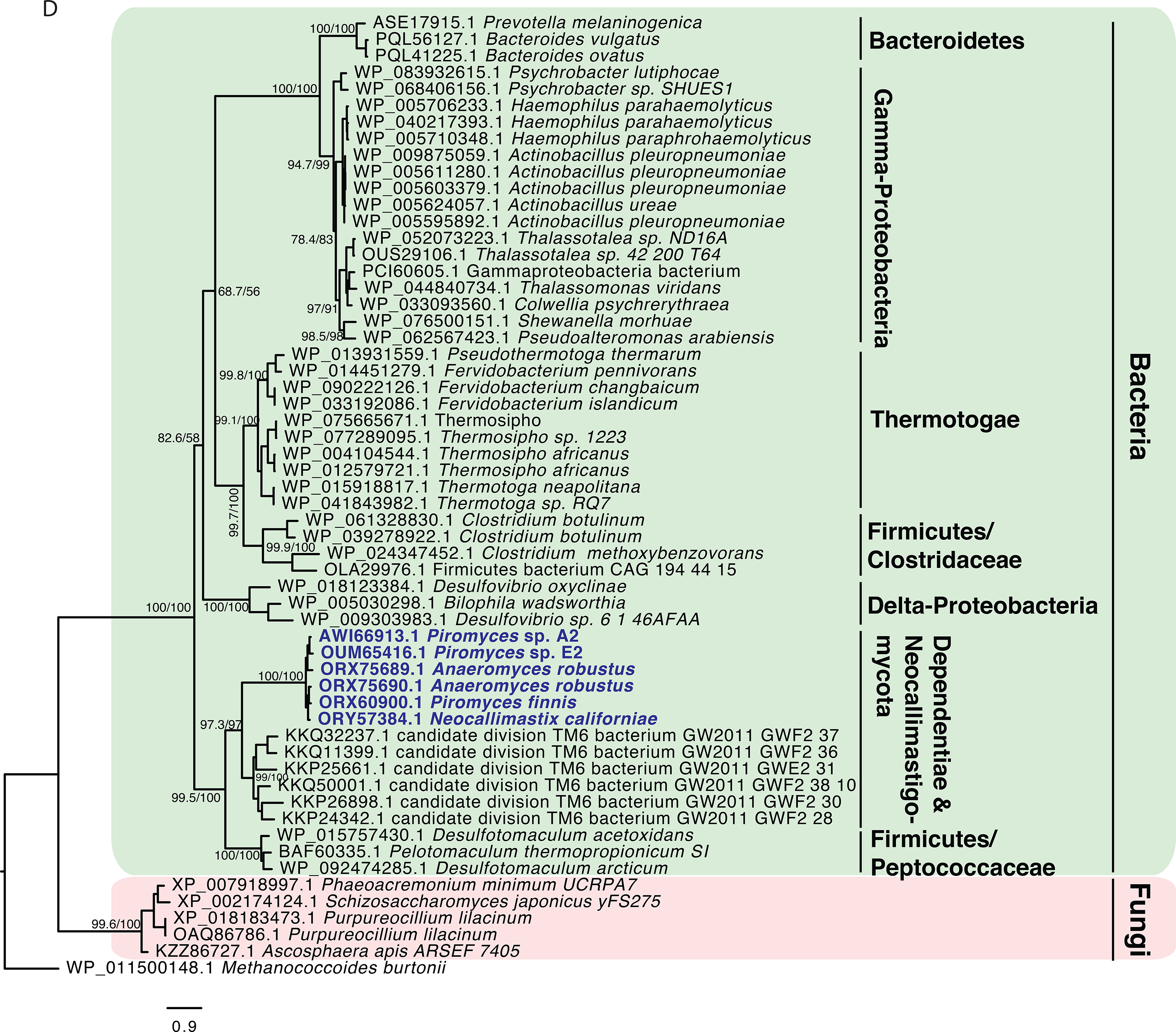
(A) Maximum likelihood tree showing the phylogenetic affiliation of AGF galactokinase. AGF genes highlighted in light blue clustered within the Flavobacteriales order of the Bacteroidetes phylum and were clearly nested within the bacterial domain (highlighted in green) attesting to their non-fungal origin. Fungal galactokinase representatives are highlighted in pink. (B) Maximum likelihood tree showing the phylogenetic affiliation of AGF Fe-only hydrogenase. AGF genes highlighted in light blue clustered within the Thermotogae phylum and were clearly nested within the bacterial domain (highlighted in green) attesting to their non-fungal origin. *Stygiella incarcerata* (anaerobic Jakobidae) clustered with the Thermotogae as well, as has recently been suggested (85). Fe-only hydrogenases from *Gonopodya prolifera* (Chytridiomycota) (shown in orange text) clustered with the AGF genes. This is an example of one of the rare occasions (n=24) where a non-AGF basal fungal representative showed an HGT pattern with the same donor affiliation as the Neocallimastigomycota. Other basal fungal Fe-only hydrogenase representatives are highlighted in pink and clustered outside the bacterial domain. (C) Maximum likelihood tree showing the phylogenetic affiliation of AGF L-aspartate oxidase (NadB). AGF genes highlighted in light blue clustered within the Delta-Proteobacteria class and were clearly nested within the bacterial domain (highlighted in green) attesting to their non-fungal origin. As de-novo NAD synthesis in fungi usually follow the five-enzyme pathway starting from tryptophan, as opposed to the two-enzyme pathway from aspartate, no NadB were found in non-AGF fungi and hence no fungal cluster is shown in the tree. (D) Maximum likelihood tree showing the phylogenetic affiliation of AGF oxygen-sensitive ribonucleotide reductase (NrdD). AGF genes highlighted in light blue clustered with representatives from Candidate phylum Dependentiae and were clearly nested within the bacterial domain (highlighted in green) attesting to their non-fungal origin. Fungal NrdD representatives are highlighted in pink. GenBank accession numbers are shown in parentheses. Alignment was done using the standalone Clustal Omega (66) and trees were constructed using IQ-tree (69).

In addition to broadening substrate range, HGT acquisitions provided additional venues for recycling reduced electron carriers via new fermentative pathways in this strictly anaerobic and fermentative lineage. The production of ethanol, D-lactate, and hydrogen appears to be enabled by HGT (Fig. 4). The acquisition of several aldehyde/alcohol dehydrogenases, and of D-Lactate dehydrogenase for ethanol and lactate production from pyruvate was identified. Although these two enzymes are encoded in other fungi as part of their fermentative capacity (e.g. *Saccharomyces* and *Schizosaccharomyces*), no homologues of these fungal genes were identified in AGF pantranscriptomes. Hydrogen production in AGF, as well as in many anaerobic eukaryotes with mitochondria-related organelles, involves pyruvate decarboxylation to acetyl CoA, followed by the use of electrons generated for hydrogen formation via an anaerobic Fe-Fe hydrogenase. In AGF, while pyruvate decarboxylation to acetyl CoA via pyruvate-formate lyase and the subsequent production of acetate via acetyl-CoA:succinyl transferase appear to be of fungal origin, the Fe-Fe hydrogenase and its entire maturation machinery (HydEFG) seem to be horizontally transferred being phylogenetically affiliated with similar enzymes in Thermotogae, Clostridiales, and the anaerobic jakobid excavate, *Stygiella incarcerate* (Fig. 5B). It has recently been suggested that *Stygiella* acquired the Fe-Fe hydrogenase and its maturation machinery from bacterial donors including Thermotogae, Firmicutes, and Spirochaetes (85), suggesting either a single early acquisition event in eukaryotes, or alternatively independent events for the same group of gene have occurred in different eukaryotes.

#### Anabolic capabilities

Multiple anabolic genes that expanded AGF biosynthetic capacities appear to be horizontally transferred (Fig. S17-S30). These include several amino acid biosynthesis genes e.g. cysteine biosynthesis from serine; glycine and threonine interconversion; and asparagine synthesis from aspartate. In addition, horizontal gene transfer allowed AGF to de-novo synthesize NAD via the bacterial pathway (starting from aspartate via L-aspartate oxidase (NadB; Fig. 5C) and quinolinate synthase (NadA) rather than the five-enzymes fungal pathway starting from tryptophan (86)). HGT also allowed AGF to salvage thiamine via the acquisition of phosphomethylpyrimidine kinase. Additionally, several genes encoding enzymes in purine and pyrimidine biosynthesis were horizontally transferred (Fig. 4). Finally, horizontal gene transfer allowed AGF to synthesize phosphatidyl-serine from CDP-diacylglycerol, and to convert phosphatidyl-ethanolamine to phosphatidyl-choline.

#### Adaptation to the host environment

Horizontal gene transfer also appears to have provided means of guarding against toxic levels of compounds known to occur in the host animal gut (Fig. S31-S37). For example, methylglyoxal, a reactive electrophilic species (87), is inevitably produced by ruminal bacteria from dihydroxyacetone phosphate when experiencing growth conditions with excess sugar and limiting nitrogen (88). Genes encoding enzymes mediating methylglyoxal conversion to D-lactate (glyoxalase I and glyoxalase II-encoding genes) appear to be acquired via HGT in AGF. Further, HGT allowed several means of adaptation to anaerobiosis. These include: 1) acquisition of the oxygen-sensitive ribonucleoside-triphosphate reductase class III (Fig. 5D) that is known to only function during anaerobiosis to convert ribonucleotides to deoxyribonucleotides (89), 2) acquisition of squalene-hopene cyclase, which catalyzes the cyclization of squalene into hopene, an essential step in biosynthesis of the cell membrane steroid tetrahymanol that replaced the molecular O_2_-requiring ergosterol in the cell membranes of AGF, 3) acquisition of several enzymes in the oxidative stress machinery including Fe/Mn superoxide dismutase, glutathione peroxidase, rubredoxin/rubrerythrin, and alkylhydroperoxidase.

In addition to anaerobiosis, multiple horizontally transferred general stress and repair enzymes were identified (Fig. S38-S45). HGT-acquired genes encoding 2-phosphoglycolate phosphatase, known to metabolize the 2-phosphoglycolate produced in the repair of DNA lesions induced by oxidative stress (90) to glycolate, were identified in all AGF transcriptomes studied (Fig. 4, Table S4). Surprisingly, two genes encoding antibiotic resistance enzymes, chloramphenicol acetyltransferase and aminoglycoside phosphotransferase, were identified in all AGF transcriptomes, presumably to improve its fitness in the eutrophic rumen habitat that harbors antibiotic-producing prokaryotes (Table S4). While unusual for eukaryotes to express antibiotic resistance genes, basal fungi such as *Allomyces, Batrachochytrium,* and *Blastocladiella* were shown to be susceptible to chloramphenicol and streptomycin (91, 92). Other horizontally transferred repair enzymes include DNA-3-methyladenine glycosylase I, methylated-DNA--protein-cysteine methyltransferase, galactoside and maltose O-acetyltransferase, and methionine-R-sulfoxide reductase (Table S4).

#### HGT transfer in AGF carbohydrate active enzymes machinery

Within the analyzed AGF transcriptomes, CAZymes belonging to 39 glycoside hydrolase (GHs), 5 polysaccharide lyase (PLs), and 10 carbohydrate esterase (CEs) families were identified (Fig. 6). The composition of the CAZymes of various AGF strains examined were broadly similar, with the following ten notable exceptions: Presence of GH24 and GH78 transcripts only in *Anaeromyces* and *Orpinomyces,* the presence of GH28 transcripts only in *Pecoramyces*, *Neocallimastix*, and *Orpinomyces*, the presence of GH30 transcripts only in *Anaeromyces*, and *Neocallimastix*, the presence of GH36 and GH95 transcripts only in *Anaeromyces*, *Neocallimastix*, and *Orpinomyces,* the presence of GH97 transcripts only in *Neocallimastix*, and *Feramyces*, the presence of GH108 transcripts only in *Neocallimastix*, and *Piromyces*, and the presence of GH37 predominantly in *Neocallimastix*, GH57 transcripts predominantly in *Orpinomyces*, GH76 transcripts predominantly in *Feramyces*, and CE7 transcripts predominantly in *Anaeromyces* (Fig. 6).

**Figure 6.**
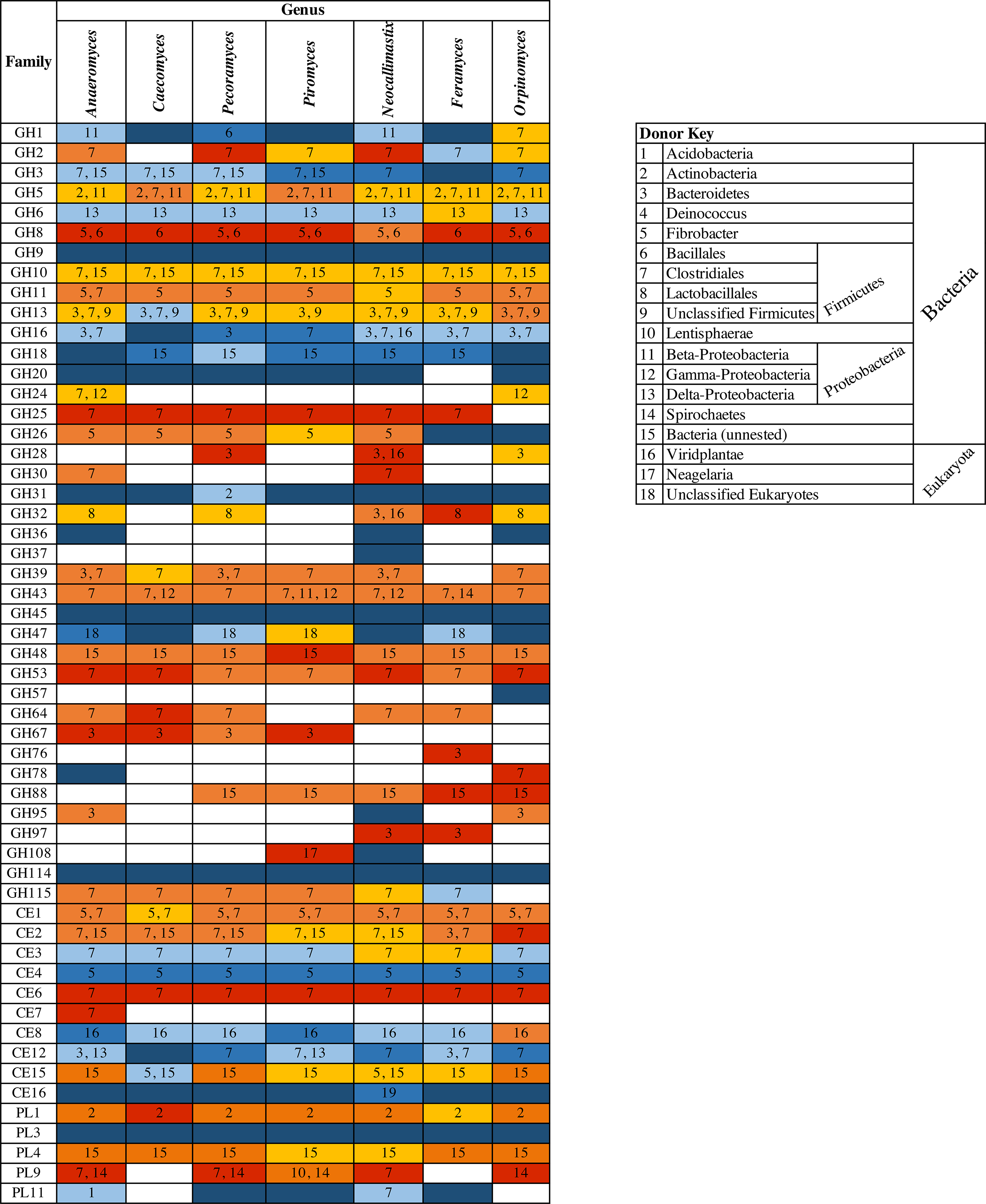
HGT in the AGF CAZyome shown across the seven genera studied. Glycosyl Hydrolase (GH), Carboxyl Esterase (CE), and Polysaccharide Lyase (PL) families are shown to the left. The color of the cells depicts the prevalence of HGT within each family. Red indicates that 100% of the CAZyme transcripts were horizontally transferred. Shades of red-orange indicate that HGT contributed to > 50% of the transcripts belonging to that CAZy family. Blue indicates that 100% of the CAZyme transcripts were of fungal origin. Shades of blue indicate that HGT contributed to < 50% of the transcripts belonging to that CAZy family. The numbers in each cell indicate the affiliation of the HGT donor as shown in the key to the right.

HGT appears to be rampant in the AGF pan-CAZyome: A total of 75 events (26.5% of total HGT events) were identified, with 40% occurring in all AGF genera examined (Fig. 6, Table S4). In 36.5% of GH families, 37.5% of CE families, and 28.6% of PL families, a single event (i.e. attributed to one donor) was observed (Fig. 6, Table S4).

Duplication of these events in AGF genomes was notable, with 134, 322, 161, and 135 copies of HGT CAZyme pfams identified in *Anaeromyces*, *Neocallimastix*, *Piromyces* and *Pecoramyces* genomes, representing 34.1%, 38.2%, 41.7%, and 25.6% of the overall organismal CAZyme machinery (Table S5). The contribution of Viridiplantae, Fibrobacteres, and Gamma-Proteobacteria was either exclusive to CAZyme-related HGT events or significantly higher in CAZyme, compared to other, events (Fig. 3A).

Transcripts acquired by HGT represented >50% of transcripts in anywhere between 13 (*Caecomyces*) to 20 (*Anaeromyces*) GH families; 3 (*Caecomyces*) to 5 (*Anaeromyces*, *Neocallimastix*, *Orpinomyces*, and *Feramyces*) CE families; and 2 (*Caecomyces* and *Feramyces*) to 3 (*Anaeromyces*, *Pecoramyces*, *Piromyces*, *Neocallimastix*, and *Orpinomyces*) PL families (Fig. 6). It is important to note that in all these families, multiple transcripts appeared to be of bacterial origin based on BLAST similarity search but did not meet the strict criteria implemented for HGT determination in this study. As such, the contribution of HGT transcripts to overall transcripts in these families is probably an underestimate. Only GH9, GH20, GH37, GH45, and PL3 families appear to lack any detectable HGT events. A PCA biplot comparing CAZyomes in AGF genomes to other basal fungal lineages strongly suggests that the acquisition and expansion of many of these foreign genes play an important role in shaping the lignocellulolytic machinery of AGF (Fig. 7). The majority of CAZyme families defining AGF CAZyome were predominantly of non-fungal origin (Fig. 7). This pattern clearly attests to the value of HGT in shaping AGF CAZyome via acquisition and extensive duplication of acquired gene families.

**Figure 7.**
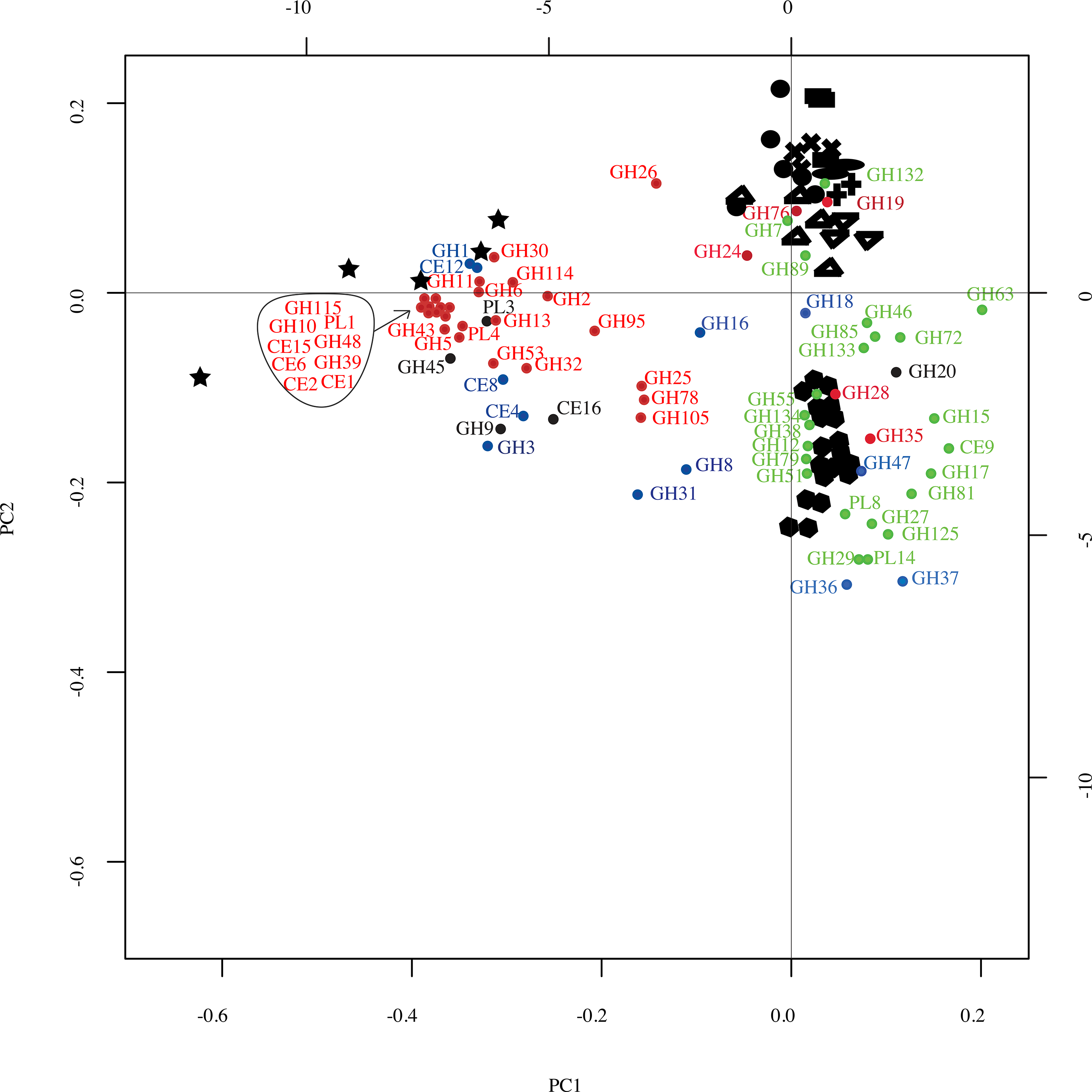
Principal-component analysis biplot of the distribution of CAZy families in AGF genomes (★), compared to representatives of other basal fungi belonging to the Mucoromycotina 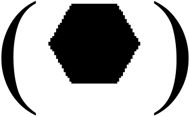, Chytridiomycota (●), Blastocladiomycota (■), Entomophthoromycotina 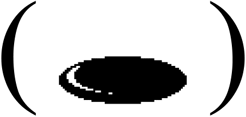, Mortierellomycotina (Δ), Glomeromycota (✚), Kickxellomycotina (▽), and Zoopagomycotina (✖). CAZy families are shown as colored dots. The color code used was as follows: green, CAZy families that are absent from AGF genomes; black, CAZy families present in AGF genomes and with an entirely fungal origin; blue, CAZy families present in AGF genomes and for which HGT contributed to < 50% of the transcripts in the examined transcriptomes; red, CAZy families present in AGF genomes and for which HGT contributed to > 50% of the transcripts in the examined transcriptomes. The majority of CAZyme families defining the AGF CAZyome were predominantly of non-fungal origin (red and blue dots).

Collectively, HGT had a profound impact on AGF plant biomass degradation capabilities. The AGF CAZyome encodes enzymes putatively mediating the degradation of twelve different polysaccharides (Fig. S46). In all instances, GH and PL families with >50% horizontally transferred transcripts contributed to backbone cleavage of these polymers; although in many polymers, e.g. cellulose, glucoarabinoxylan, and rhamnogalactouronan, multiple different GHs can contribute to backbone cleavage. Similarly, GH, CE, and PL families with >50% horizontally transferred transcripts contributed to 10 out of 13 side-chain-cleaving activities, and 3 out of 5 oligomer-to-monomer breakdown activities (Fig. S46).

## Discussion

Here, we present a systematic analysis of HGT patterns in 27 transcriptomes and 4 genomes belonging to the Neocallimastigomycota. Our analysis identified 283 events, representing 2.1-3.6% of genes in examined AGF genomes. Further, we consider these values to be conservative estimates due to the highly stringent criteria and employed. Only events with *h*_*U*_ of >30 were considered, and all putative events were further subjected to manual inspection and phylogenetic tree construction to confirm incongruence with organismal evolution and bootstrap-supported affiliation to donor lineages. Further, events identified in less than 50% of strains in a specific genus were excluded, and parametric gene composition approaches were implemented in conjunction with sequence-based analysis.

Multiple factors could be postulated to account for the observed high HGT frequency in AGF. The sequestration of AGF into the anaerobic, prokaryotes-dominated herbivorous gut necessitated the implementation of the relatively faster adaptive mechanisms for survival in this new environment, as opposed to the slower strategies of neofunctionalization and gene birth. Indeed, niche adaptation and habitat diversification events are widely considered important drivers for HGT in eukaryotes (16, 23, 93, 94). Further, AGF are constantly exposed to a rich milieu of cells and degraded DNA in the herbivorous gut. Such close physical proximity between donors/ extracellular DNA and recipients is also known to greatly facilitate HGT (95-97). Finally, AGF release asexual motile free zoospores into the herbivorous gut as part of their life cycle (39). According to the weak-link model (98), these weakly protected and exposed structures provide excellent entry point of foreign DNA to eukaryotic genomes. It is important to note that AGF zoospores also appear to be naturally competent, capable of readily uptaking nucleic acids from their surrounding environment (50).

The distribution of HGT events across various AGF taxa (Fig. 2), identities of HGT donors (Fig. 3), and abilities imparted (Figs. 4–5) could offer important clues regarding the timing and impact of HGT on Neocallimastigomycota evolution. The majority of events (70.7%) were Neocallimastigomycota-wide and were mostly acquired from lineages known to inhabit the herbivorous gut, e.g. Firmicutes, Proteobacteria, Bacteroidetes, and Spirochaetes (Figs. 2–3). This pattern strongly suggests that such acquisitions occurred post (or concurrent with) AGF sequestration into the herbivorous gut, but prior to AGF genus level diversification. Many of the functions encoded by these events represented novel functional acquisitions that impart new abilities, e.g. galactose metabolism, methyl glyoxal detoxification, pyruvate fermentation to d-lactate and ethanol, and chloramphenicol resistance (Fig. 3). Others represented acquisition of novel genes or pfams augmenting existing capabilities within the AGF genomes, e.g. acquisition of GH5 cellulases to augment the fungal GH45, acquisition of additional GH1 and GH3 beta gluco- and galactosidases to augment similar enzymes of apparent fungal origin in AGF genomes (Fig. 6–7, Fig. S46). Novel functional acquisition events enabled AGF to survive and colonize the herbivorous gut by: 1. Expanding substrate-degradation capabilities (Fig. 5a, 6, 7, S5-S17, Table S4), hence improving fitness by maximizing carbon and energy acquisition from available plant substrates, 2. Providing additional venues for electron disposal via lactate, ethanol, and hydrogen production, and 3. Enabling adaptation to anaerobiosis (Fig. 4, S32-S38, Table S4).

A smaller number of observed events (n=33) were genus-specific (Fig. 2, Table S4). This group was characterized by being significantly enriched in CAZymes (60.6% of genus-specific horizontally transferred events have a predicted CAZyme function, as opposed to 26.5% in the overall HGT dataset), and being almost exclusively acquired from donors that are known to inhabit the herbivorous gut (99) (26 out of the 33 events were acquired from the orders Clostridiales, Bacillales, and Negativicutes within Firmicutes, Burkholderiales within the Beta-Proteobacteria, Flavobacteriales and Bacteroidales within Bacteroidetes, and the Spirochaetes, Actinobacteria, and Lentisphaerae), or from Viridiplantae (4 out of the 33 events). Such pattern suggests the occurrence of these events relatively recently, in the herbivorous gut post AGF genus level diversification. We reason that the lower frequency of such events is a reflection of the relaxed pressure for acquisition and retention of foreign genes at this stage of AGF evolution.

Gene acquisition by HGT necessitates physical contact between donor and recipient organisms. Many of the HGT acquired traits by AGF are acquired from prokaryotes that are prevalent in the herbivorous gut microbiota (Fig. 3). However, since many of these traits are absolutely necessary for survival in the gut, the establishment of AGF ancestors in this seemingly inhospitable habitat is, theoretically, unfeasible. This dilemma is common to all HGT processes enabling niche adaptation and habitat diversification (22). We put forth two evolutionary scenarios that could explain this dilemma not only for AGF, but also for other gut-dwelling anaerobic microeukaryotes, e.g. *Giardia, Blastocystis*, and *Entamoeba*, where HGT was shown to play a vital role in enabling survival in anaerobic conditions (100-102). The first is a coevolution scenario in which the progressive evolution of the mammalian gut from a short and predominantly aerobic structure characteristic of carnivores/insectivores to the longer, more complex, and compartmentalized structure encountered in herbivores was associated with a parallel progressive and stepwise acquisition of genes required for plant polymers metabolism and anaerobiosis by AGF ancestors, hence assuring its survival and establishment in the current herbivorous gut. The second possibility is that AGF ancestors were indeed acquired into a complex and anaerobic herbivorous gut, but initially represented an extremely minor component of the gut microbiome and survived in locations with relatively higher oxygen concentration in the alimentary tract e.g. mouth, saliva, esophagus or in micro-niches in the rumen where transient oxygen exposure occurs. Subsequently, HGT acquisition has enabled the expansion of their niche, improved their competitiveness and their relative abundance in the herbivorous gut to the current levels.

In conclusion, our survey of HGT in AGF acquisition demonstrates that the process is absolutely crucial for the survival and growth of AGF in its unique habitat. This is not only reflected in the large number of events, massive duplication of acquired genes, and overall high HGT frequency observed in AGF genomes, but also in the nature of abilities imparted by the process. HGT events not only facilitated AGF adaptation to anaerobiosis, but also allowed them to drastically improve their polysaccharide degradation capacities, provide new venues for electron disposal via fermentation, and acquire new biosynthetic abilities. As such, we reason that the process should not merely be regarded as a conduit for supplemental acquisition of few additional beneficial traits. Rather, we posit that HGT enabled AGF to forge a new evolutionary trajectory, resulting in Neocallimastigomycota sequestration, evolution as a distinct fungal lineage in the fungal tree of life, and subsequent genus and species level diversification. This provides an excellent example of the role of HGT in forging the formation of high rank taxonomic lineages during eukaryotic evolution.

## Supporting information

Entire supplementary document

## Acknowledgments

This work has been funded by the NSF-DEB Grant numbers 1557102 to N.Y. and M.E. and 1557110 to J.E.S.

