## Supplementary material for "Horizontal gene transfer as an indispensable driver for Neocallimastigomycota evolution into a distinct gut-dwelling fungal lineage": Entire supplementary document

**Table S1. Validation of HGT-identification pipeline using previously published datasets.** The frequency of HGT occurrence in the genomes of a filamentous ascomycete and a microsporidian were determined using our pipeline. The results were compared to previously published results.

| Organism | NCBI Assembly accession number | Reference to original study | Method used in the original study | Value reported | Value obtained in this study |
|---|---|---|---|---|---|
| *Colletotrichum graminicola* | GCA_000149035.1 | (1) | Blast and tree building approaches | 11 | 11 |
| *Encephalitozoon hellem* | GCA_000277815.3 | (2) | Blast against custom database, AI score calculation, and tree building | 12-22 | 4 |

**Table S2.** Results of transcriptomic sequencing.

| **Genus** | **Species** | **Strain** | **Number of reads** | **Assembled transcripts**[a] | **Predicted peptides (Longest Orfs)**[b] | **% completeness**[c] | **genome coverage**[d] (%) | Accession number | Ref. |
|---|---|---|---|---|---|---|---|---|---|
| *Anaeromyces* | *contortus* | C3G | 33,374,692 | 50,577 | 22,187 | 96.55 | | GGWR00000000 | This study |
| *Anaeromyces* | *contortus* | C3J | 54,320,879 | 57,658 | 26,052 | 97.24 | | GGWO00000000 | This study |
| *Anaeromyces* | *contortus* | G3G | 43,154,980 | 52,929 | 21,681 | 91.38 | | GGWP00000000 | This study |
| *Anaeromyces* | *contortus* | Na | 42,857,287 | 47,378 | 19,386 | 93.45 | | GGWN00000000 | This study |
| *Anaeromyces* | *contortus* | O2 | 60,442,723 | 62,300 | 27,322 | 96.9 | | GGWQ00000000 | This study |
| *Anaeromyces* | *robustus* | S4 | 21,955,935 | NA | 17,127 | 92.41 | 88.7 | SRX3329608 | (3) |
| *Caecomyces* | sp. | Iso3 | 21,766,139 | 88,894 | 42,308 | 96.55 | | GGXE00000000 | This study |
| *Caecomyces* | sp. | Brit4 | 15,199,296 | 59,747 | 24,950 | 92.76 | | GGWS00000000 | This study |
| *Feramyces* | *austinii* | WSF2 | 23,024,082 | 110,716 | 37,114 | 91.03 | | GGWT00000000 | This study |
| *Feramyces* | *austinii* | WSF3 | 25,911,634 | 155,190 | 42,823 | 91.38 | | GGWU00000000 | This study |
| *Neocallimastix* | *californiae* | G1 | 36,250,970 | NA | 29,649 | 95.52 | 85.2 | SRX2598479 | (3) |
| *Neocallimastix* | cf. *cameroonii* | G3 | 54,184,578 | 98,268 | 35,649 | 96.21 | | GGXC00000000 | This study |
| *Neocallimastix* | cf. *frontalis* | Hef5 | 22,033,159 | 95,833 | 47,305 | 96.21 | | GGXJ00000000 | This study |
| *Orpinomyces* | cf. *joyonii* | D3A | 27,863,051 | 36,927 | 17,648 | 91.38 | | GGWV00000000 | This study |

| | | | | | | | | | |
|---|---|---|---|---|---|---|---|---|---|
| *Orpinomyces* | cf. *joyonii* | D3B | 25,397,280 | 31,765 | 16,500 | 90.69 | | GGWW00000000 | This study |
| *Orpinomyces* | cf. *joyonii* | D4C | 25,949,247 | 31,155 | 16,756 | 93.45 | | GGWX00000000 | This study |
| *Pecoramyces* | *ruminantium* | C1A | 468,159,494 | 35,126 | 27,506 | 97.59 | 85.7 | SRX1030108 | (4) |
| *Pecoramyces* | *ruminantium* | S4B | 46,78,2033 | 68,418 | 28,731 | 96.21 | | GGWY00000000 | This study |
| *Pecoramyces* | *ruminantium* | FS3C | 23,575,308 | 97,381 | 41,329 | 93.45 | | GGXF00000000 | This study |
| *Pecoramyces* | *ruminantium* | FX4B | 25,011,052 | 66,074 | 27,589 | 92.07 | | GGWZ00000000 | This study |
| *Pecoramyces* | *ruminantium* | YC3 | 23,080,103 | 58,071 | 27,392 | 91.72 | | GGXA00000000 | This study |
| *Piromyces* | *finnis* | finn | 25,770,853 | NA | 17,008 | 95.17 | 91.4 | SRX2770525 | (3) |
| *Piromyces* | sp. | A1 | 39,201,276 | 50,514 | 22,628 | 85.52 | | GGXB00000000 | This study |
| *Piromyces* | sp. | A2 | 50,945,955 | 55,306 | 30,581 | 94.83 | | GGXG00000000 | This study |
| *Piromyces* | sp. | B4 | 104,293,067 | 140,717 | 70,061 | 93.1 | | GGXH00000000 | This study |
| *Piromyces* | sp. | B5 | 110,842,283 | 187,158 | 49,460 | 82.76 | | GGXI00000000 | This study |
| *Piromyces* | sp. | E2 | NA | NA | 17,080 | 81.72 | 70.9 | | (3) |

a: Trinity (v 2.5.0) was used for the read assembly

b: Predicted using Transdecoder using transcript orthologues (95%) clustered using CD-HIT-EST
c: Calculated using BUSCO (V2)
d: Percentage of genes in the strain's genome for which a transcript was identified. Numbers were obtained from the individual genome webpages in IMG portal

Tables S3-S5 are provided as separate Excel sheets.

**Table S3.** The identified 283 HGT events. HGT events were identified using a combination of non-fungal Blastp bit score cutoff (>100), HGT index cutoff (>30), and downstream phylogenetic analysis.

**Table S4.** HGT events identified with the affiliation of the donor, distribution across AGF genera studied, and specificity of gene acquisition to the Neocallimastigomycota phylum.

**Table S5.** Mapping HGT events to available fungal genomes (*An. rob, Anaeromyces robustus; Ne. cal, Neocallimastix californiae; Pi. fin, Piromyces finnis; Pe. rum, Pecoramyces ruminantium*), and general characteristics of HGT genes identified.

**Supplementary Figures.**

**Figure S1. Cartoon depicting the evolutionary history and the life cycle of the AGF in the herbivorous gut.** Ancestors of the AGF have been introduced to ancient herbivores through a yet unknown mechanism, possibly ingestion. We hypothesize that the sequestration of AGF into the herbivorous gut was conducive to trait acquisition by HGT as a relatively faster strategy for niche adaptation. We further hypothesize that the donors were mostly bacterial residents of the herbivorous gut, since the life cycle of the AGF occurs entirely in the herbivorous gut, providing little opportunity for HGT acquisition from other environments. The lifecycle of AGF involves the migration of AGF zoospores towards ingested plant material (step 1), followed by the attachment and the encystment of the zoospore (Step 2). The cyst germinates (step 3), resulting in rhizoid development (Step 4) and sporangium maturation (step 5). Zoospores differentiate in the sporangia and are subsequently released to complete the life cycle (step 6).

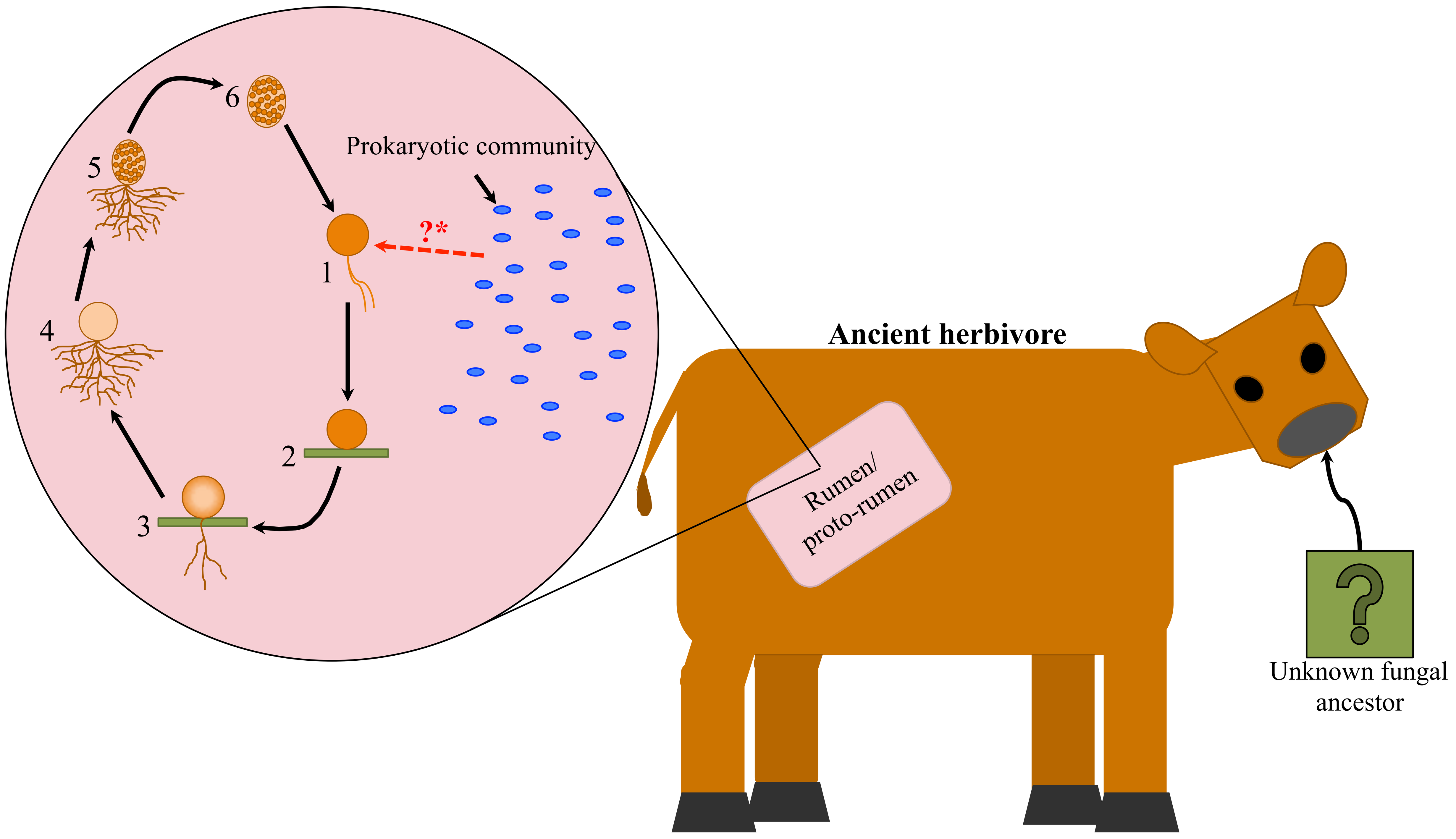

**Figure S2. Multi-domain CAZymes.** The modular nature of many CAZyme containing genes in AGF necessitates the utilization of a pfam-based rather than entire gene/transcriptome-based strategy for HGT detection. In AGF genomes, CAZyme domains of apparent different origins (fungal or non-fungal origins) are often encountered, leading to inaccurate HGT assessments when using the entire gene for similarity searches. Examples from a single *(P. finnis)* AGF genome are provided below. GenBank protein accession number are shown above each gene. Different domains are shown as boxes with different colors and the identity of the first hit are shown within the boxes. JGI protein ID are shown to the left.

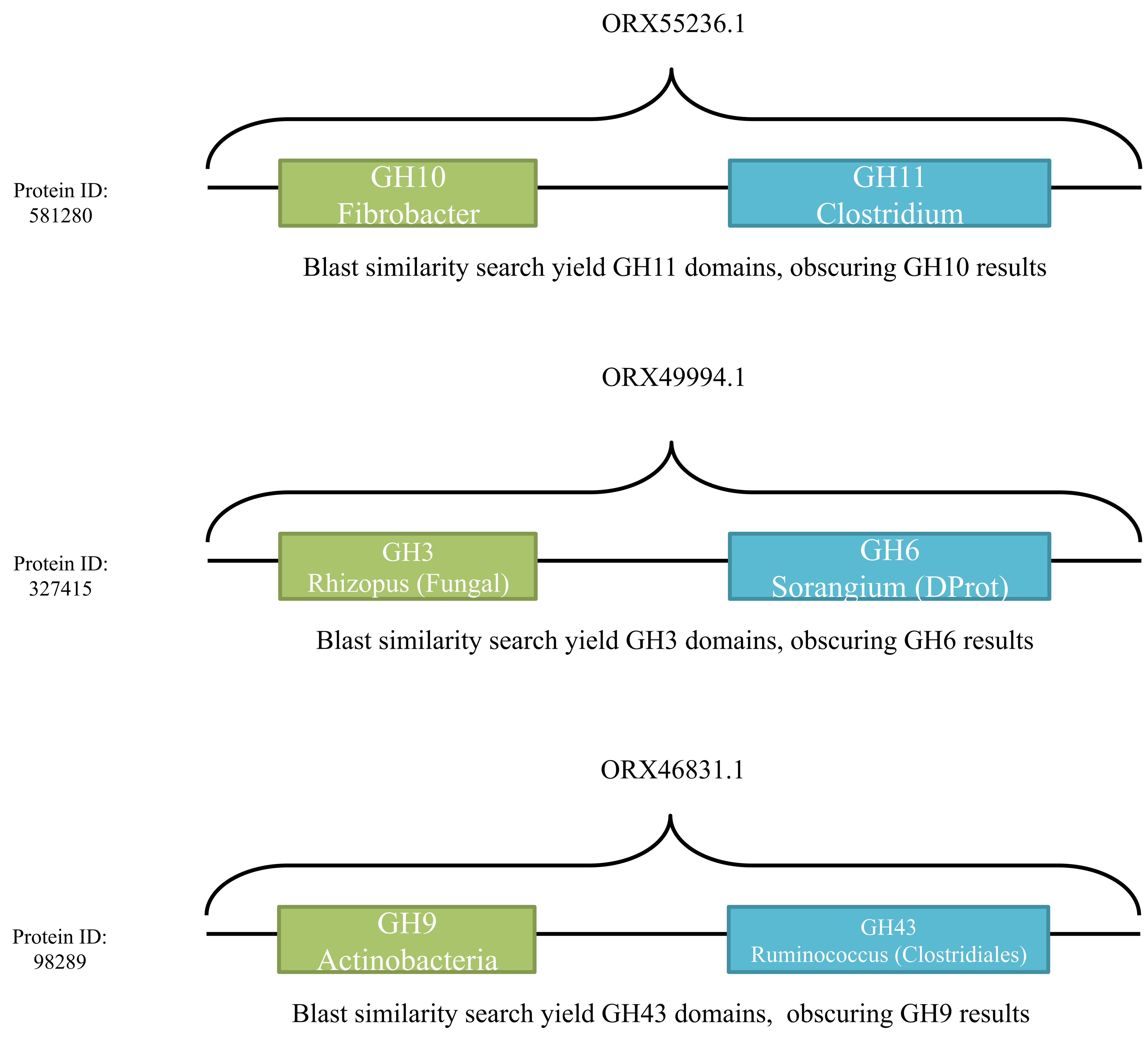

**Figure S3.** Maximum likelihood tree constructed using the D1–D2 domains of 28S rRNA gene. Isolates whose transcriptomes were sequenced in this study are shown in red, while publicly available transcriptomes included in the analysis are shown in blue. The tree was obtained using a maximum likelihood approach with Tajima-Nei model. Bootstrap values (from 100 replicates) are shown for nodes with more than 50% bootstrap support. Analysis was conducted in MEGA7 (5).

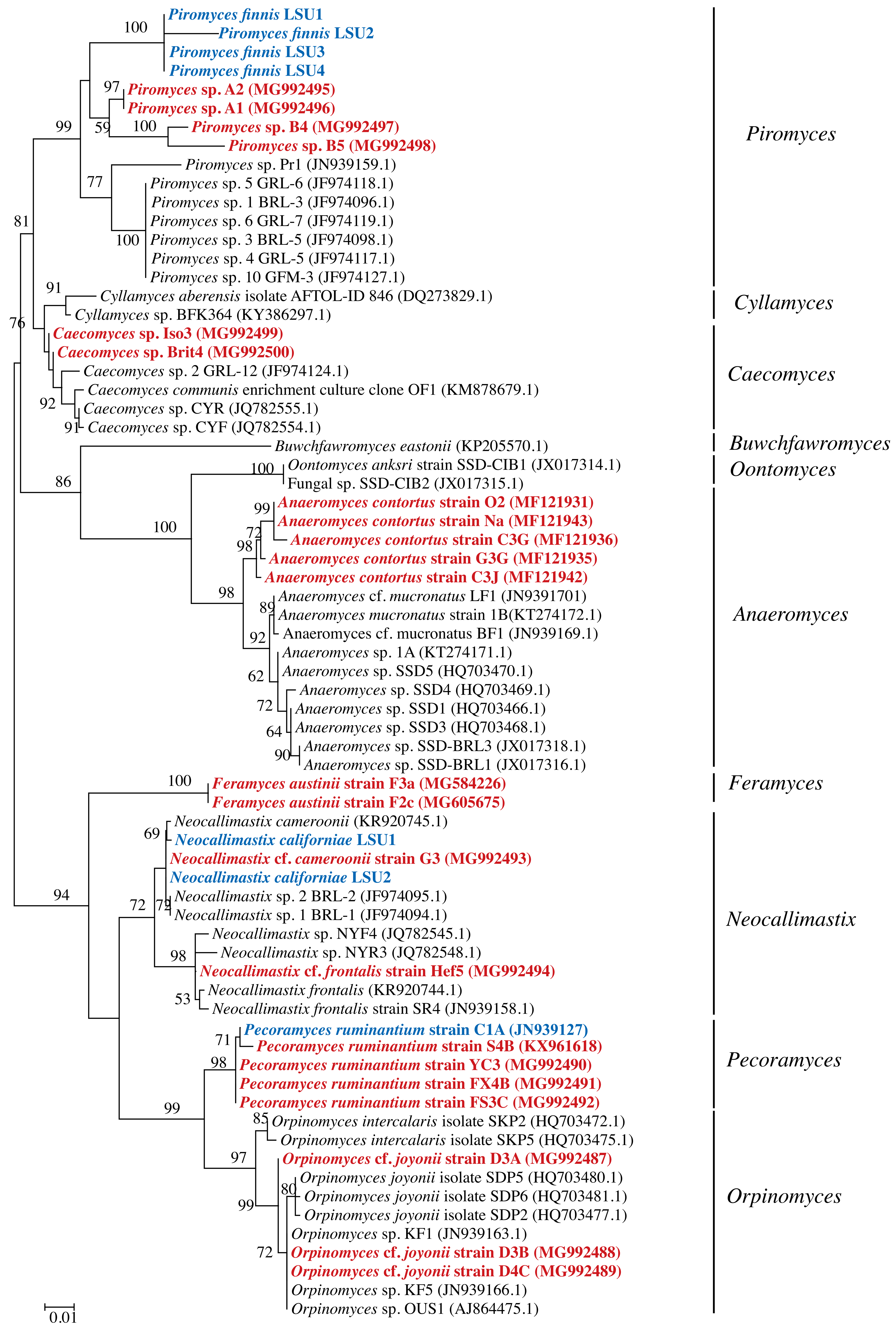

Piromyces finnis LSU1
Piromyces finnis LSU2
Piromyces finnis LSU3
Piromyces finnis LSU4
Piromyces sp. A2 (MG992495)
Piromyces sp. A1 (MG992496)
Piromyces sp. B4 (MG992497)
Piromyces sp. B5 (MG992498)
Piromyces sp. Pr1 (JN939159.1)
Piromyces sp. 5 GRL-6 (JF974118.1)
Piromyces sp. 1 BRL-3 (JF974096.1)
Piromyces sp. 6 GRL-7 (JF974119.1)
Piromyces sp. 3 BRL-5 (JF974098.1)
Piromyces sp. 4 GRL-5 (JF974117.1)
Piromyces sp. 10 GFM-3 (JF974127.1)
Cyllamyces aberensis isolate AFTOL-ID 846 (DQ273829.1)
Cyllamyces sp. BFK364 (KY386297.1)
Caecomyces sp. Iso3 (MG992499)
Caecomyces sp. Brit4 (MG992500)
Caecomyces sp. 2 GRL-12 (JF974124.1)
Caecomyces communis enrichment culture clone OF1 (KM878679.1)
Caecomyces sp. CYR (JQ782555.1)
Caecomyces sp. CYF (JQ782554.1)
Buwchfawromyces eastonii (KP205570.1)
Oontomyces anksri strain SSD-CIB1 (JX017314.1)
Fungal sp. SSD-CIB2 (JX017315.1)
Anaeromyces contortus strain O2 (MF121931)
Anaeromyces contortus strain Na (MF121943)
Anaeromyces contortus strain C3G (MF121936)
Anaeromyces contortus strain G3G (MF121935)
Anaeromyces contortus strain C3J (MF121942)
Anaeromyces cf. mucronatus LF1 (JN9391701)
Anaeromyces mucronatus strain 1B(KT274172.1)
Anaeromyces cf. mucronatus BF1 (JN939169.1)
Anaeromyces sp. 1A (KT274171.1)
Anaeromyces sp. SSD5 (HQ703470.1)
Anaeromyces sp. SSD4 (HQ703469.1)
Anaeromyces sp. SSD1 (HQ703466.1)
Anaeromyces sp. SSD3 (HQ703468.1)
Anaeromyces sp. SSD-BRL3 (JX017318.1)
Anaeromyces sp. SSD-BRL1 (JX017316.1)
Feramyces austinii strain F3a (MG584226)
Feramyces austinii strain F2c (MG605675)
Neocallimastix cameroonii (KR920745.1)
Neocallimastix californiae LSU1
Neocallimastix cf. cameroonii strain G3 (MG992493)
Neocallimastix californiae LSU2
Neocallimastix sp. 2 BRL-2 (JF974095.1)
Neocallimastix sp. 1 BRL-1 (JF974094.1)
Neocallimastix sp. NYF4 (JQ782545.1)
Neocallimastix sp. NYR3 (JQ782548.1)
Neocallimastix cf. frontalis strain Hef5 (MG992494)
Neocallimastix frontalis (KR920744.1)
Neocallimastix frontalis strain SR4 (JN939158.1)
Pecoramyces ruminantium strain C1A (JN939127)
Pecoramyces ruminantium strain S4B (KX961618)
Pecoramyces ruminantium strain YC3 (MG992490)
Pecoramyces ruminantium strain FX4B (MG992491)
Pecoramyces ruminantium strain FS3C (MG992492)
Orpinomyces intercalaris isolate SKP2 (HQ703472.1)
Orpinomyces intercalaris isolate SKP5 (HQ703475.1)
Orpinomyces cf. joyonii strain D3A (MG992487)
Orpinomyces joyonii isolate SDP5 (HQ703480.1)
Orpinomyces joyonii isolate SDP6 (HQ703481.1)
Orpinomyces joyonii isolate SDP2 (HQ703477.1)
Orpinomyces sp. KF1 (JN939163.1)
Orpinomyces cf. joyonii strain D3B (MG992488)
Orpinomyces cf. joyonii strain D4C (MG992489)
Orpinomyces sp. KF5 (JN939166.1)
Orpinomyces sp. OUS1 (AJ864475.1)
Piromyces
Cyllamyces
Caecomyces
Buwchfawromyces
Oontomyces
Anaeromyces
Feramyces
Neocallimastix
Pecoramyces
Orpinomyces
0.01

**Figure S4. Mapping frequency of HGT events in various AGF clades and genera.** The number of HGT events mapped on the nodes are in red. The majority of AGF events occurred prior to Genus level diversification, while few (51 events) appearing to be genus specific. Maximum likelihood tree constructed using the D1–D2 domains of 28S rRNA gene. Only isolates whose transcriptomes were analyzed in this study are shown. Bootstrap values (from 100 replicates) are shown for nodes with 50-70% bootstrap support as grey circles and for nodes with 70-100% bootstrap support as black circles. The size of the circle is proportional to the bootstrap value. Analysis was conducted in MEGA7 (5).

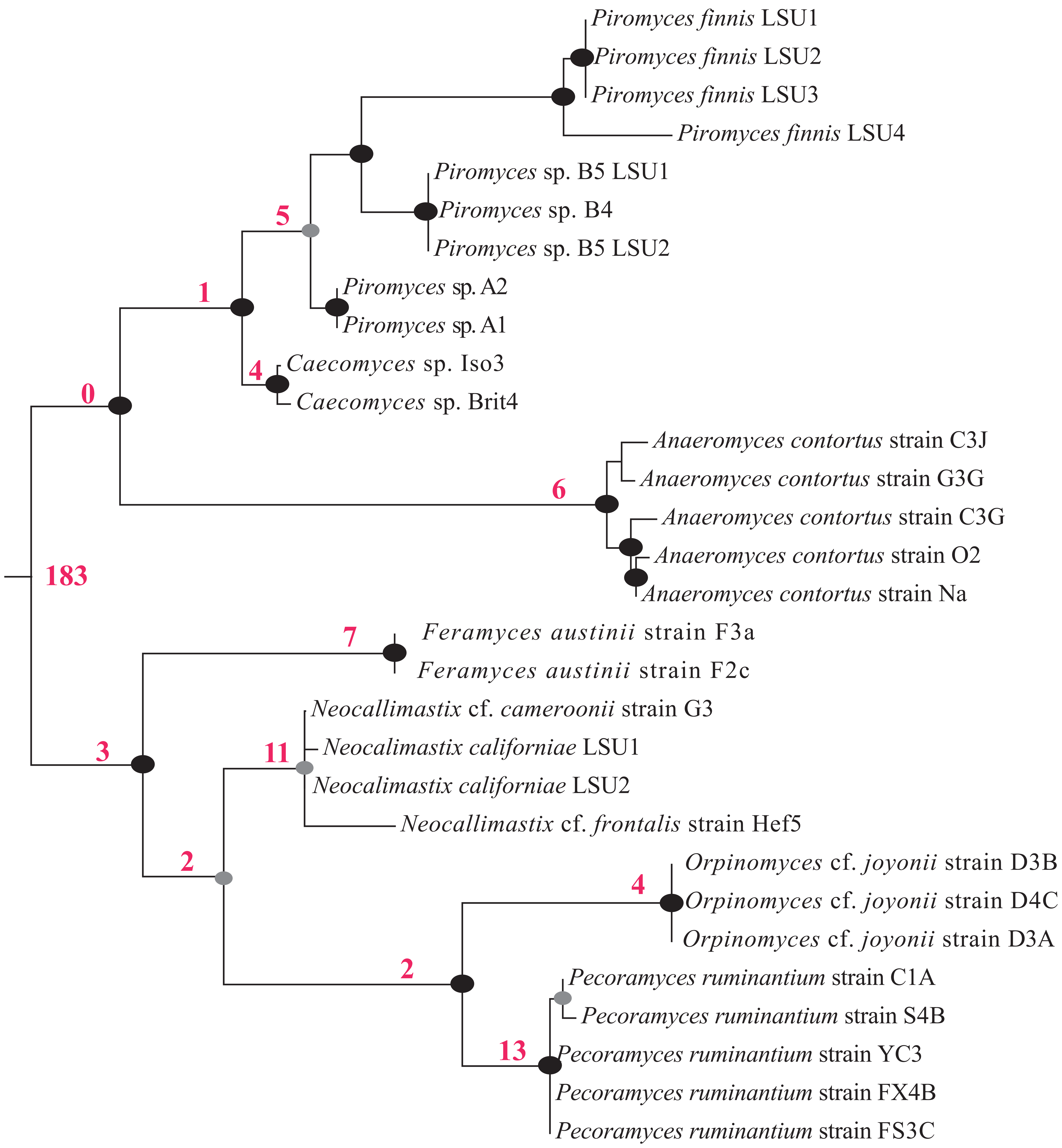

**Figures S5-S45.** Maximum likelihood trees showing the phylogenetic affiliation of 52 different horizontally transferred genes. The predicted function is shown in the upper left corner of each tree. AGF genes (shown in blue) are clearly nested within non-fungal clusters with strong bootstrap support reinforcing the inference of horizontal transfer. GenBank accession numbers are shown for reference sequences.

Fig S5. Aldose-1-epimerase

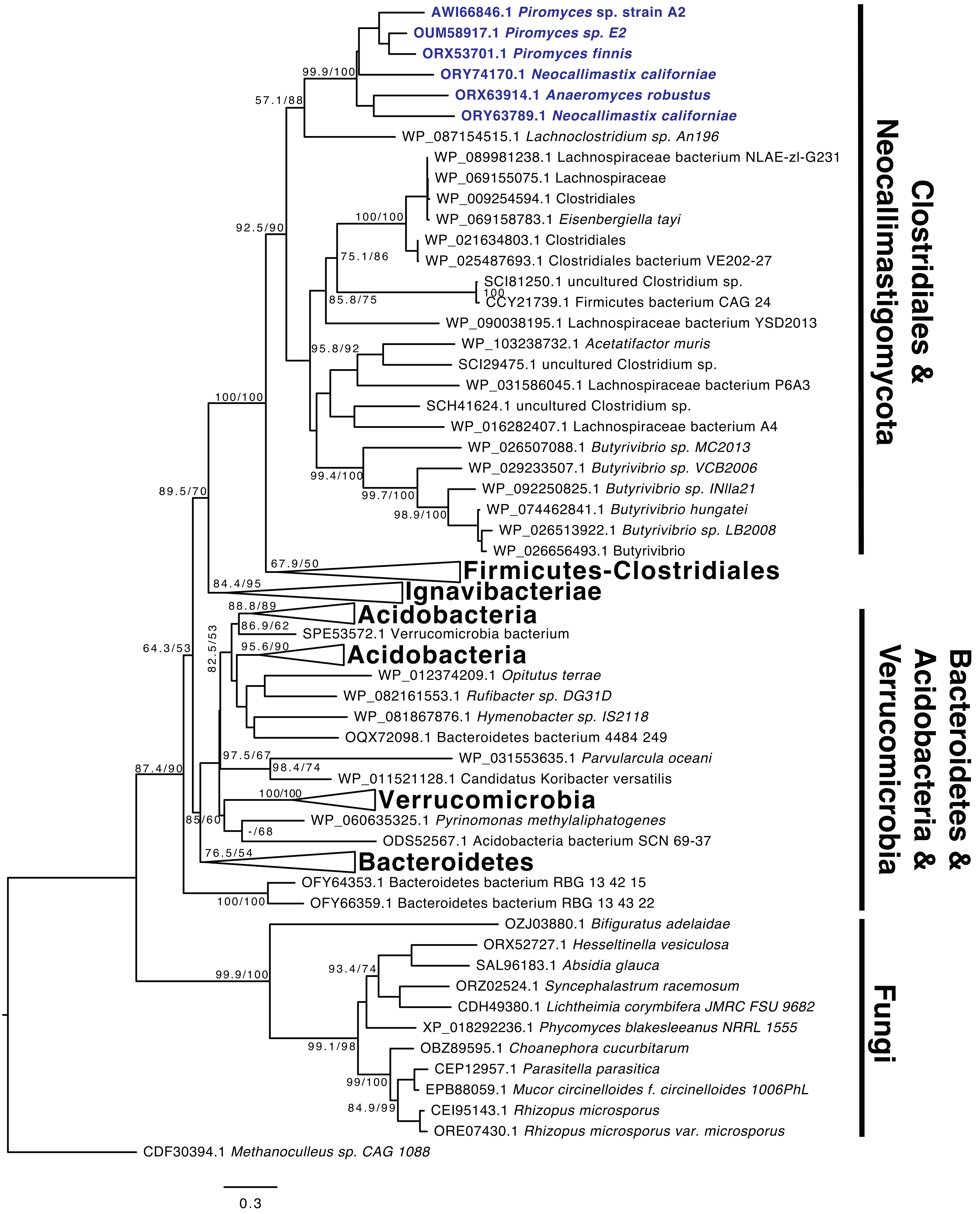

Fig S6. Galactose-1-P uridyltransferase

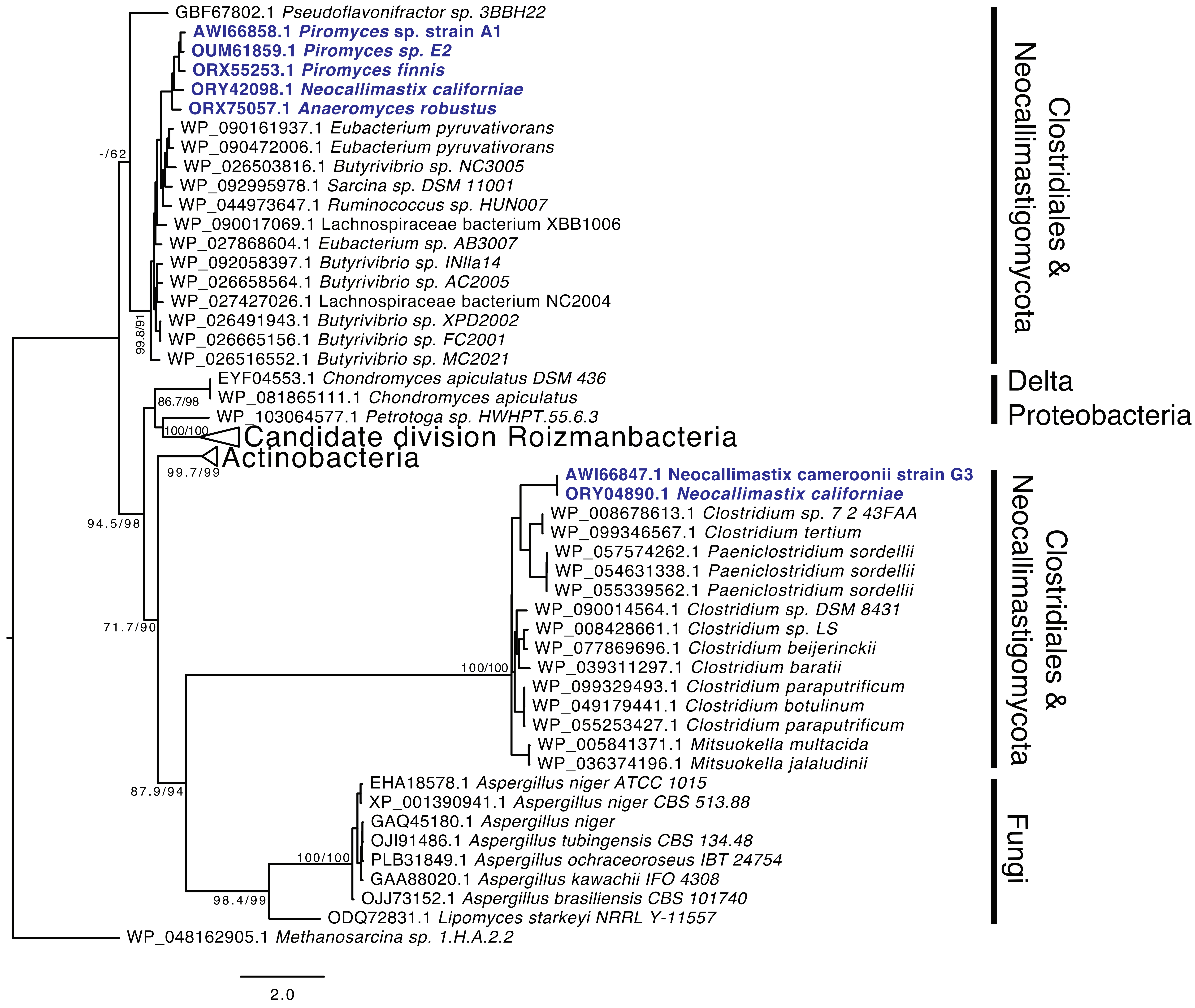

Fig S7. Ribokinase

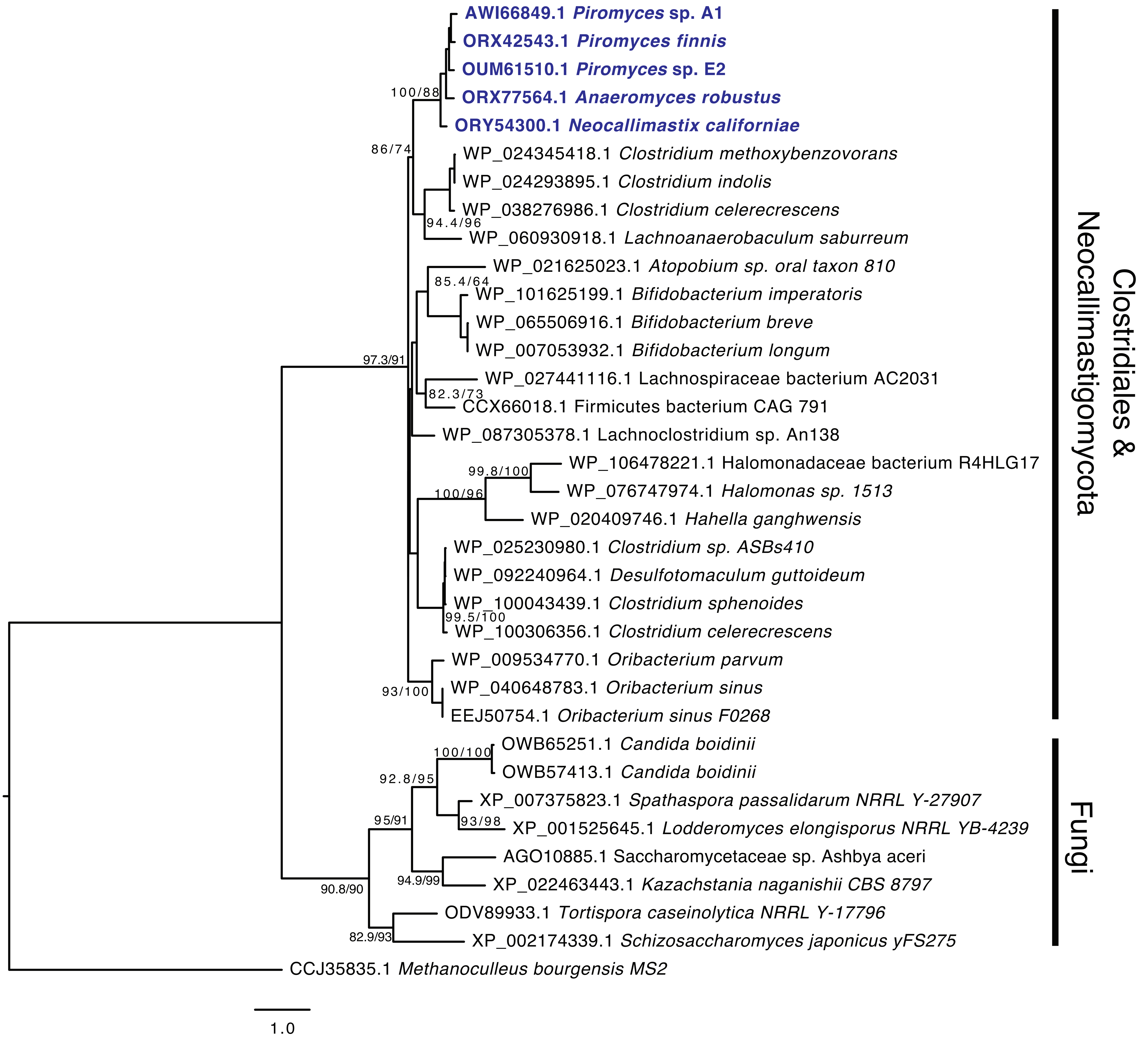

Fig S8. Xylose isomerase

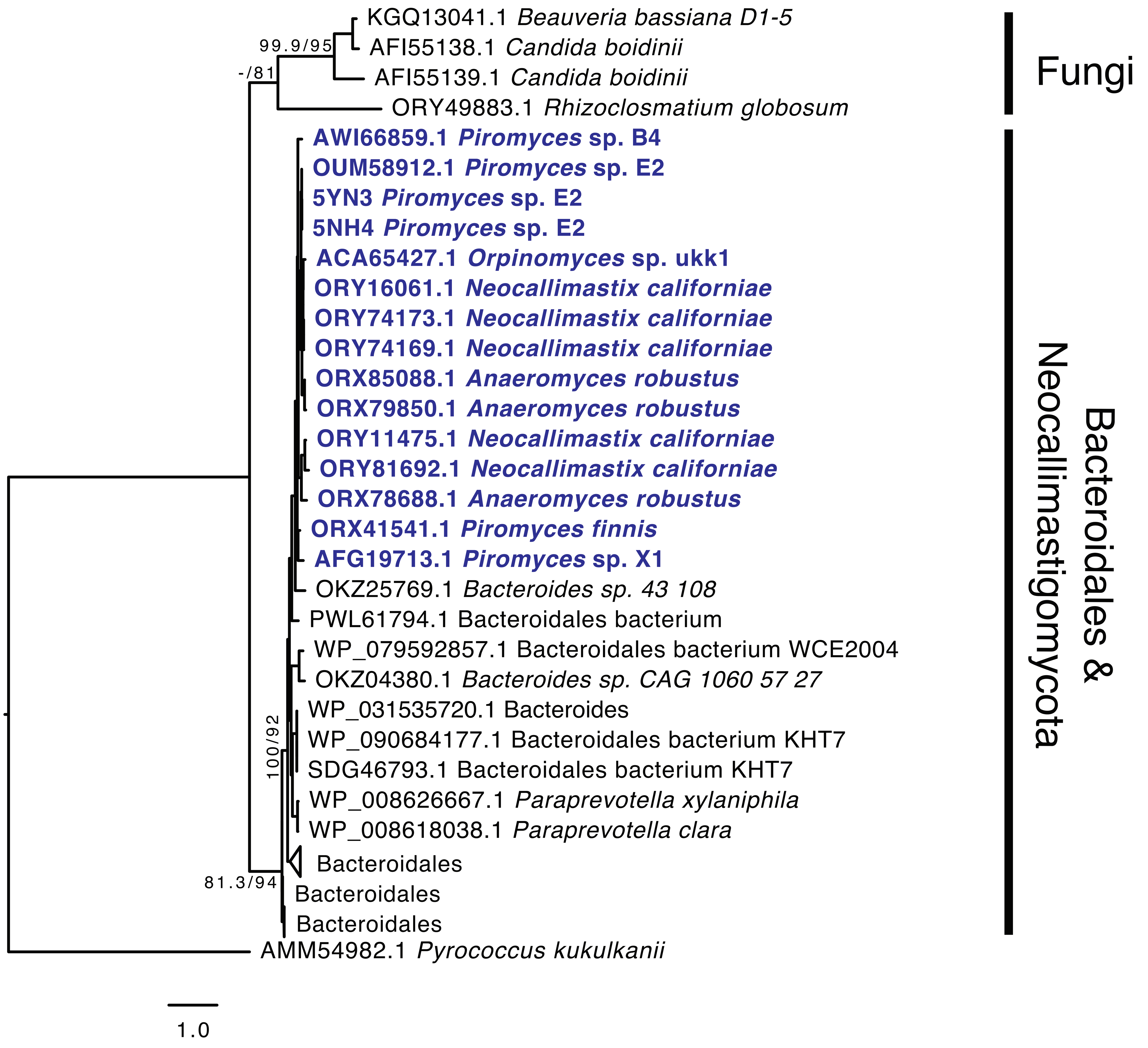

Fig S9. Xylulokinase

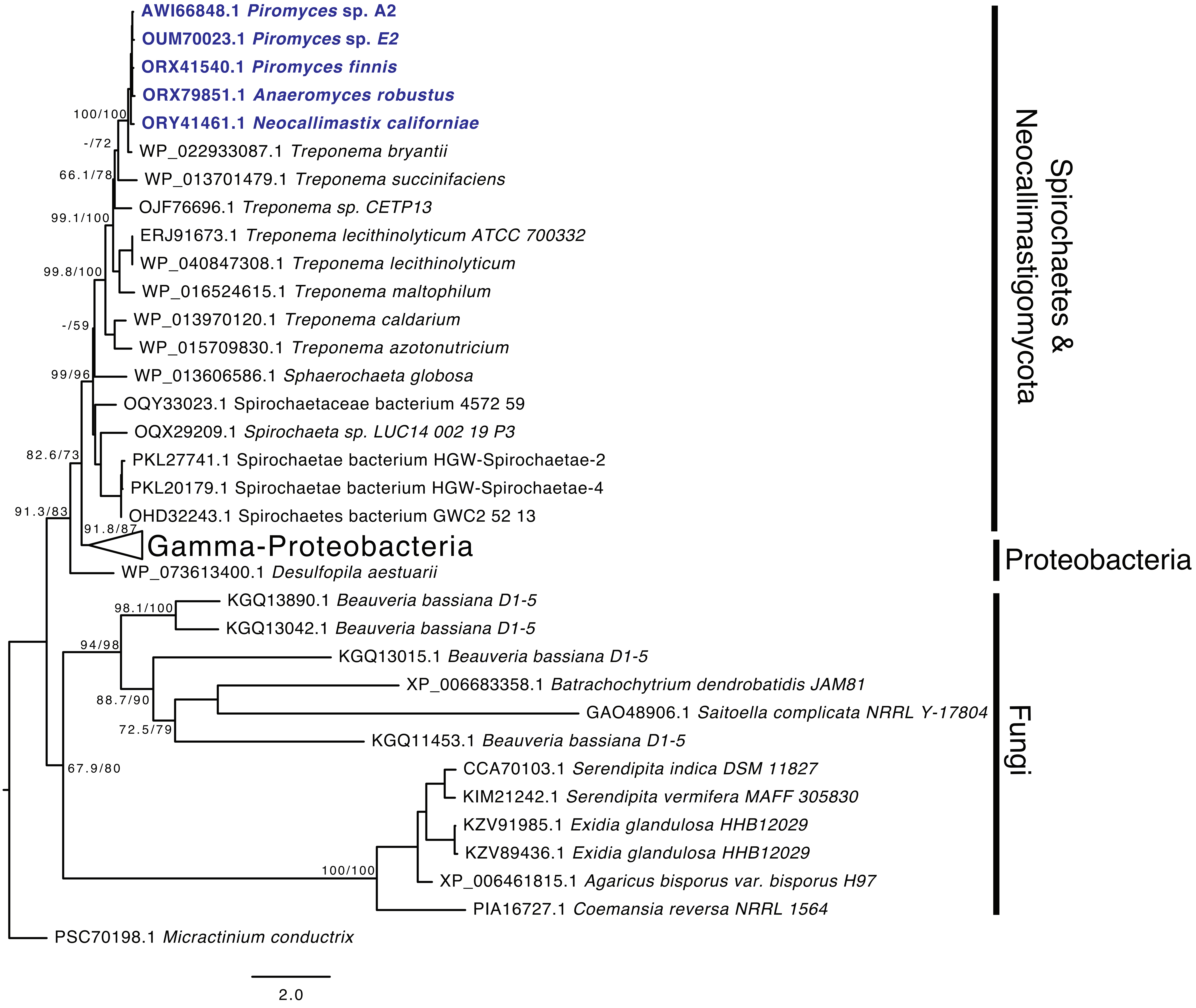

Fig S10. Deoxyribose-phosphate aldolase

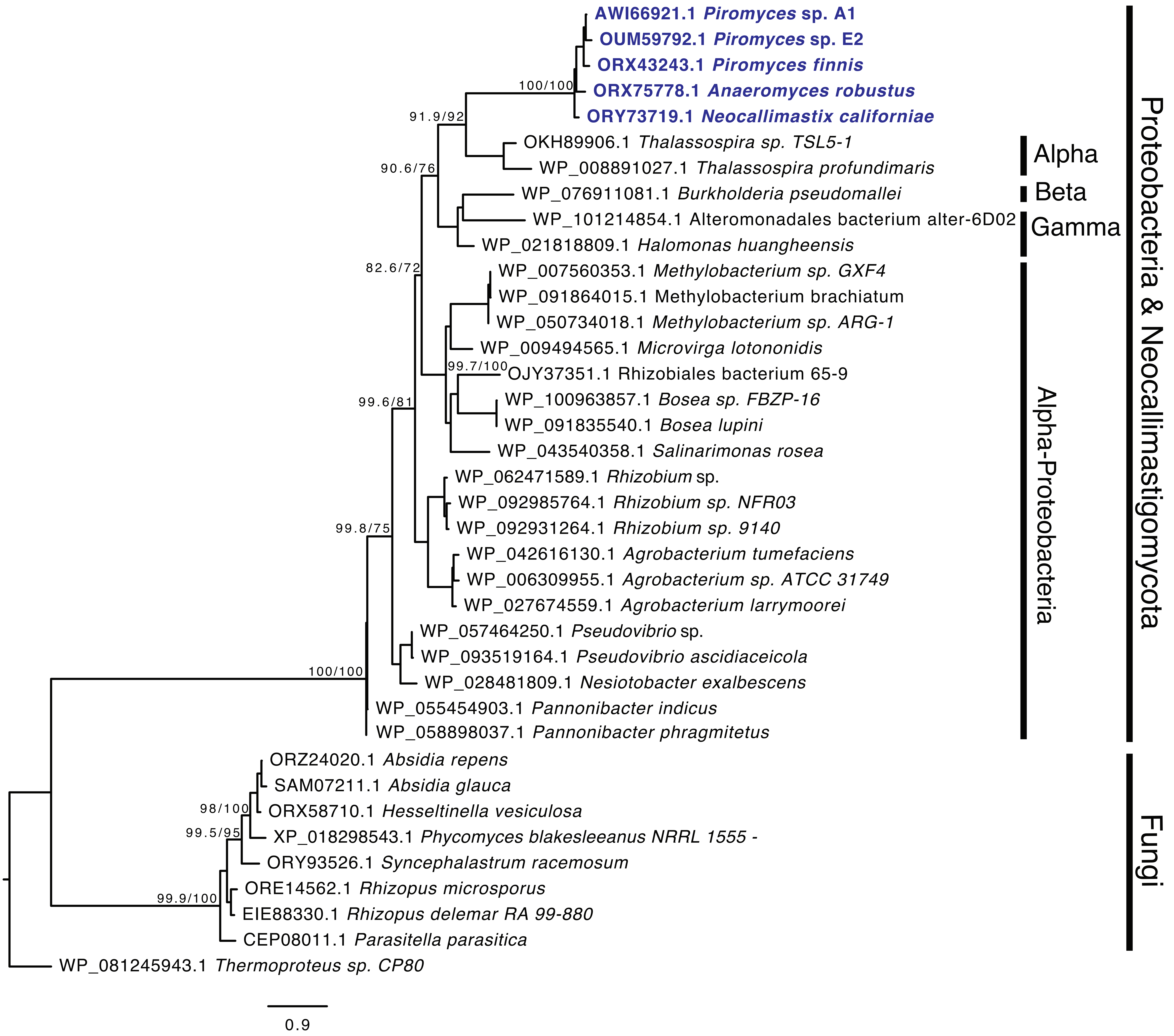

Fig S11. Phosphoenol pyruvate synthase

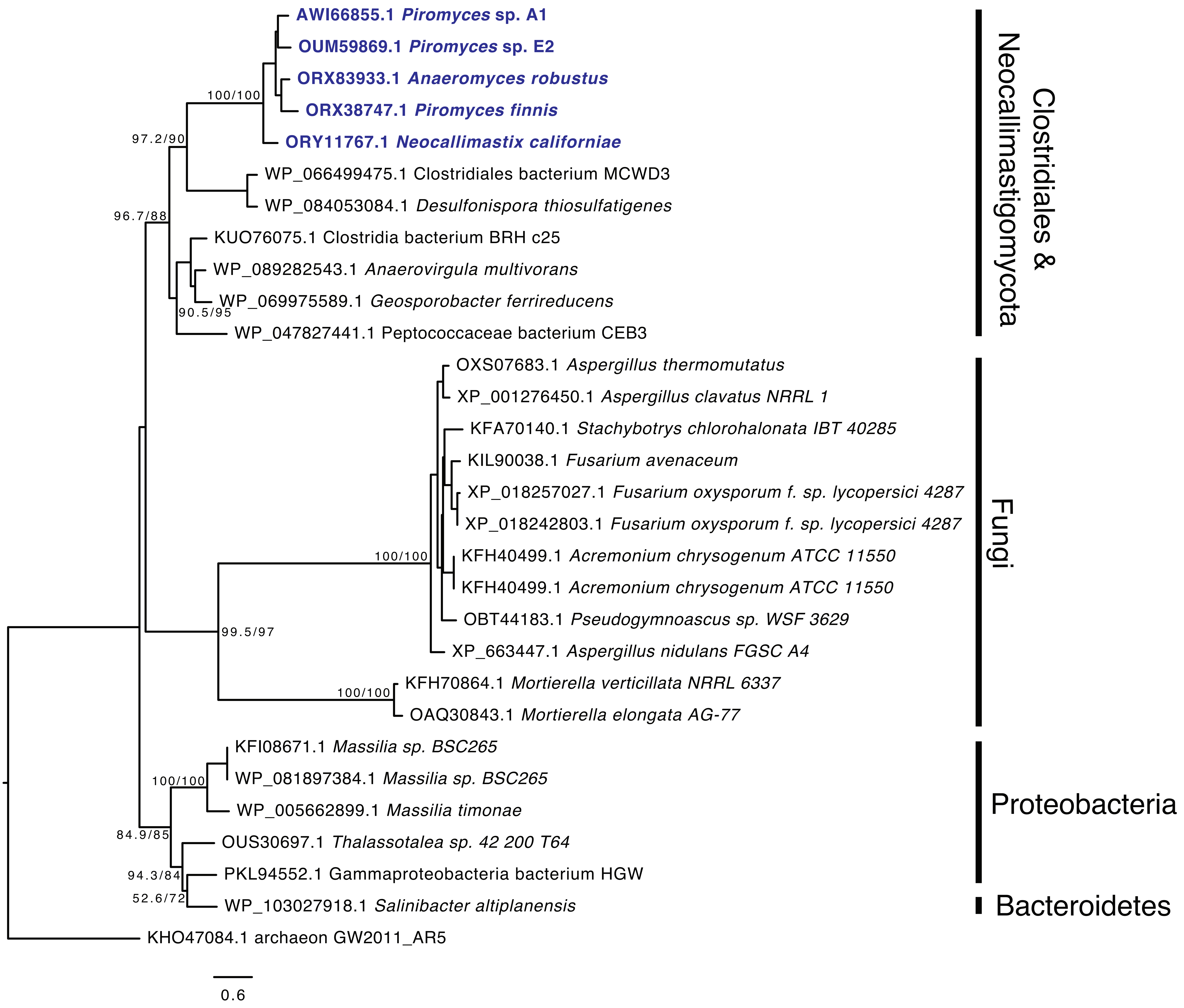

Fig S12. Bisphosphoglycerate-dependent phosphoglycerate mutase

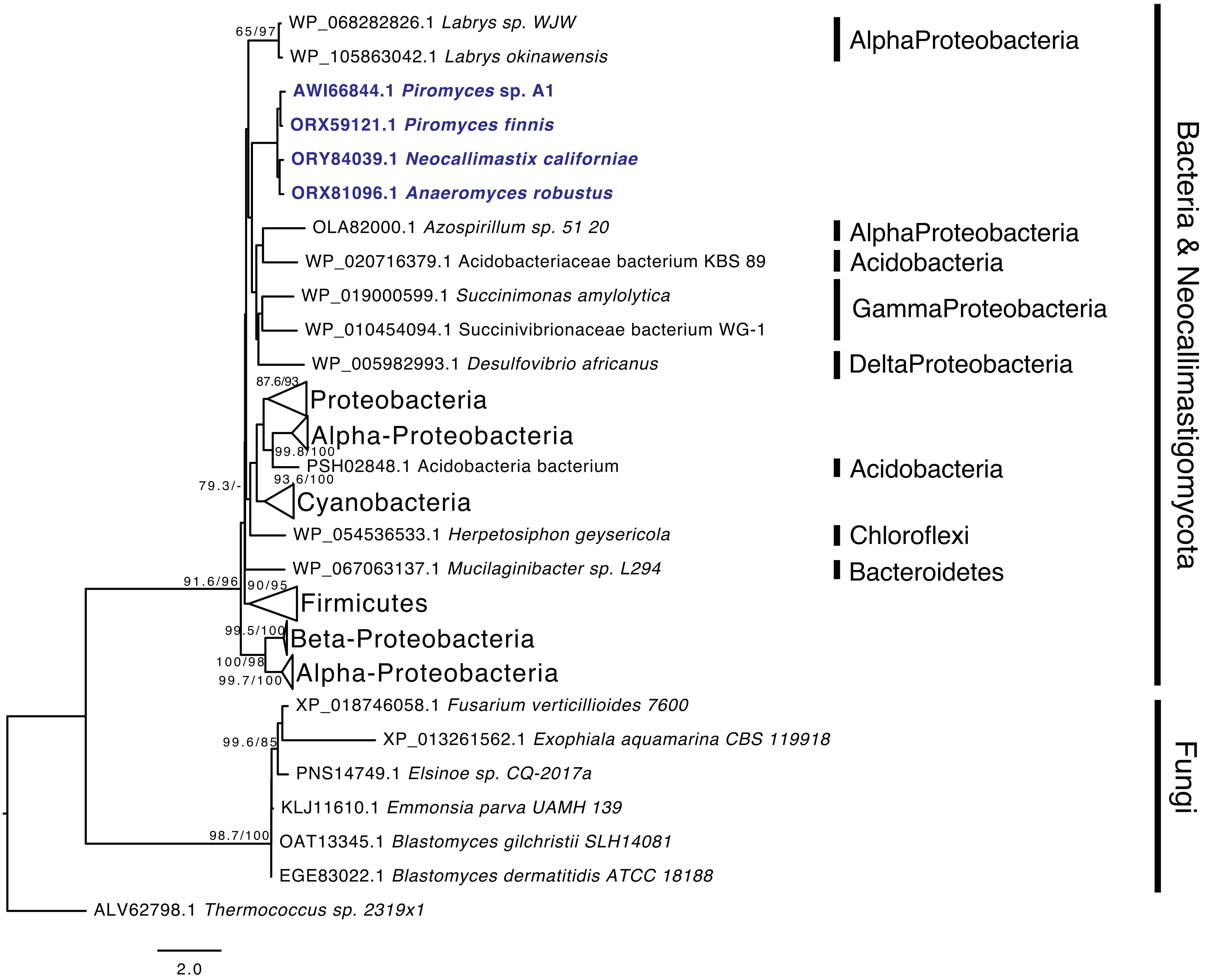

Fig S13. Bisphosphoglycerate-independent phosphoglycerate mtase

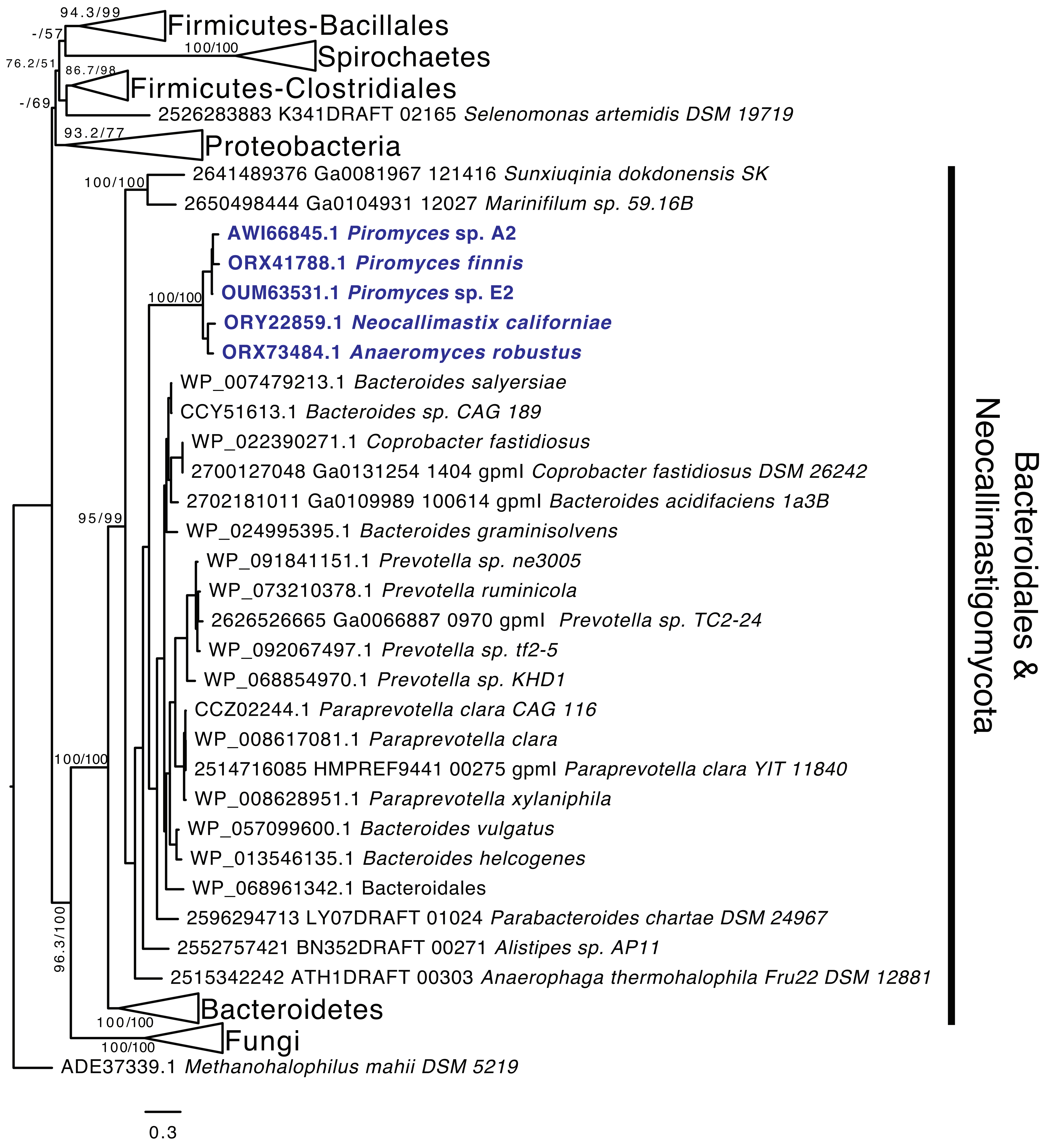

Fig S14. Bifunctional aldehyde/alcohol dehydrogenase

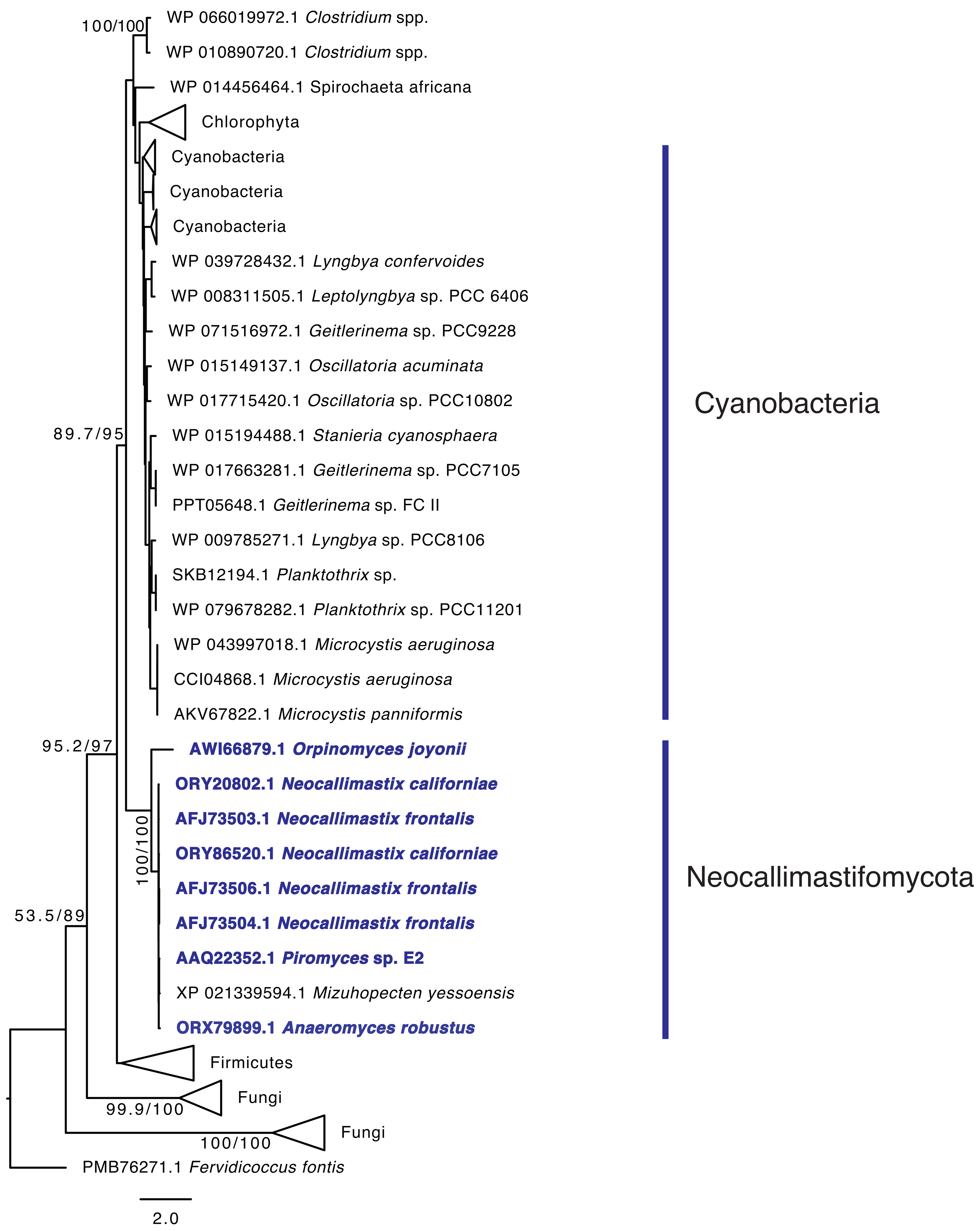

Fig S15. D-lactate dehydrogenase

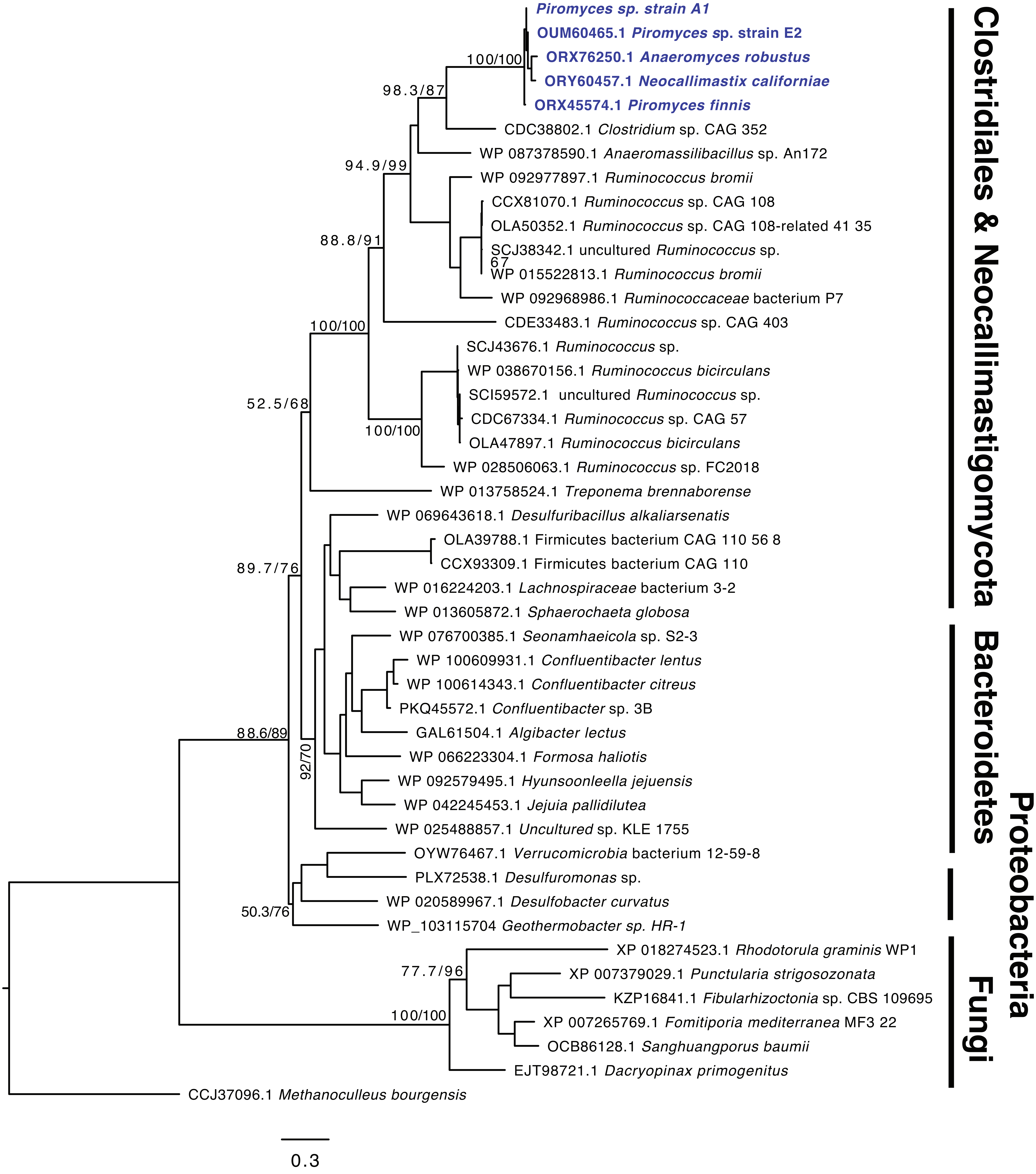

Fig S16. Fe-only hydrogenase maturation enzyme HydE

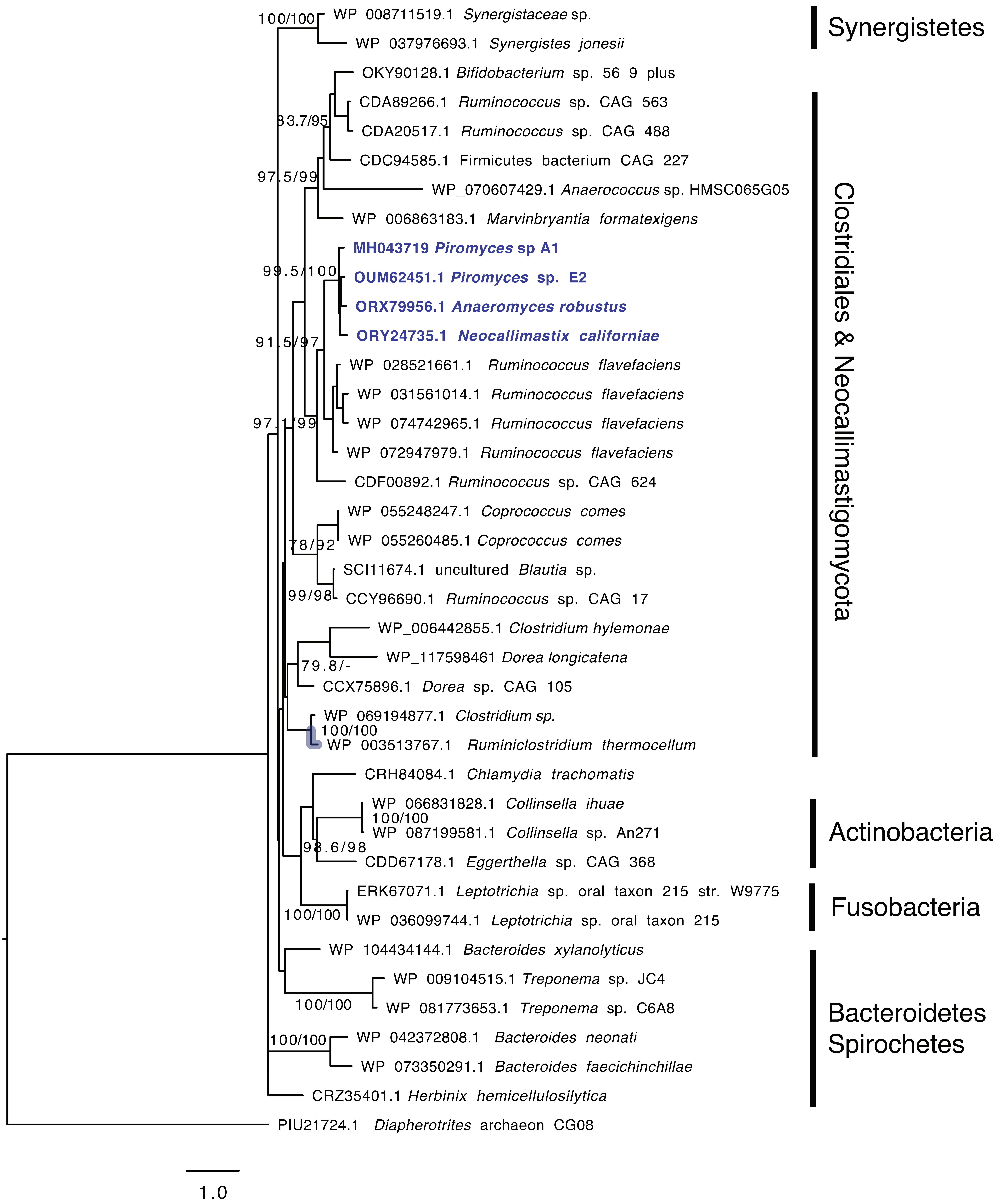

Fig S17. Serine-O-acetyltransferase

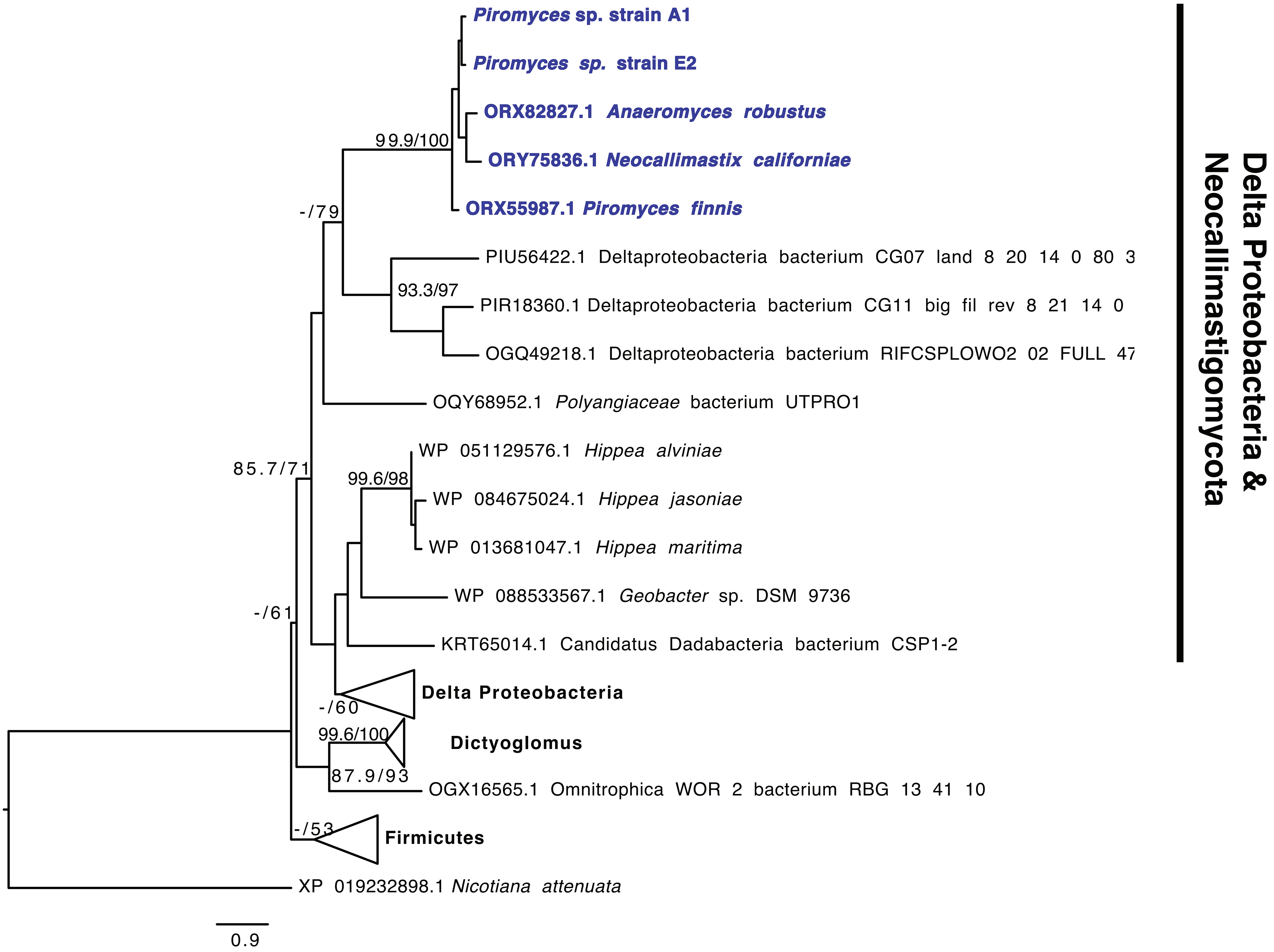

Fig S18. O-acetylhomoserine/O-acetylserine sulfhydrylase

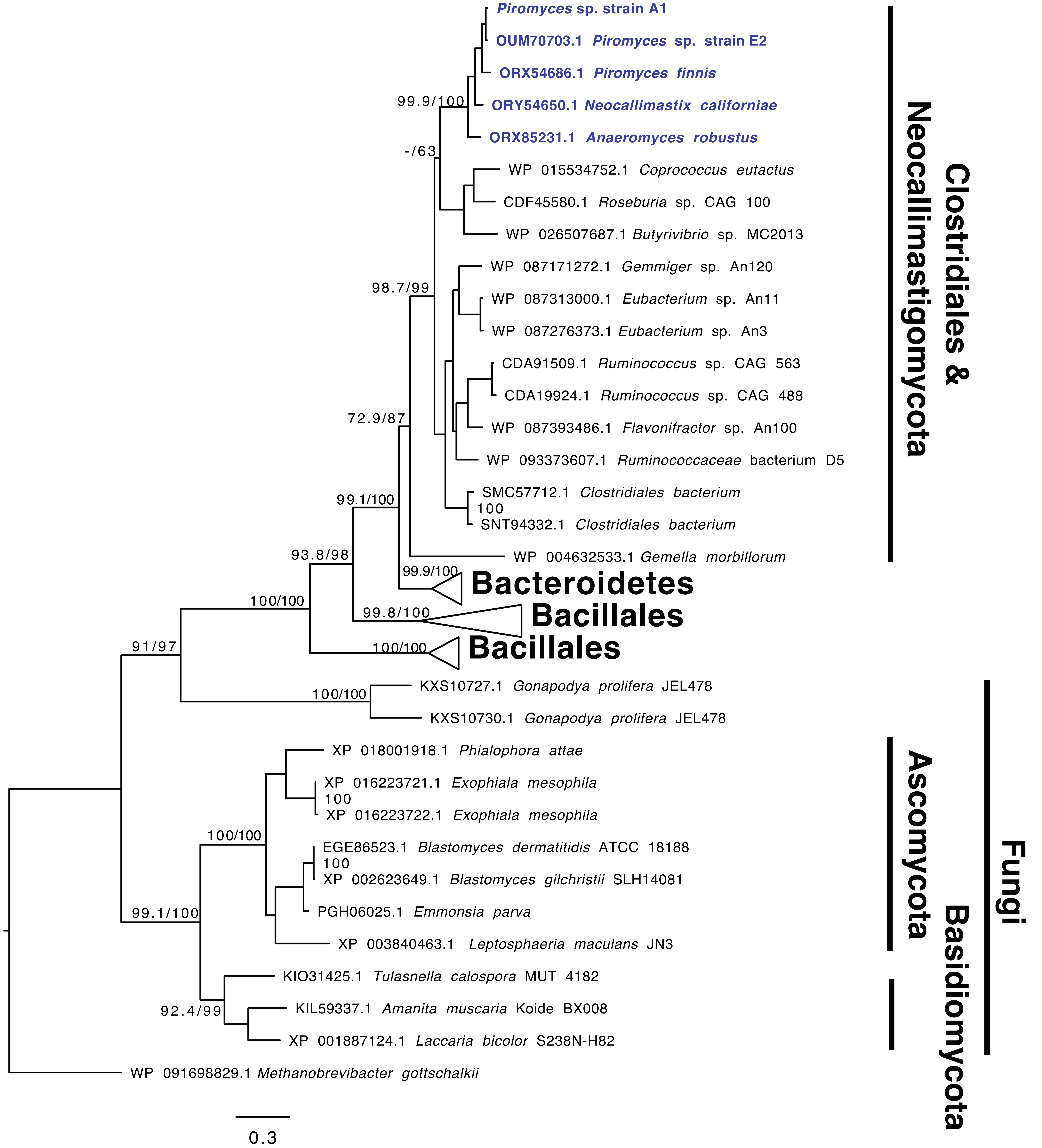

Fig S19. Cysteine synthase

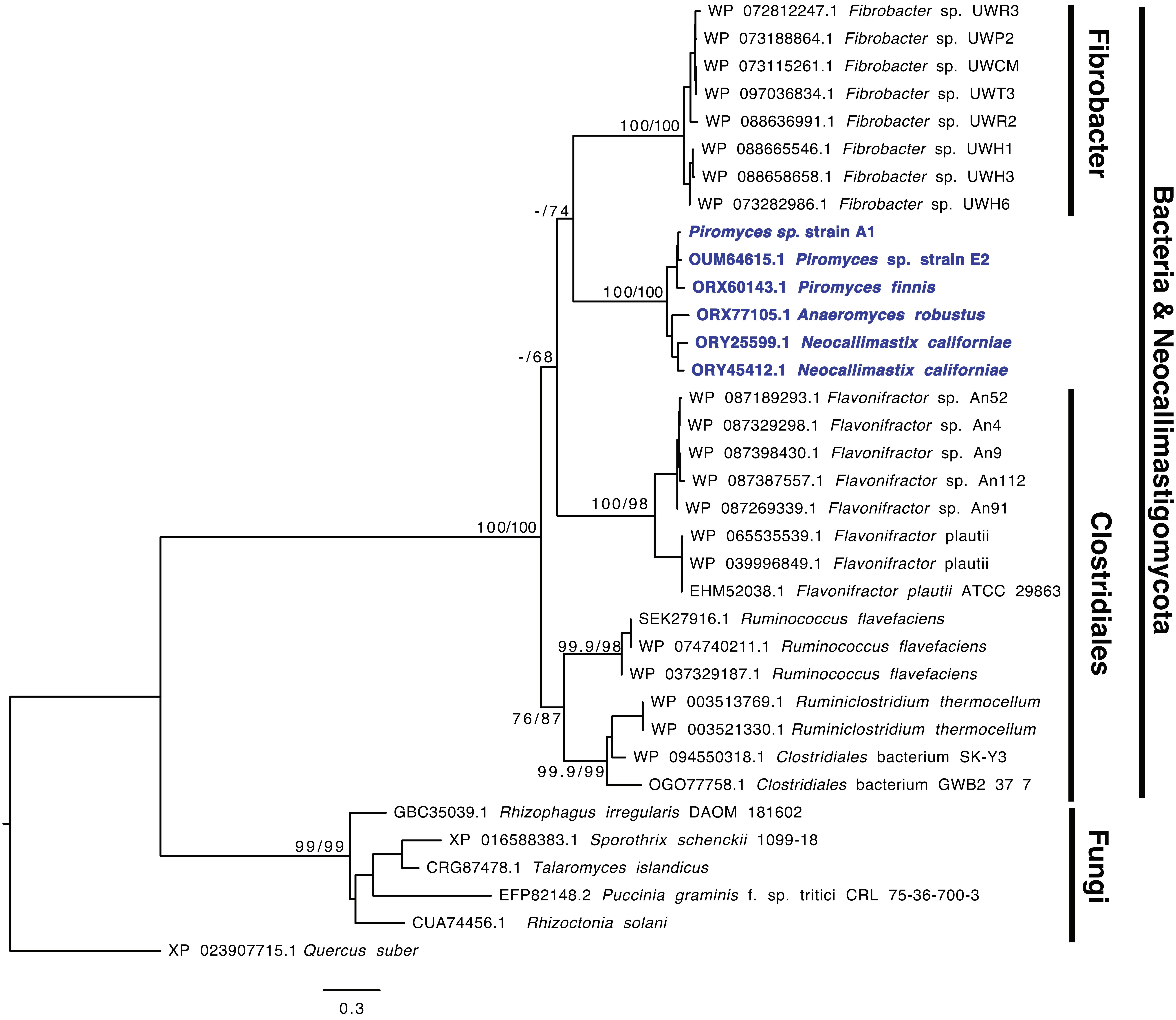

Fig S20. Low-specificity L-threonine aldolase

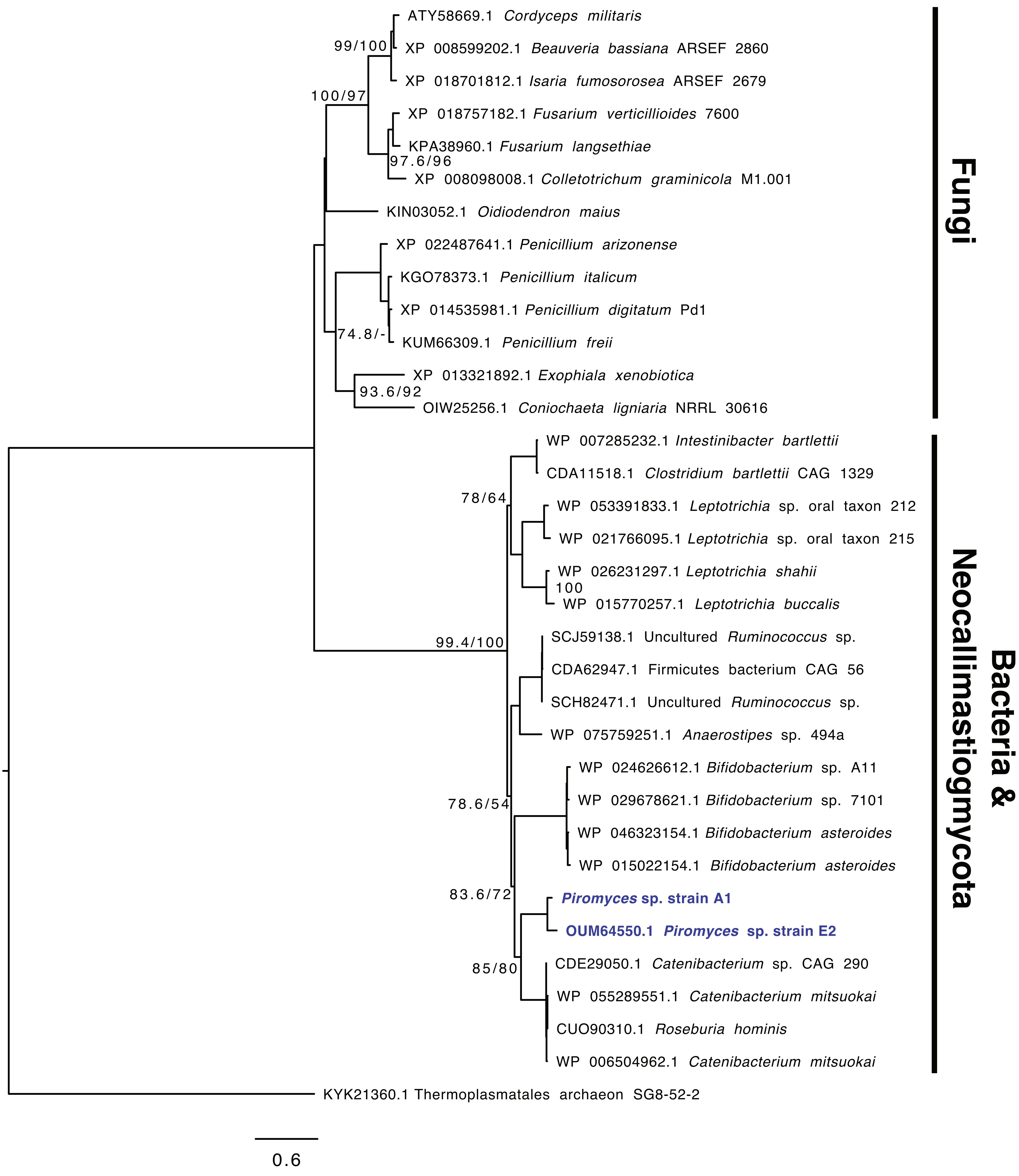

Fig S21. Aspartate-ammonia ligase

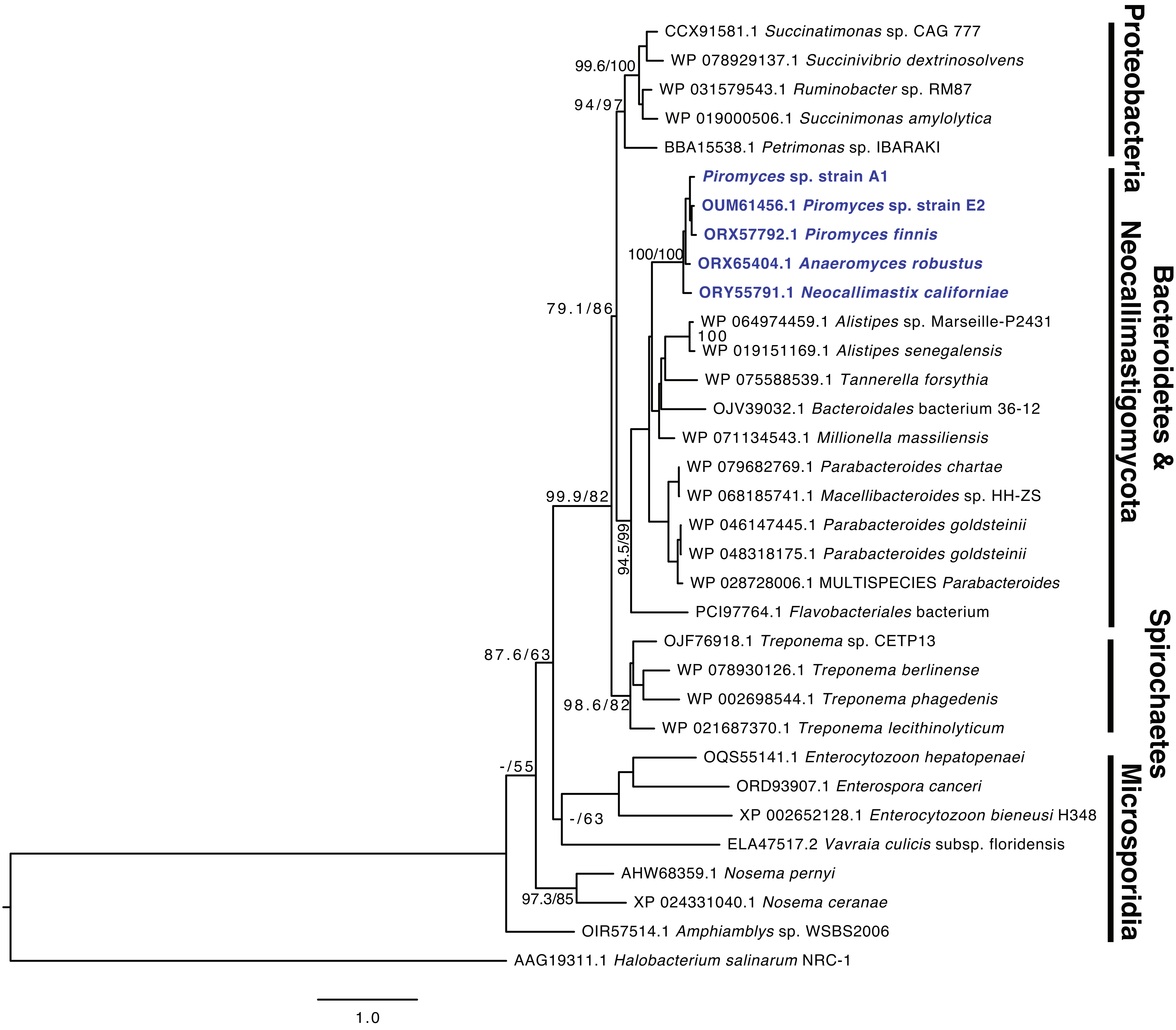

Fig S22. NadA (Quinolinate synthase)

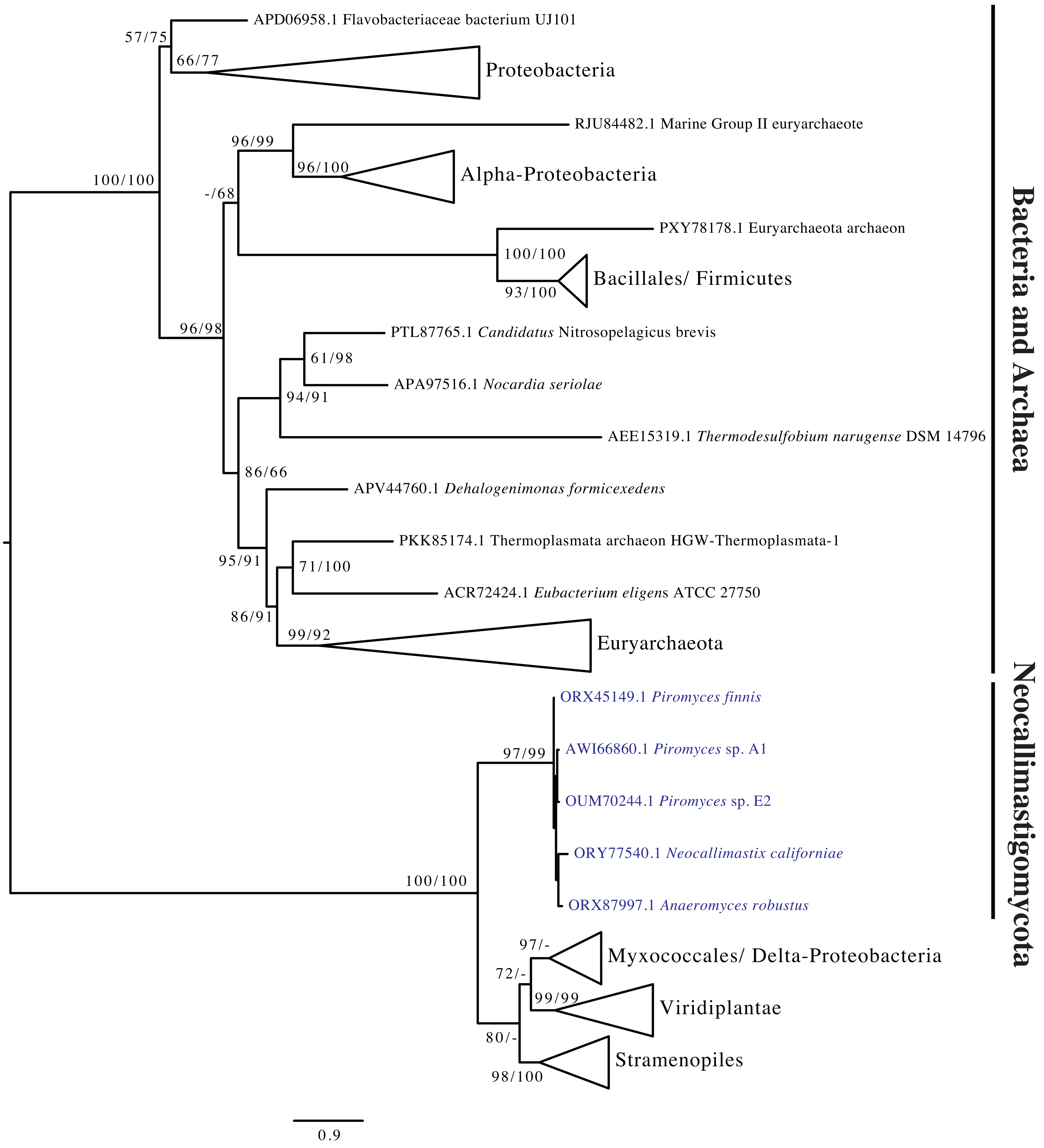

Fig S23. Phosphomethyl pyrimidine kinase

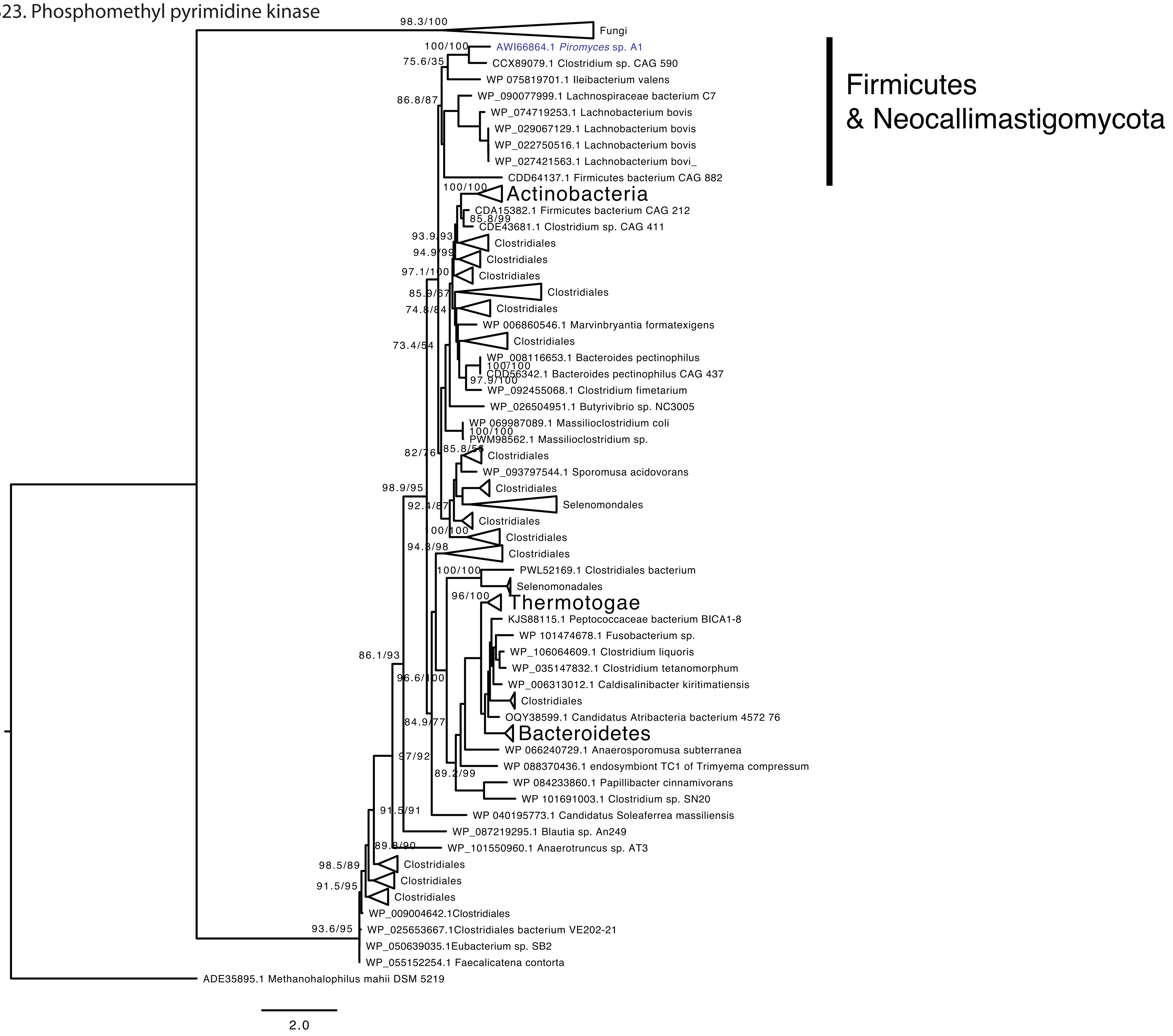

Fig S24. GMP reductase

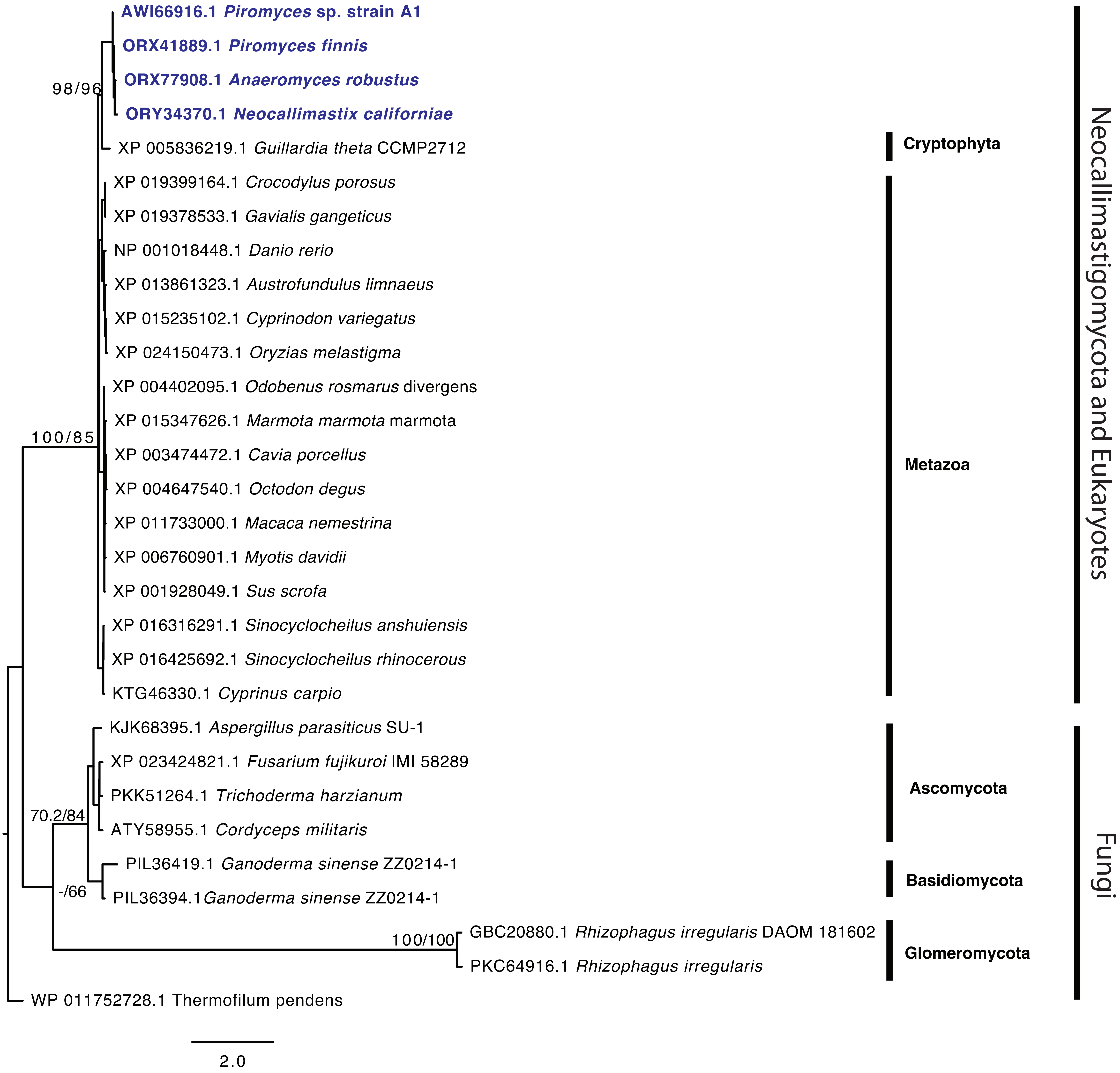

Fig S25. Nucleoside/nucleotide kinase family protein

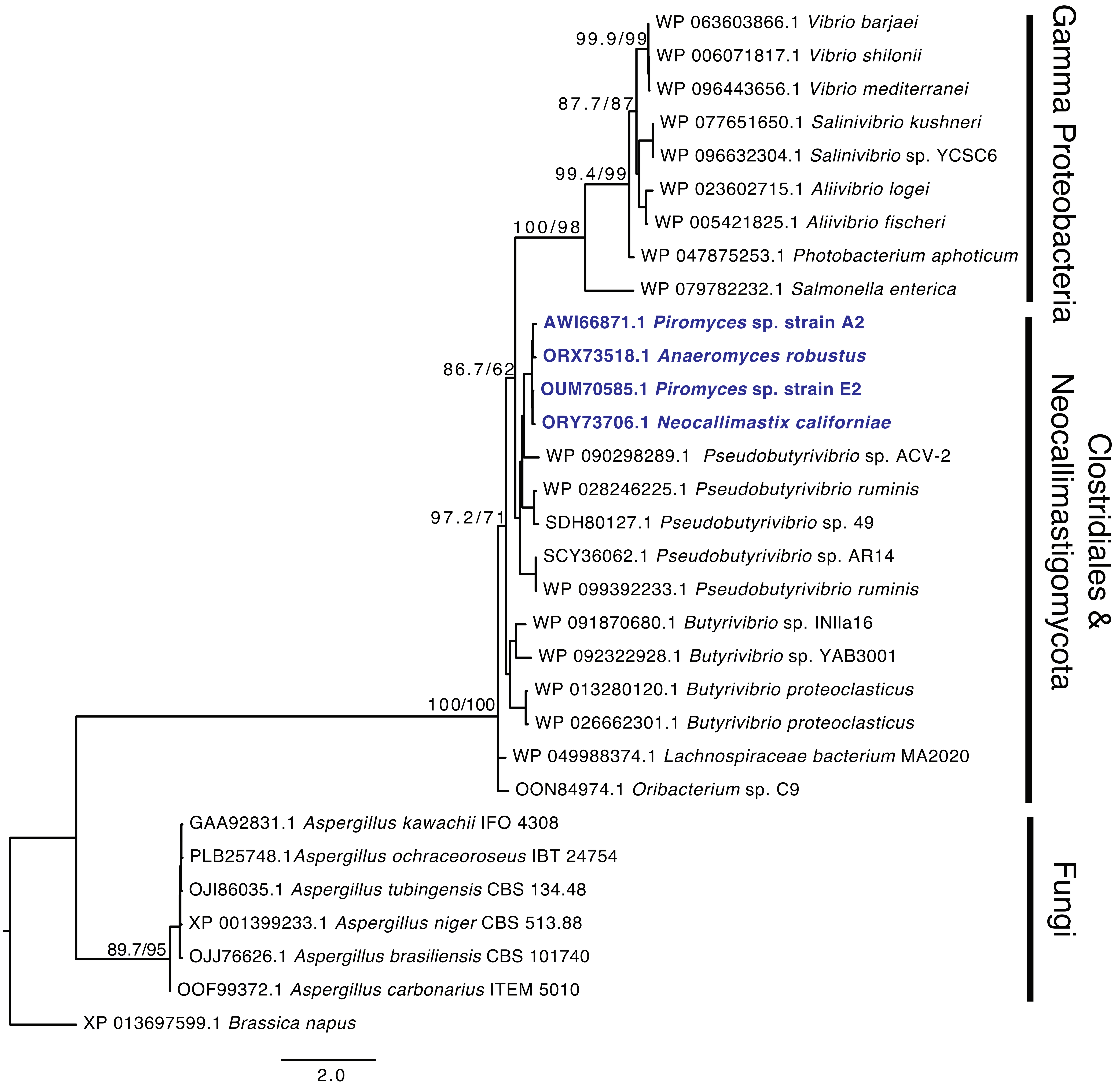

Fig S26. Cytidylate kinase

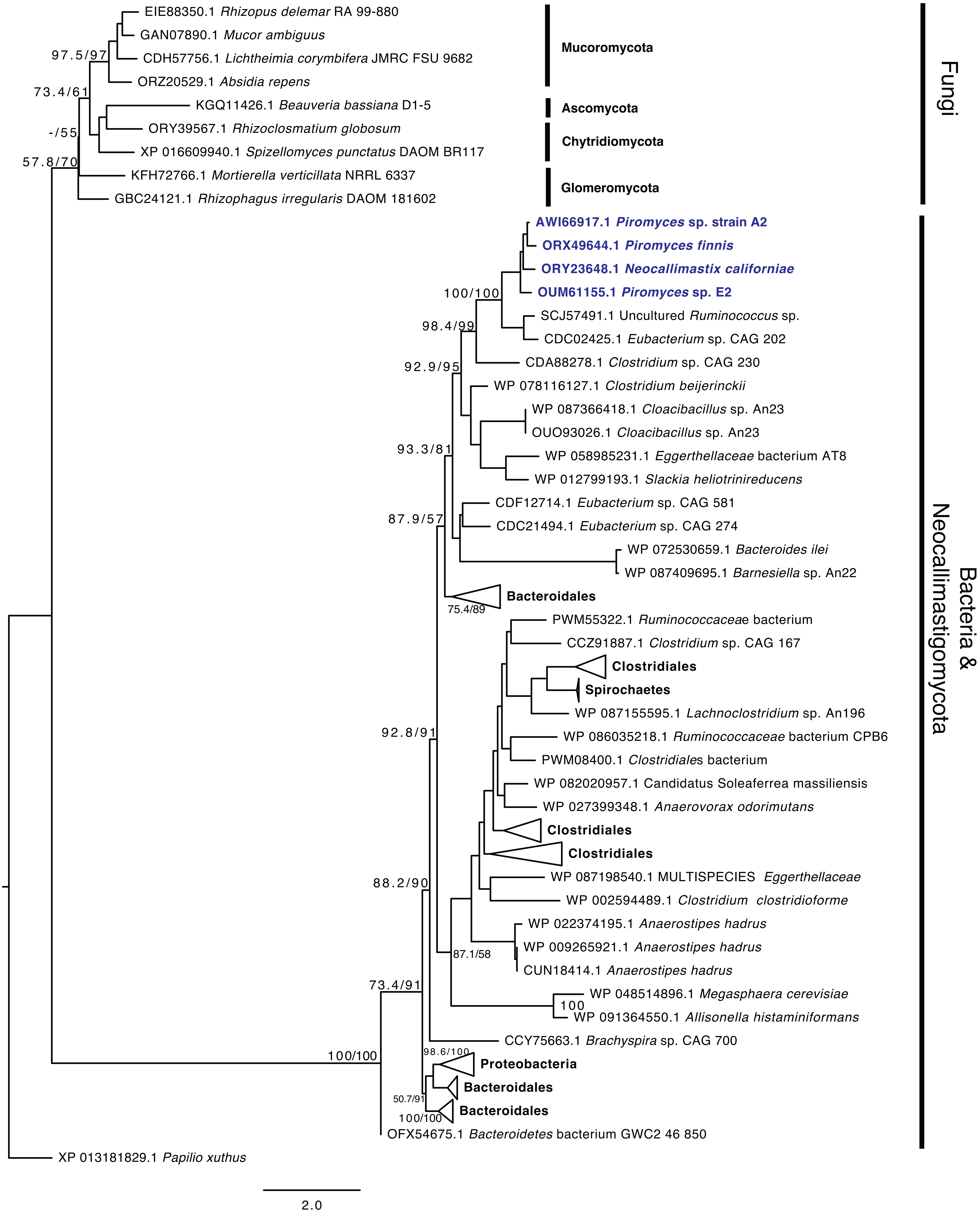

Fig S27. Thymidylate synthase

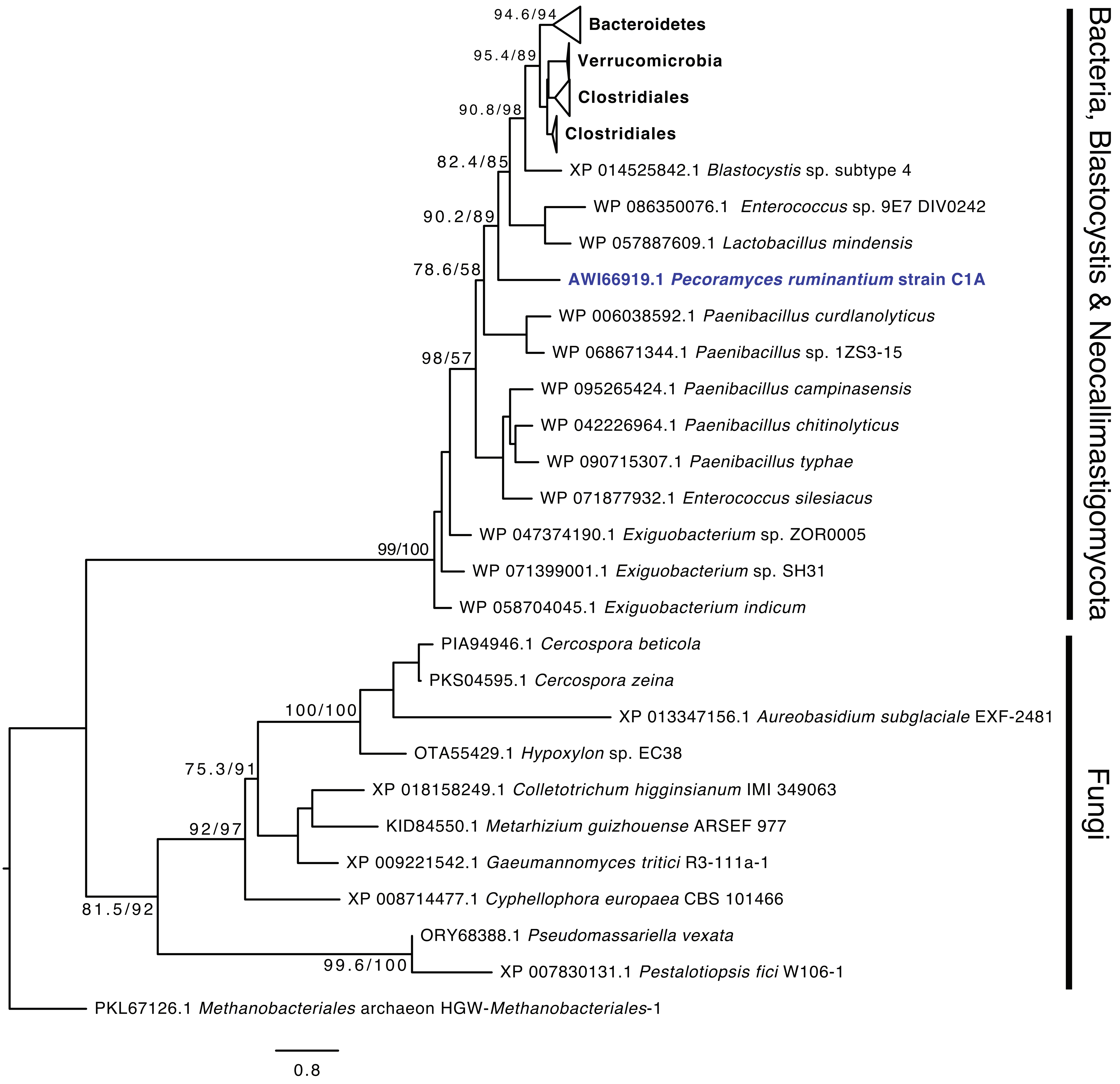

Fig S28. Thymidine kinase

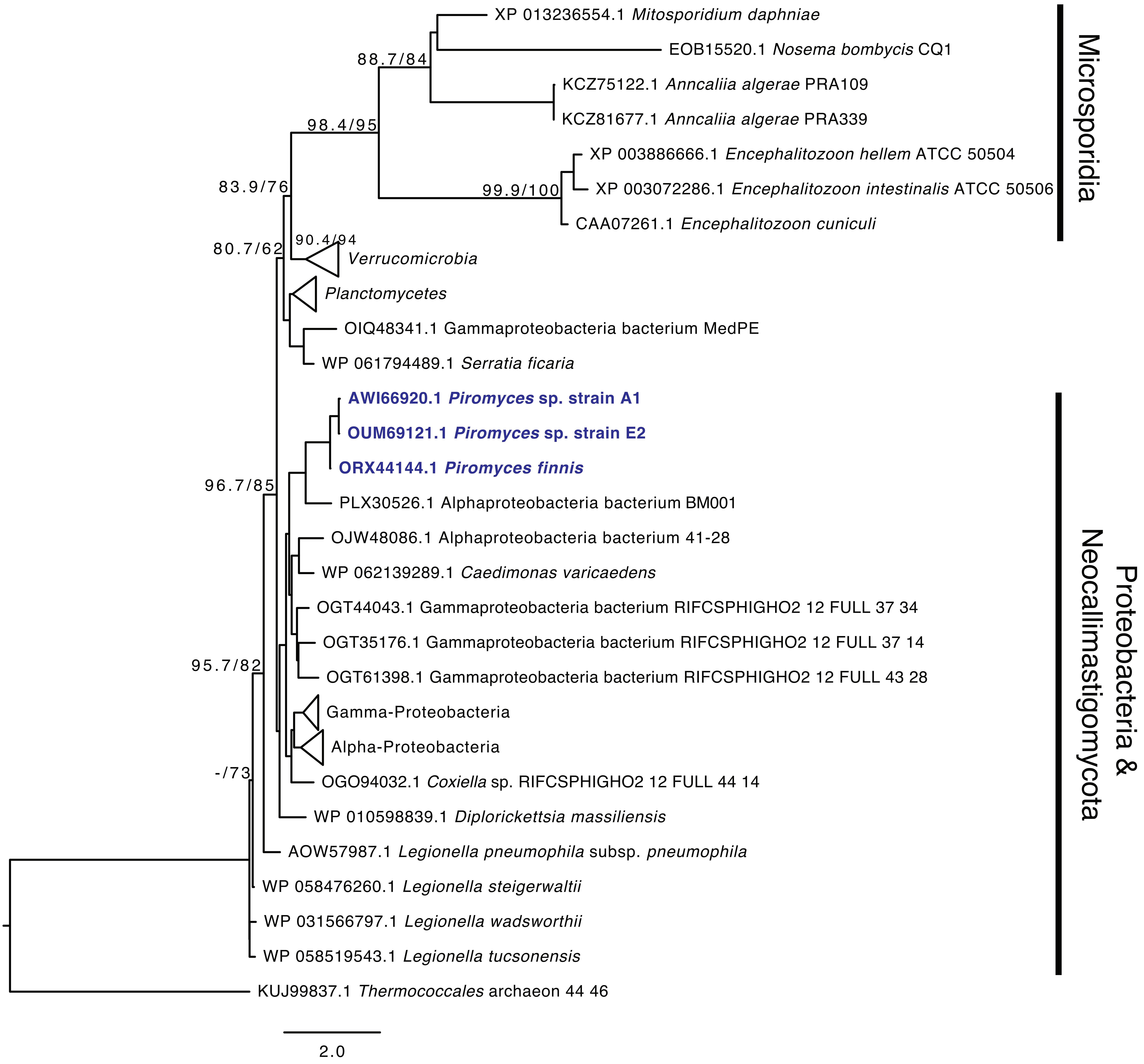

Fig S29. CDP-diacylglycerol--serine-O-phosphatidyl transferase

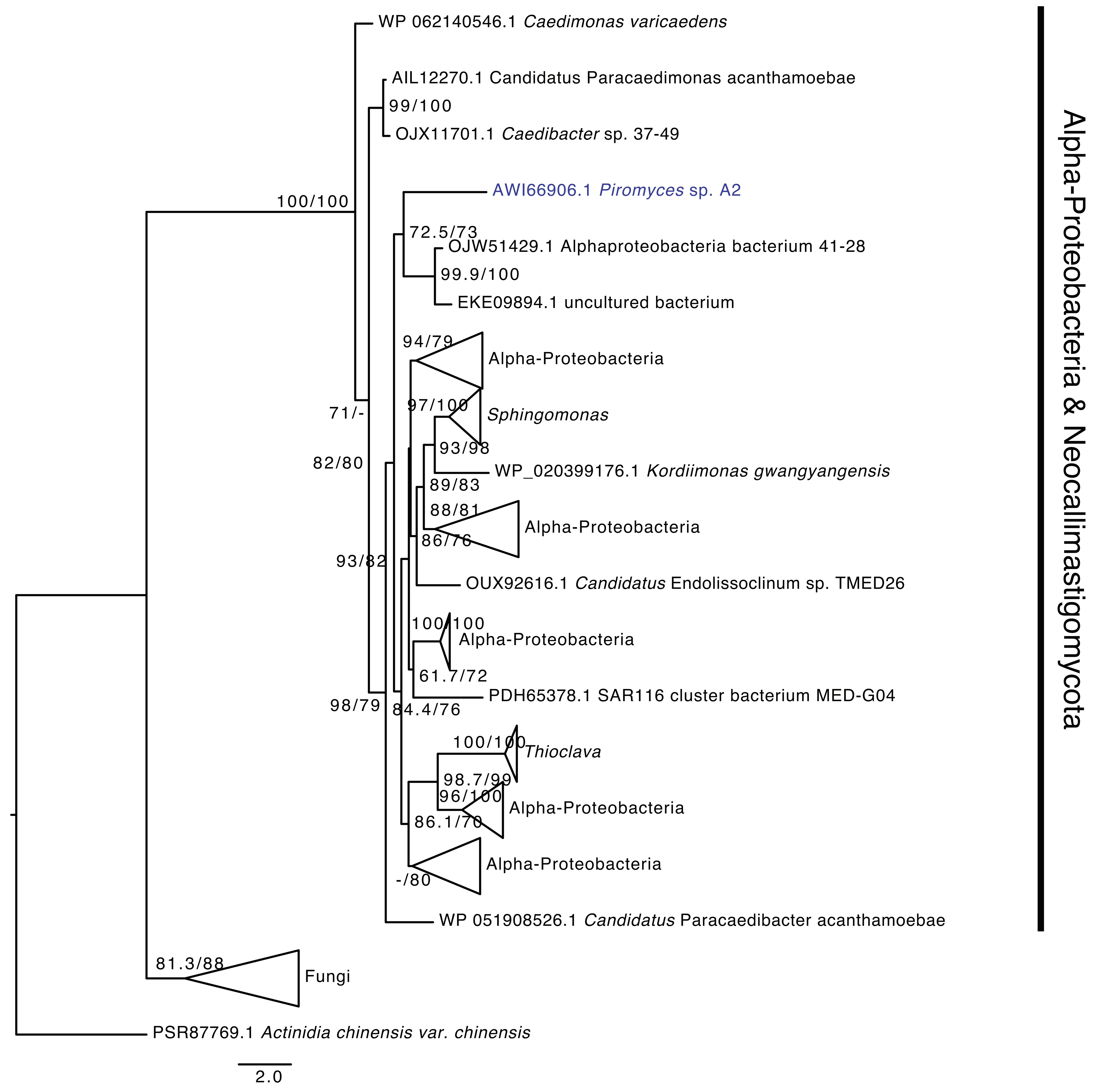

Fig S30. Methylene-fatty-acyl-phospholipid synthase

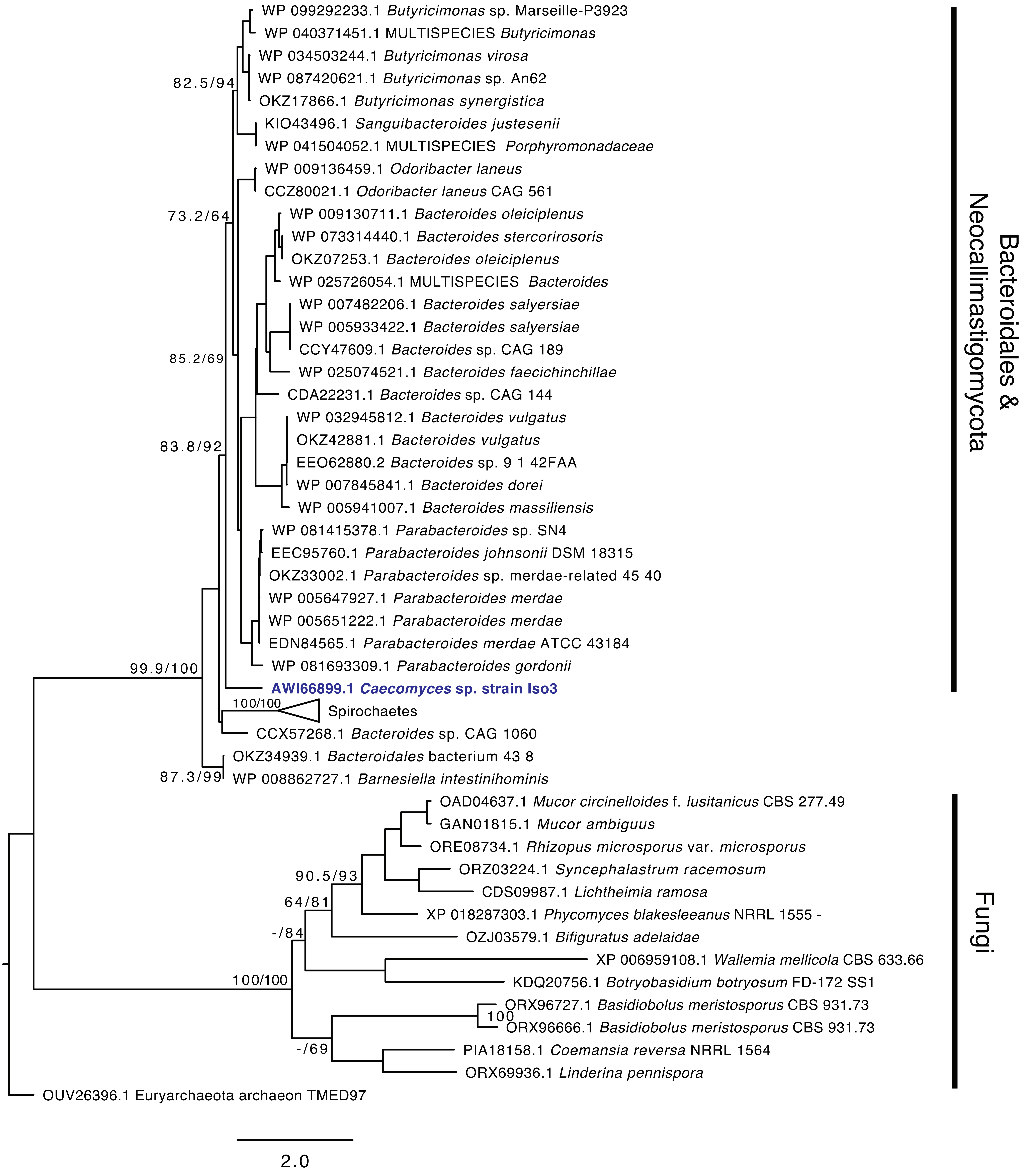

Fig S31. Glyoxalase I

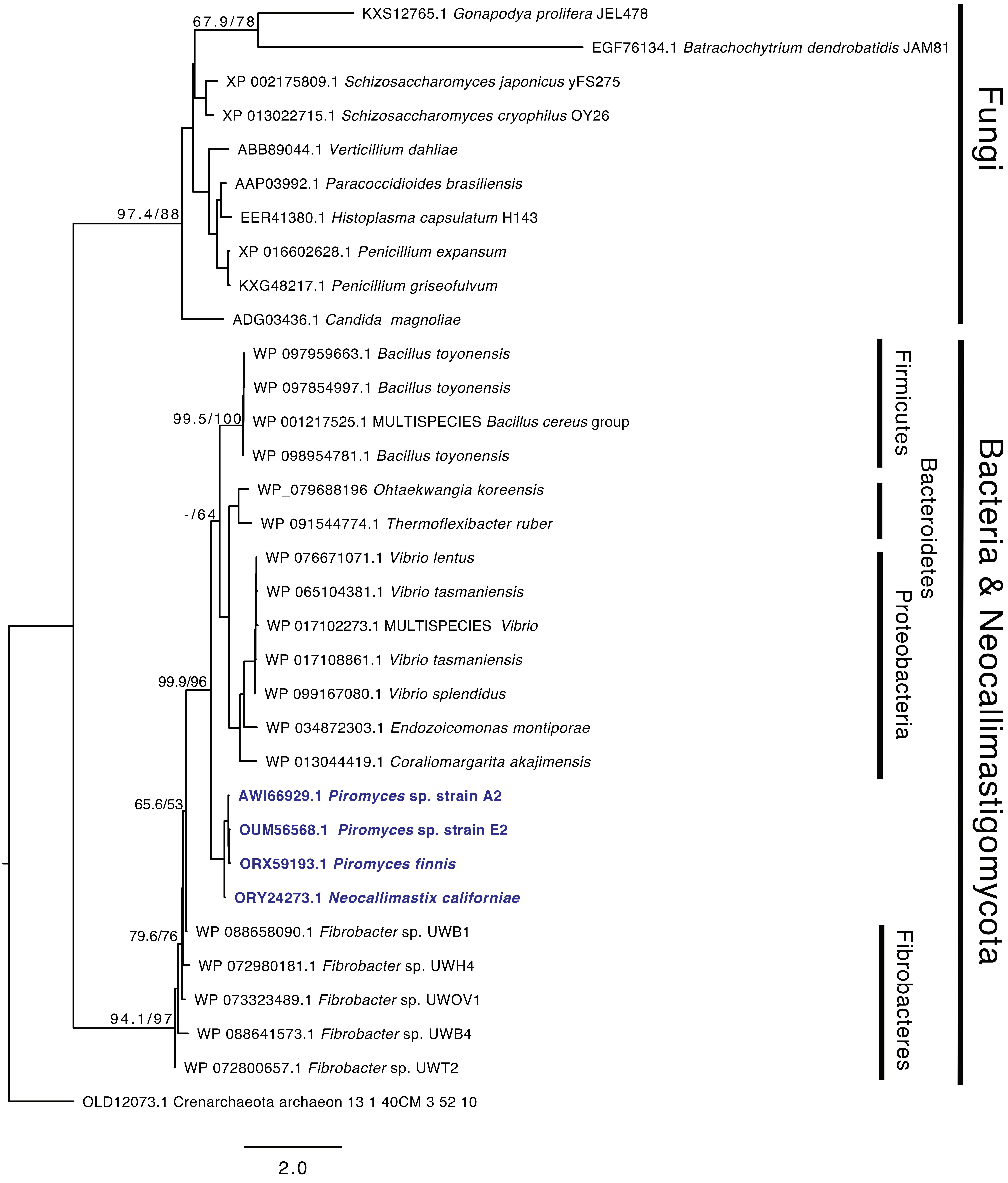

Fig S32. Glyoxalase II

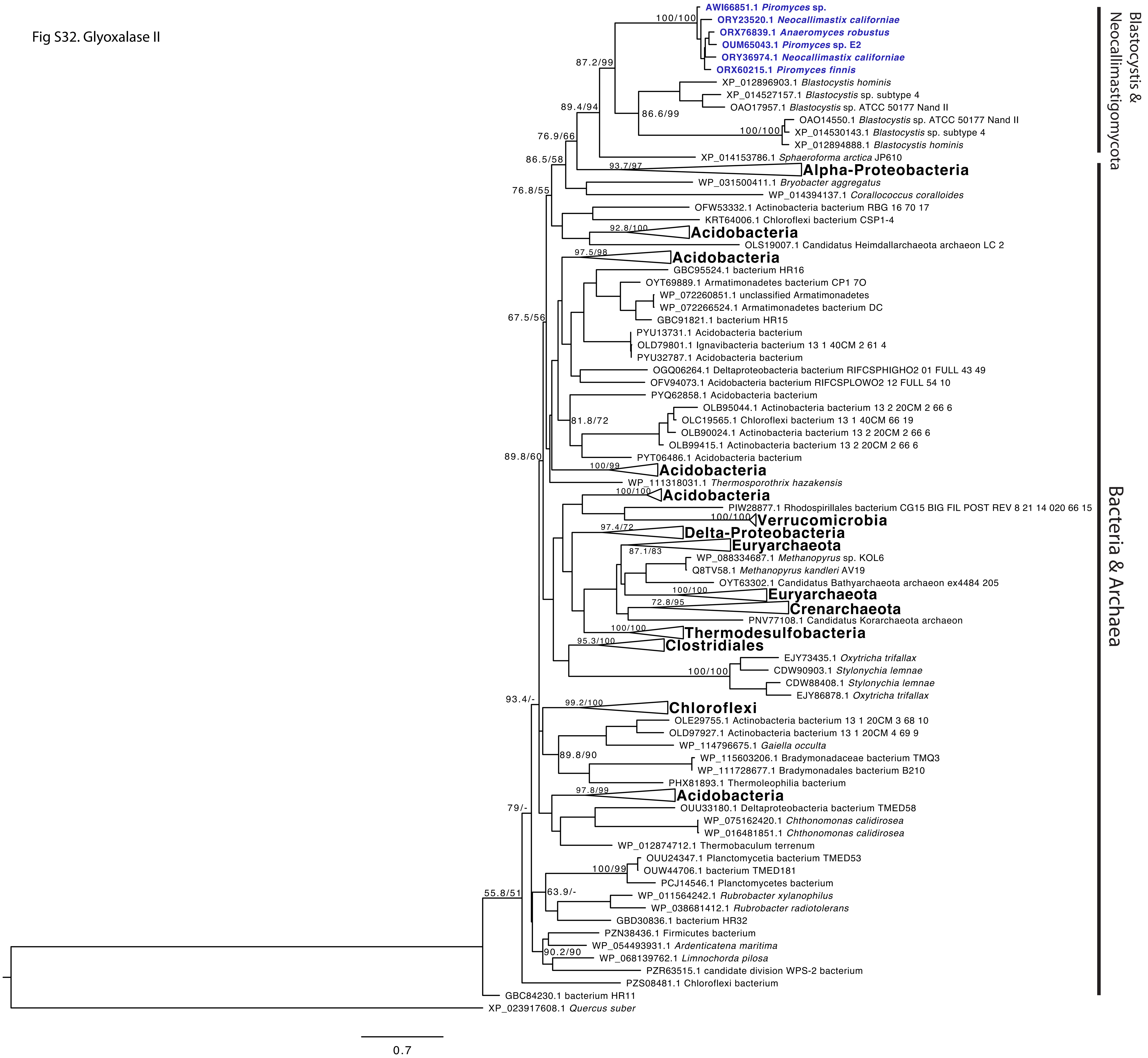

Fig S33. Squalene-hopene cyclase

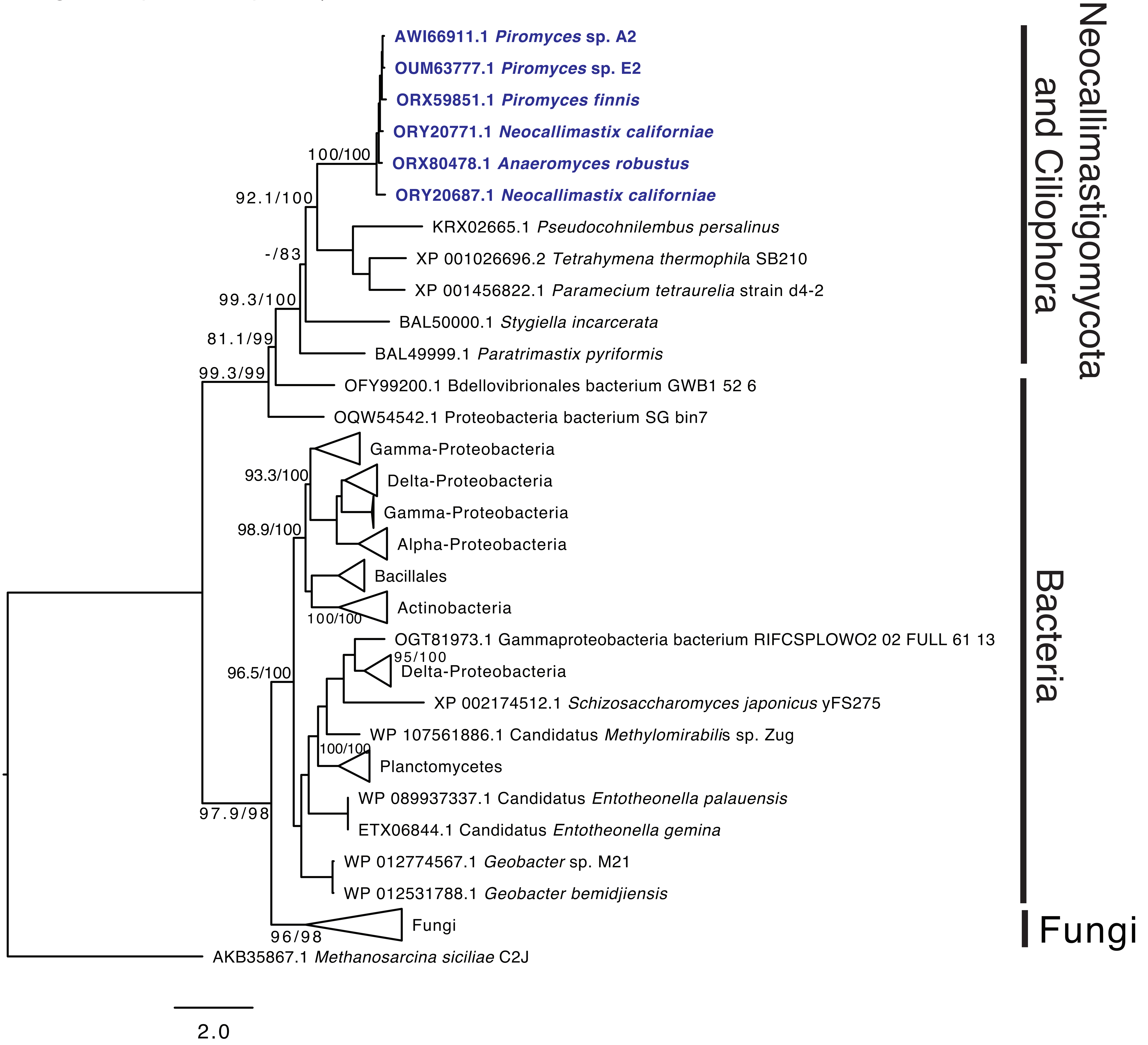

Fig S34. Superoxide dismutase Fe/Mn type

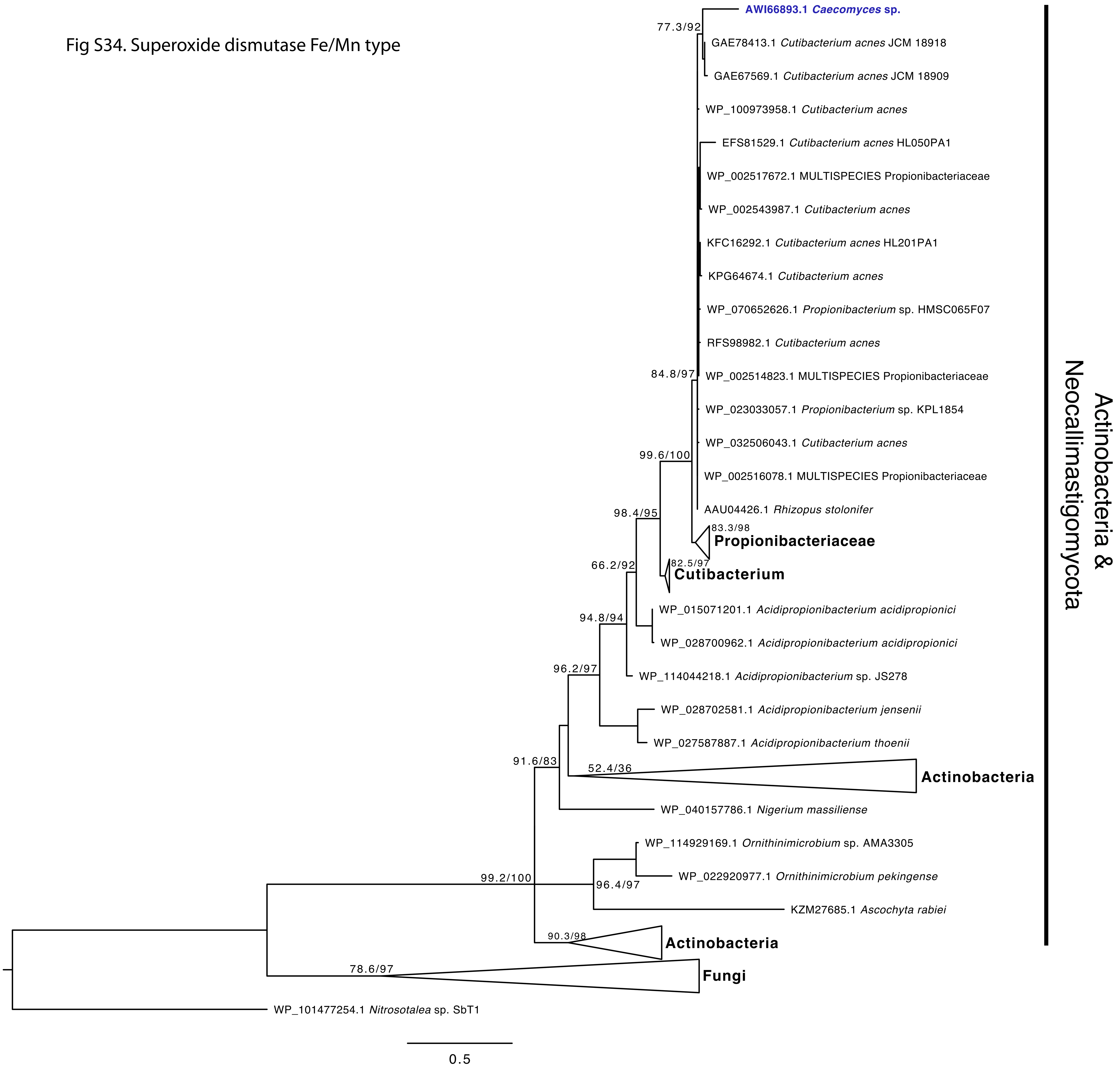

Fig S35. Glutathione peroxidase

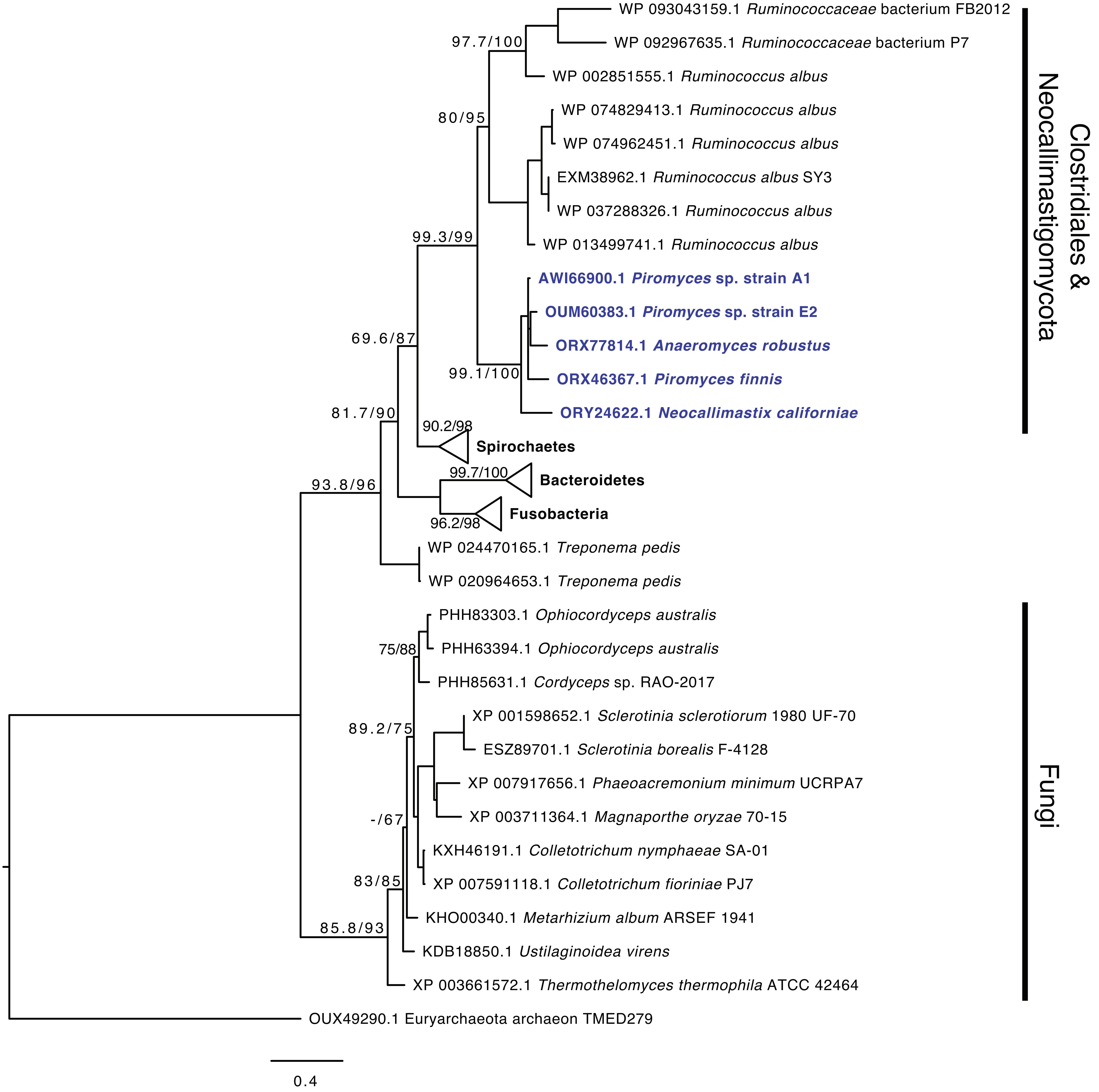

Fig S36. Rubrerythrin

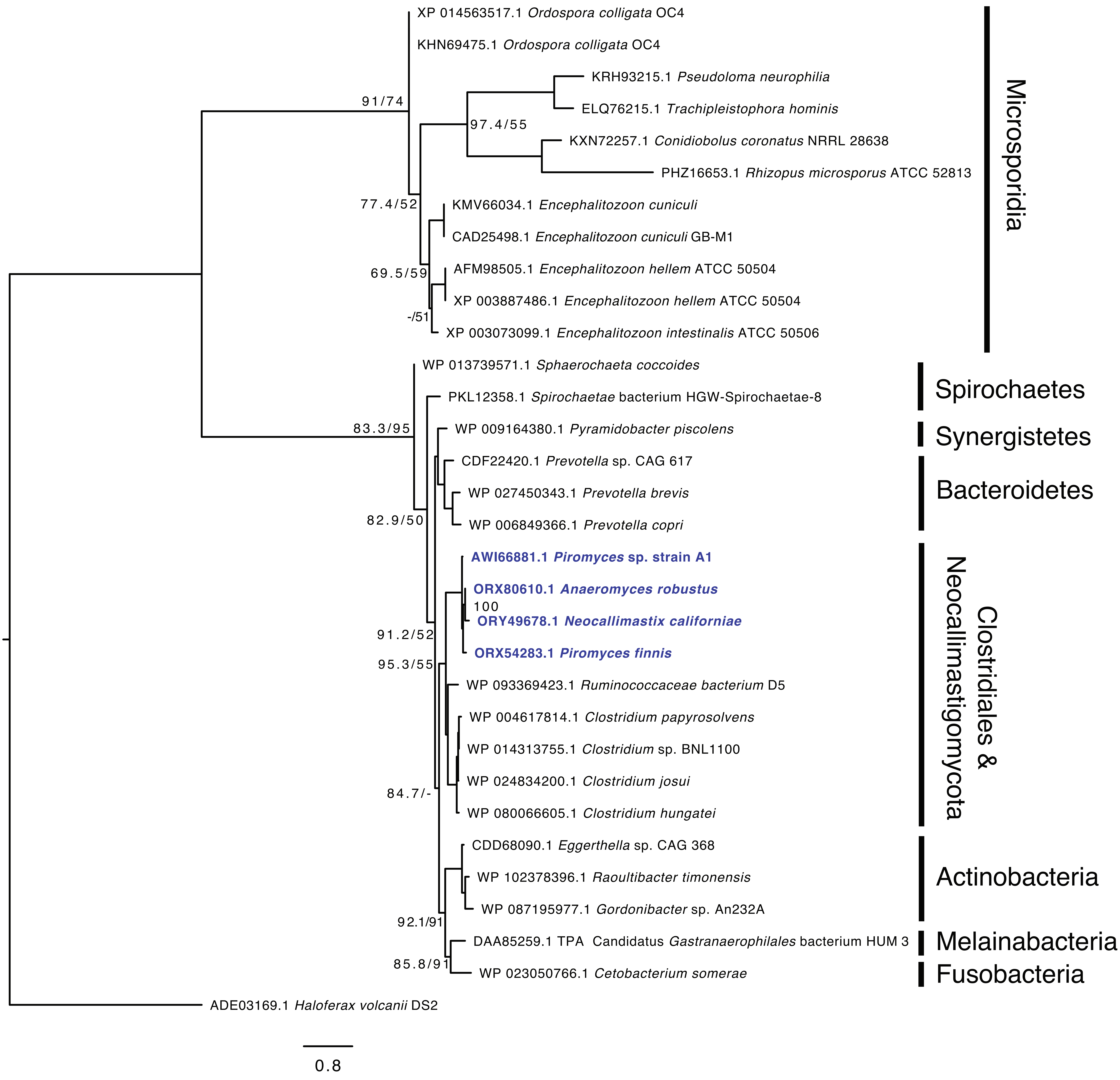

Fig S37. Alkyl hydroperoxidase

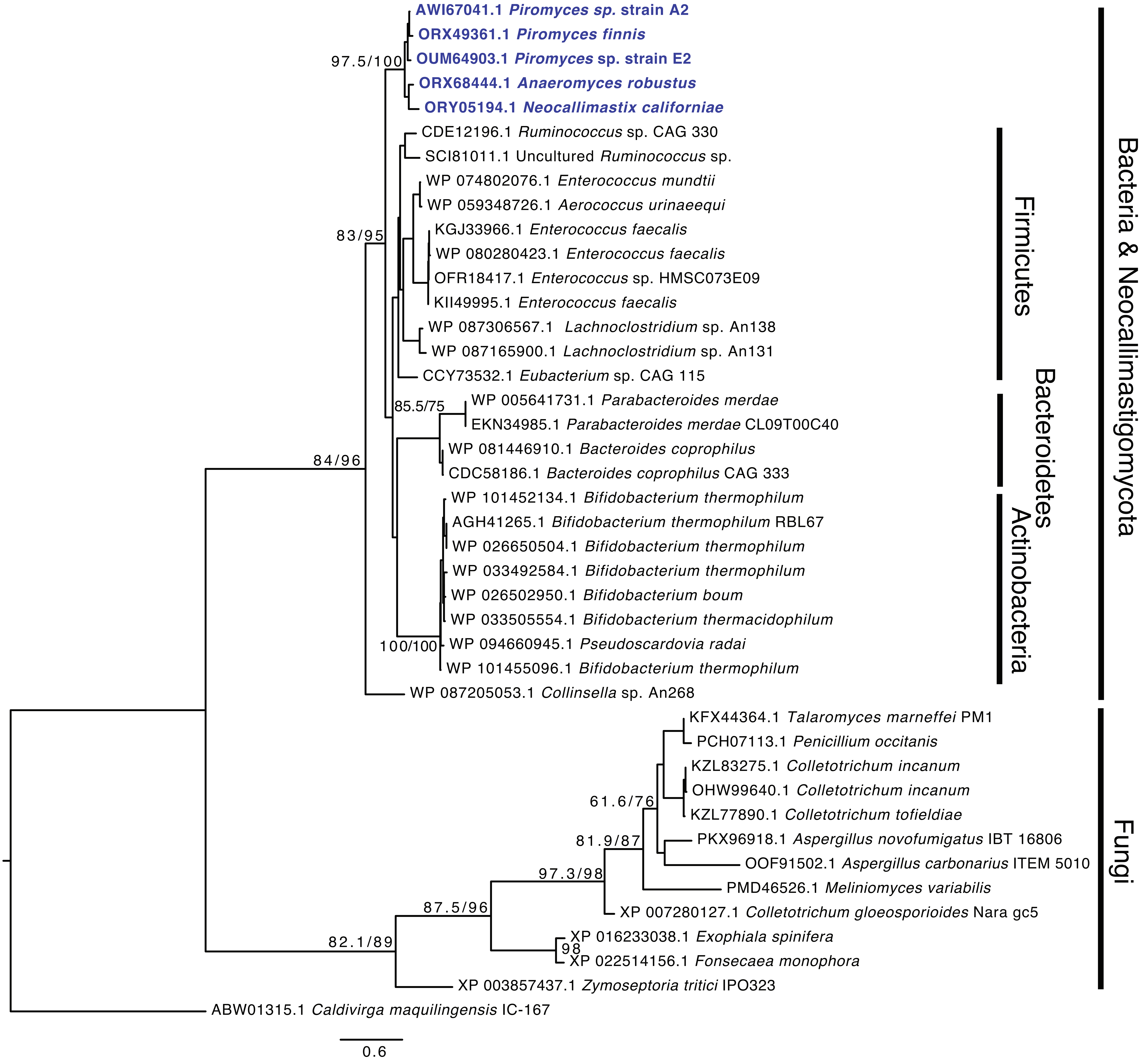

Fig S38. Phosphoglycolate phosphatase

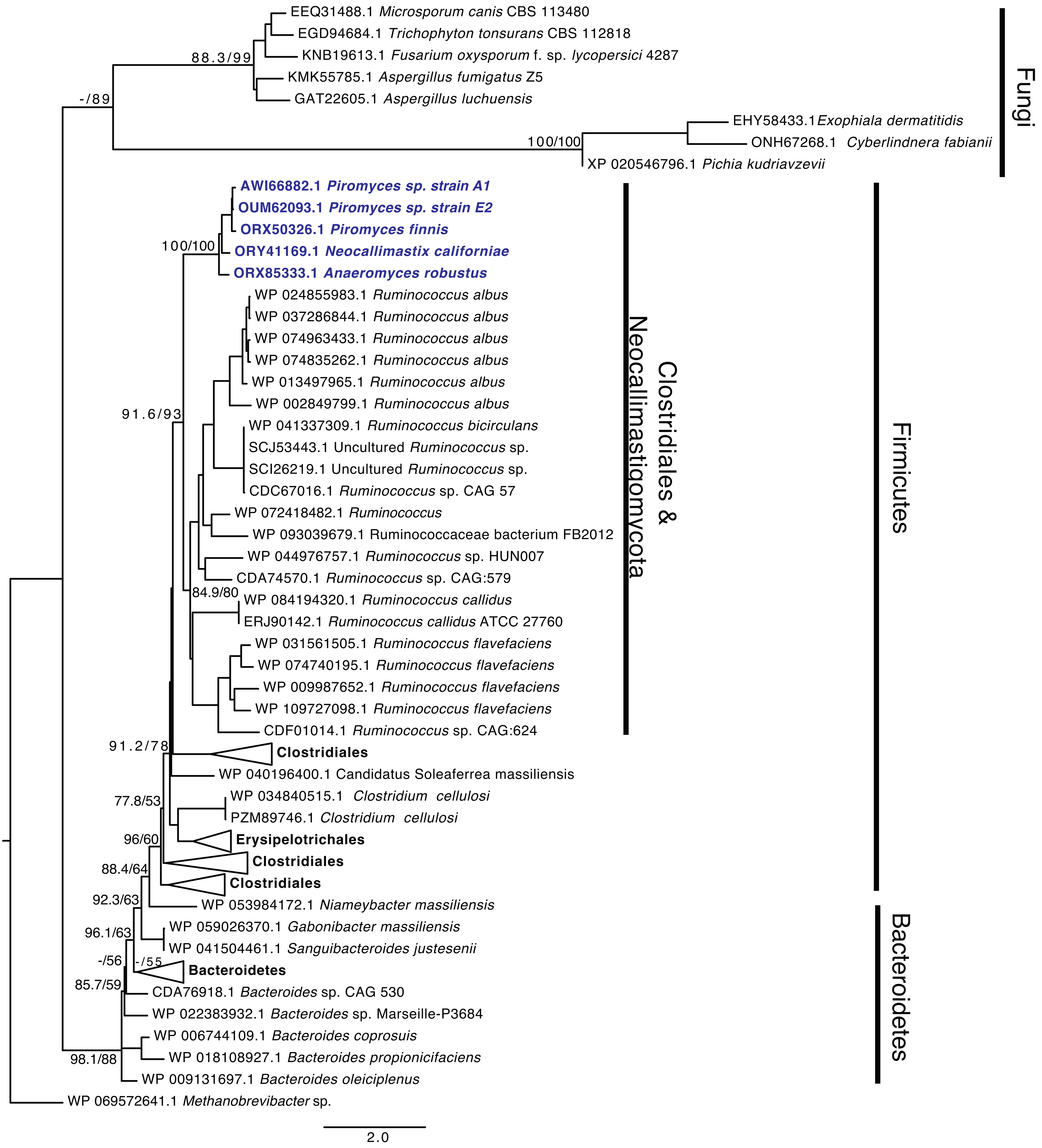

Fig S39. Chloramphenicol acetyltransferase

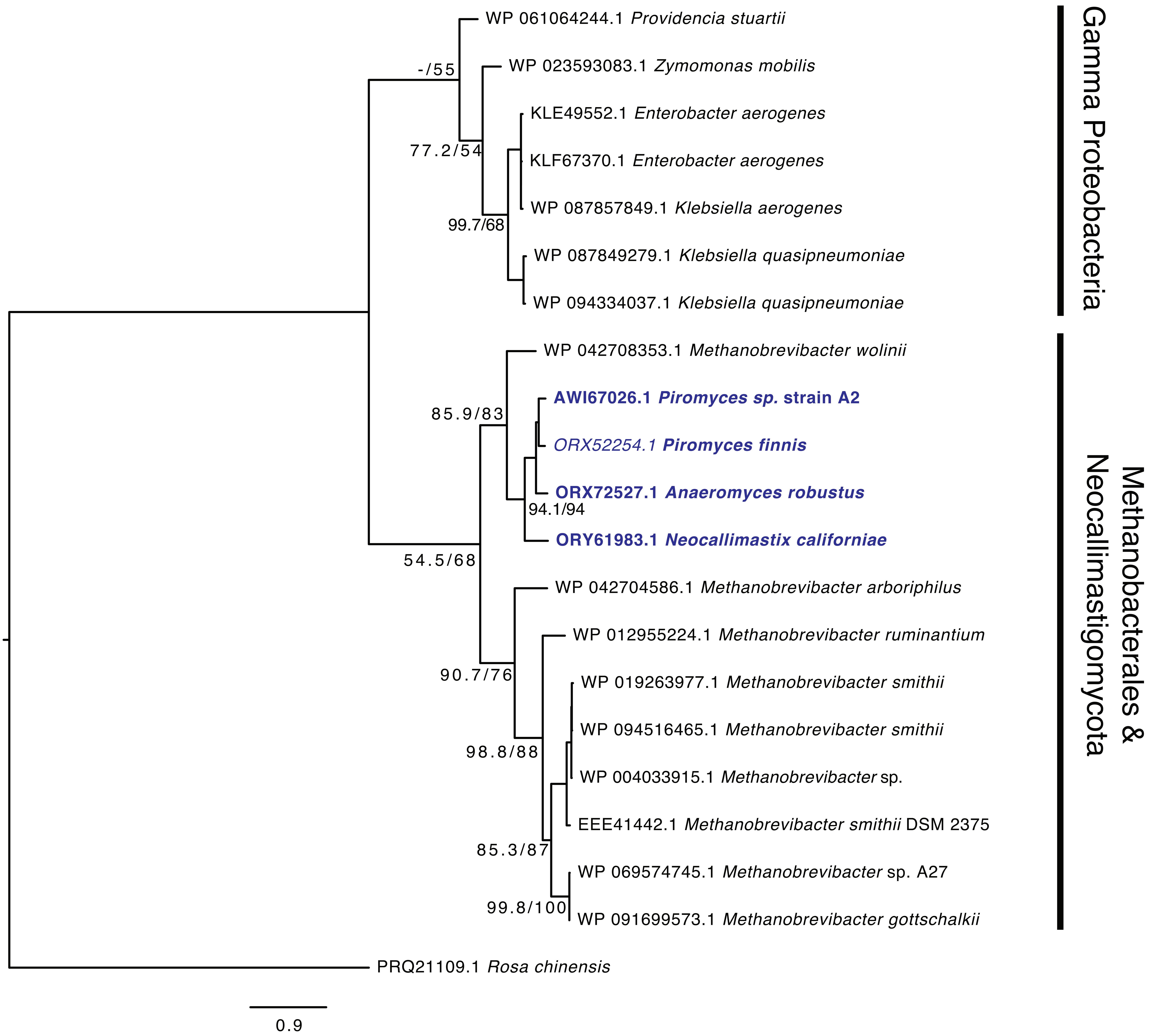

Fig S40. Aminoglycoside Phosphotransferase

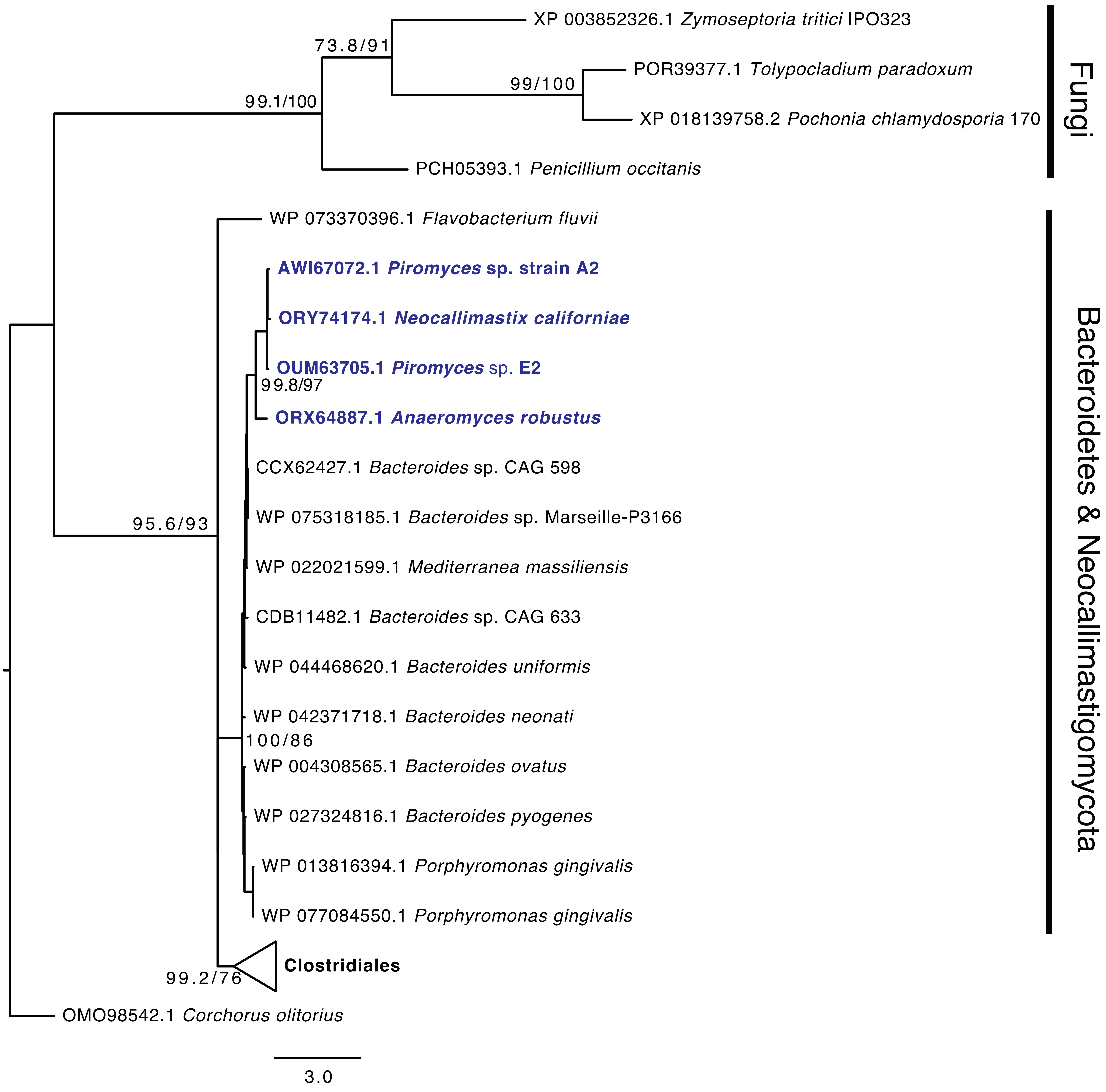

Fig S41. DNA-3-methyl adenine glycosylase I

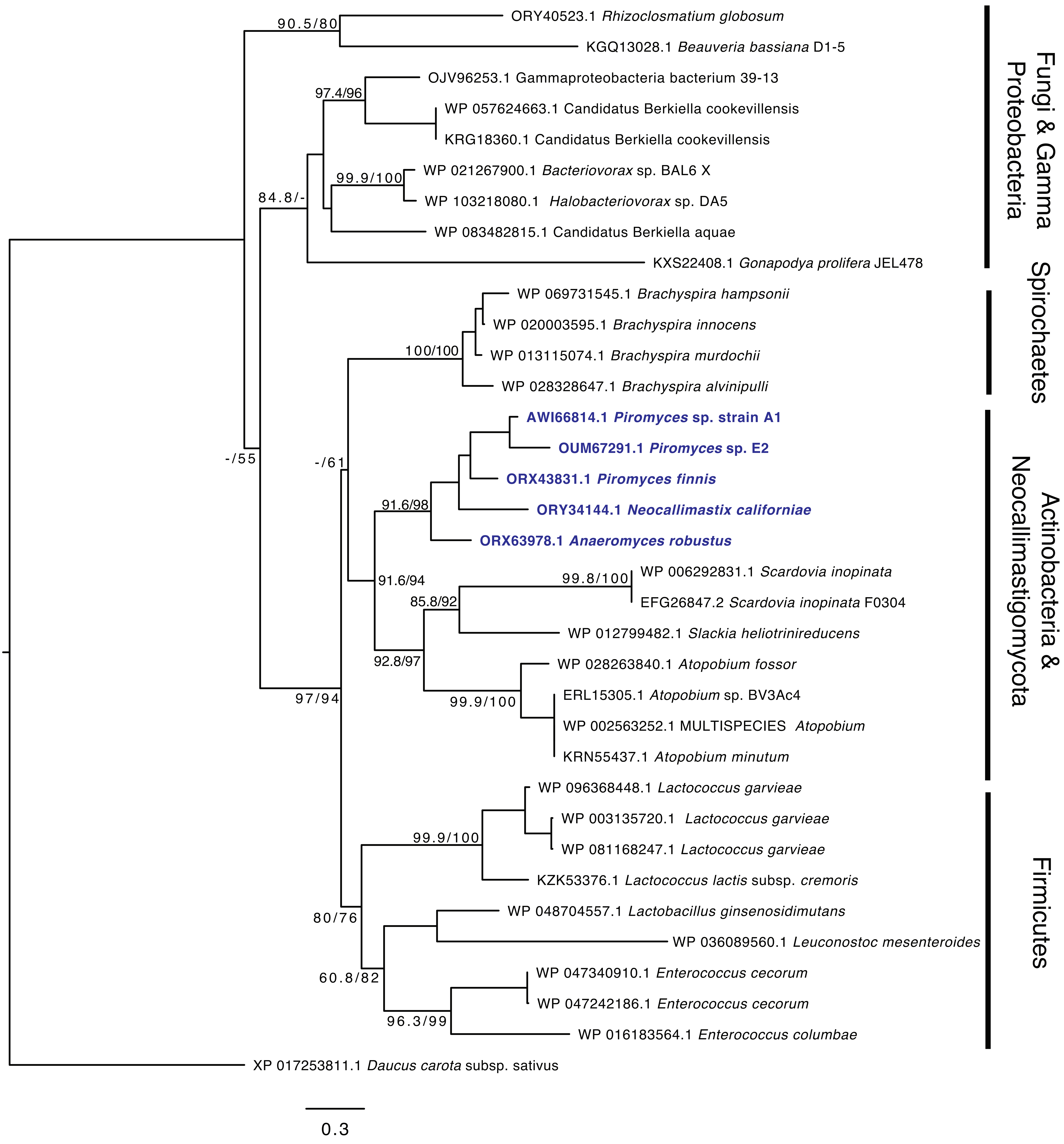

Fig S42. Methylated DNA protein-cysteine methyl transferase

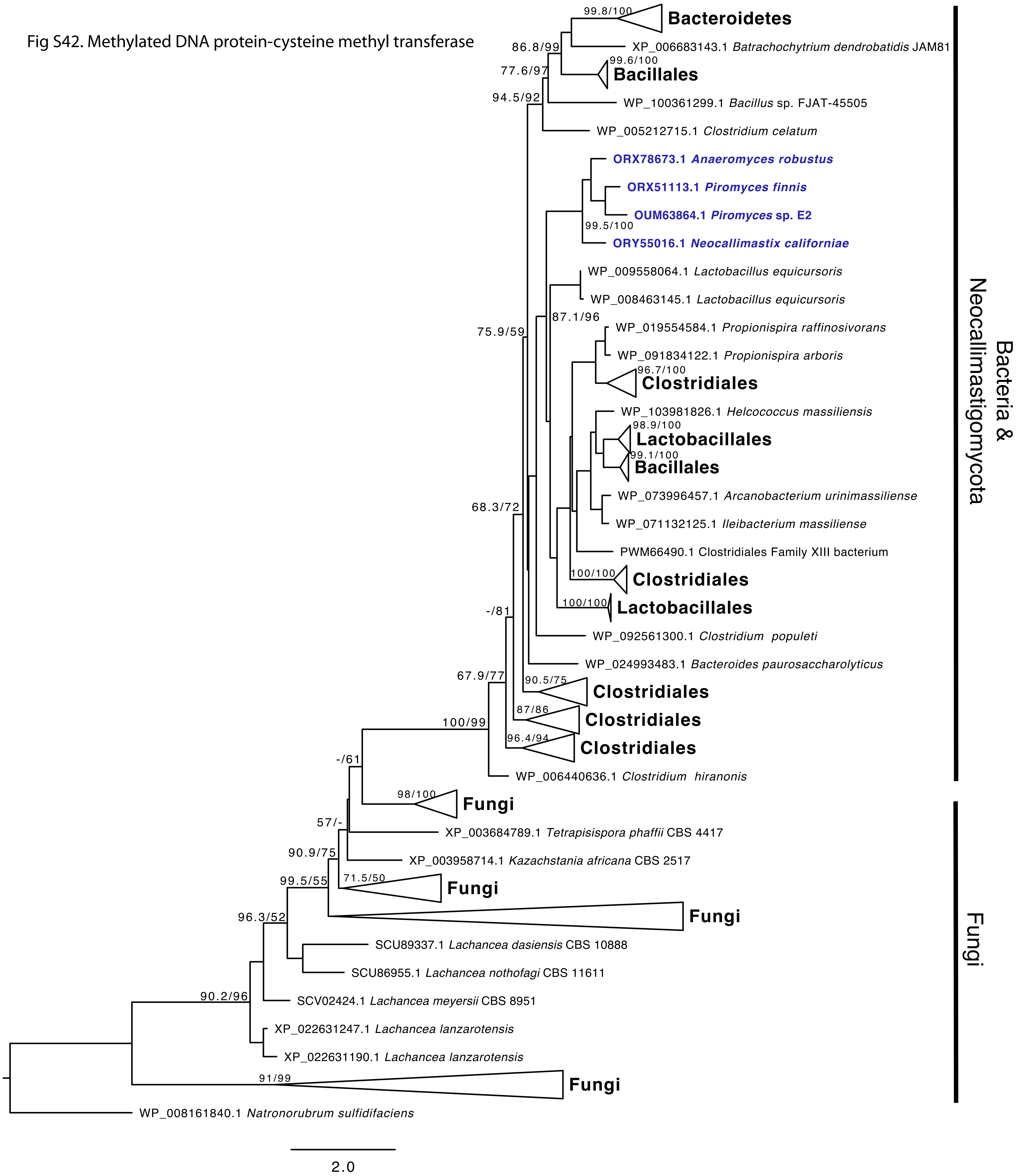

Fig S43. Galactoside-O-acetyl transferase

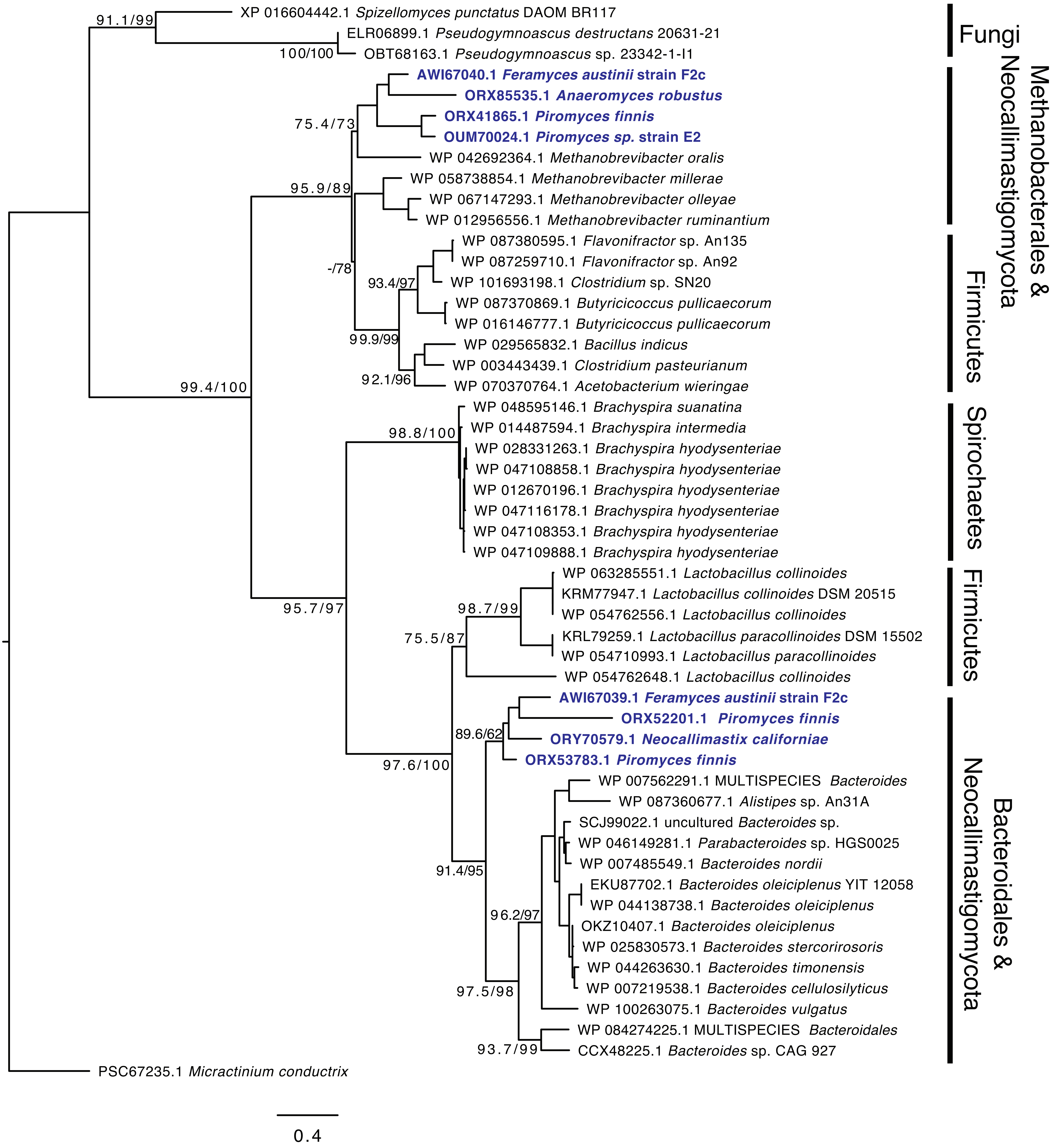

Fig S44. Maltose acetyl transferase

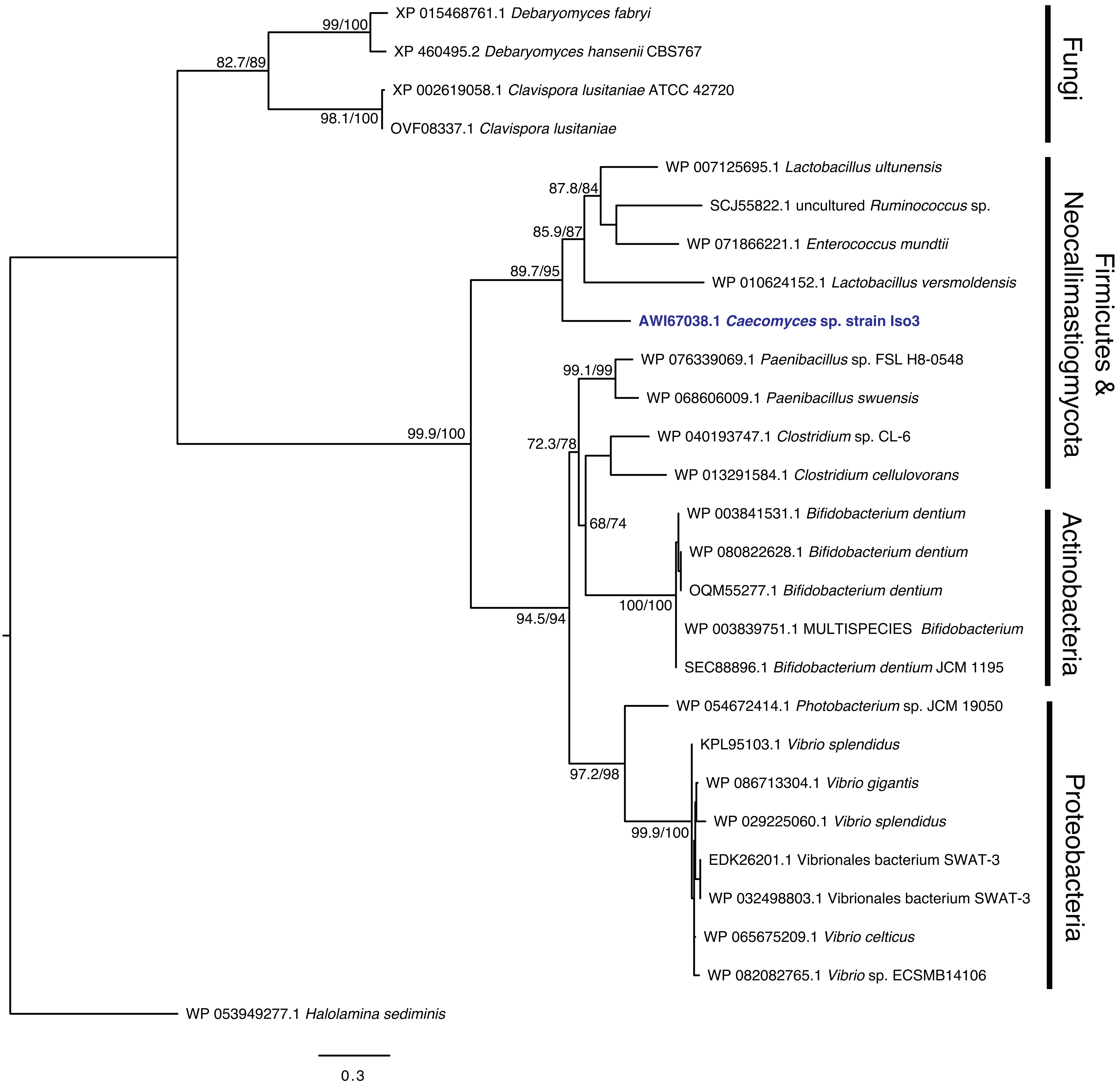

Fig S45. Methionine-R-sulfoxide reductase

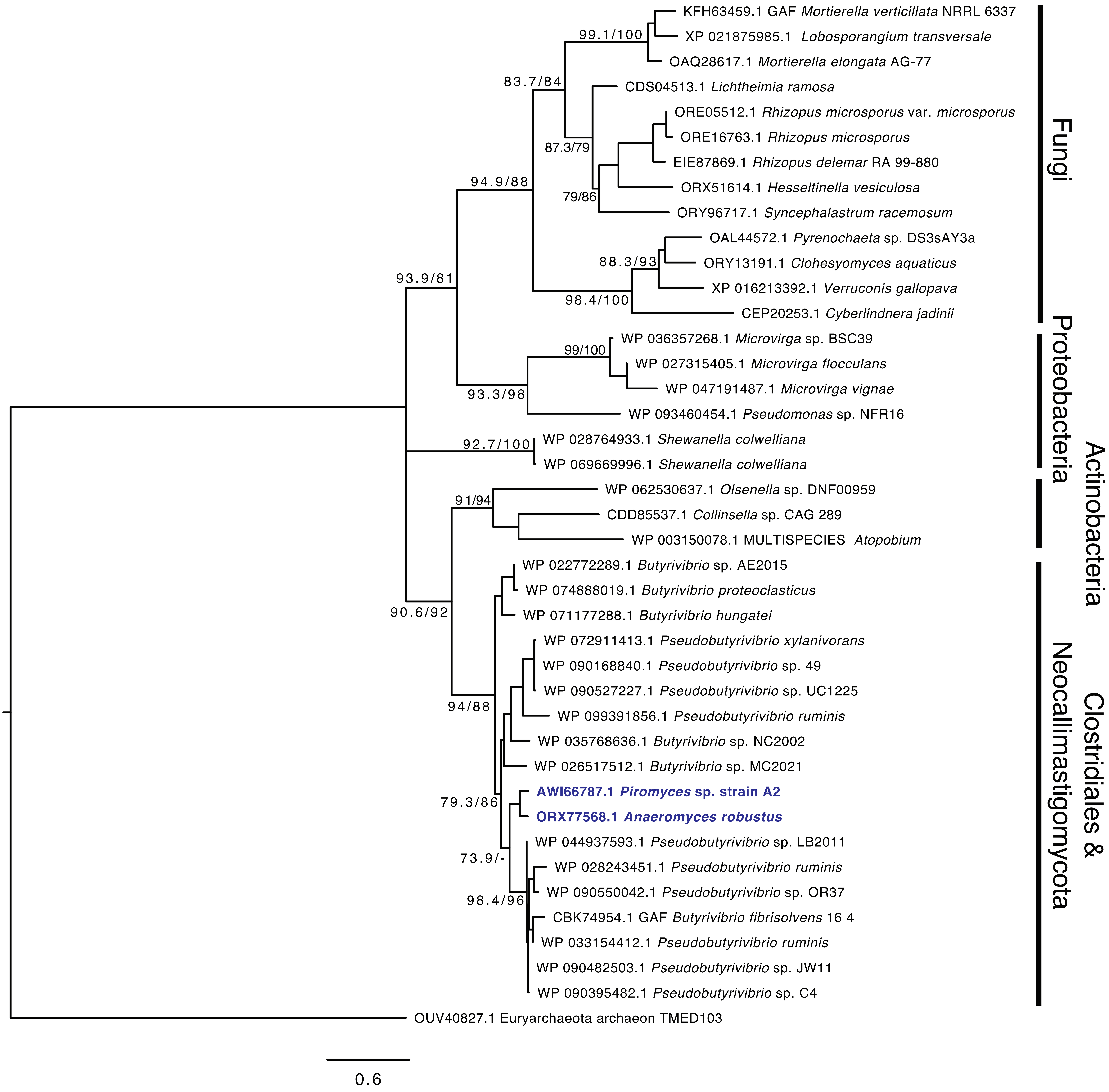

**Figure S46.** Cartoon depicting the impact HGT had on the ability of AGF to degrade several plant polysaccharides including (A) cellulose, (B) glucoarabinoxylans/ glucuronoarabinoxylan, (C) galactoglucomannan, (D) xyloglucan, (E) lichenesin (β-1,3 and β-1,4), (F) laminarin (β-1,3 and β-1,6), (G) starch (α-1,4 and α-1,6), (H) chitin, (I) trehalose, (J) poly-N-acetyl-α-D-galactosamine, (K) peptidoglycan, (L) rhamnogalactouronan, and (M) homogalactouronan. The major CAZymes identified in the transcriptomes studied are shown, with the highlight color indicating the extent of HGT in that CAZy family as follows; (1) 100% of transcripts identified belonging to families highlighted in red were horizontally transferred, (2) the majority (>50%) of transcripts identified belonging to families highlighted in orange were horizontally transferred, (3) a small fraction (< 50%) of transcripts identified belonging to families highlighted in light blue were horizontally transferred, and (4) 100% of transcripts identified belonging to families highlighted in dark blue were of fungal origin.

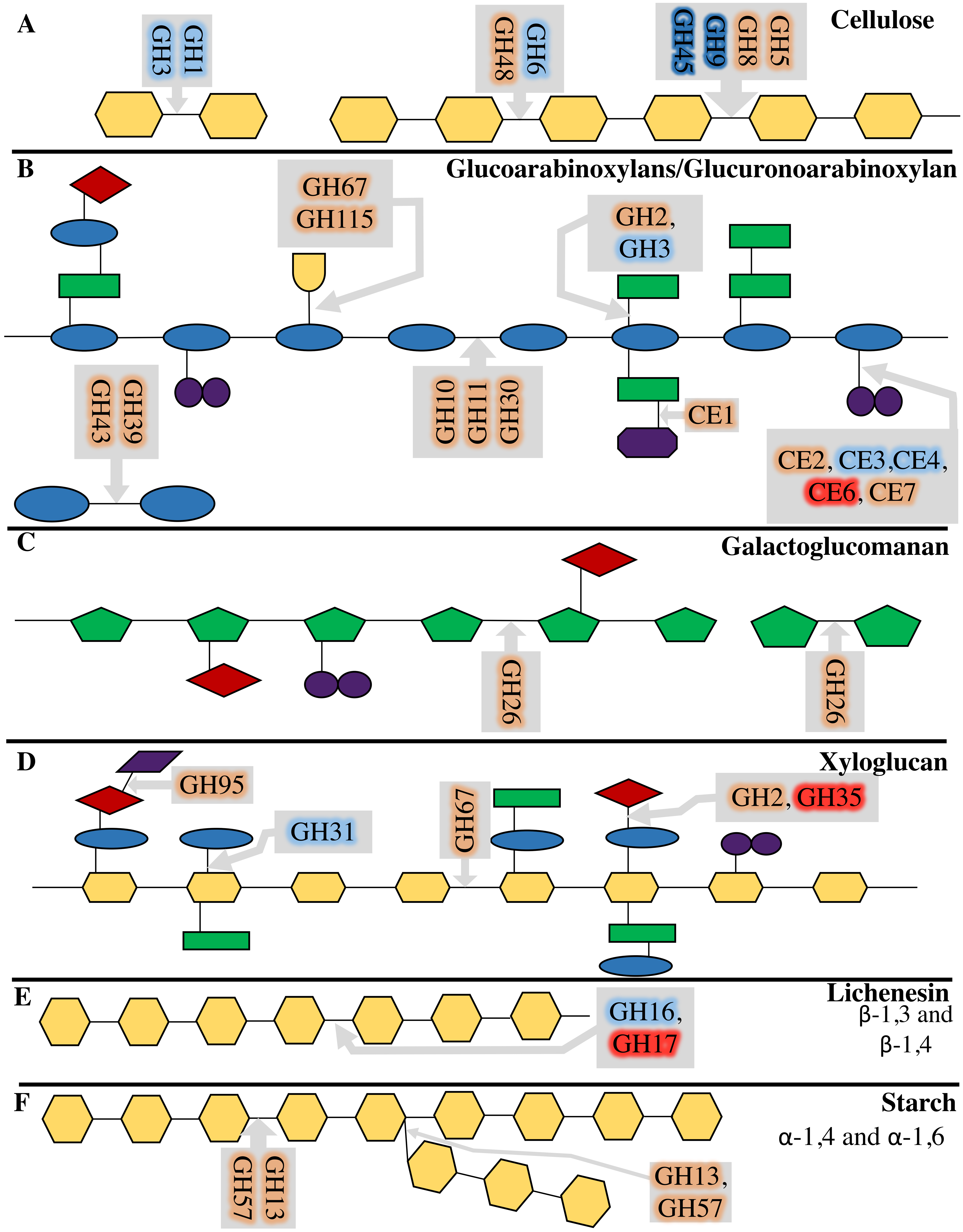
A
Cellulose
GH1
GH3
GH6
GH48
GH5
GH8
GH9
GH45
B
Glucoarabinoxylans/Glucuronoarabinoxylan
GH67
GH115
GH2,
GH3
GH43
GH39
GH10
GH11
GH30
CE1
CE2, CE3,CE4,
CE6, CE7
C
Galactoglucomanan
GH26
GH26
D
Xyloglucan
GH95
GH31
GH67
GH2, GH35
E
Lichenesin
β-1,3 and
β-1,4
GH16,
GH17
F
Starch
α-1,4 and α-1,6
GH13
GH57
GH13,
GH57

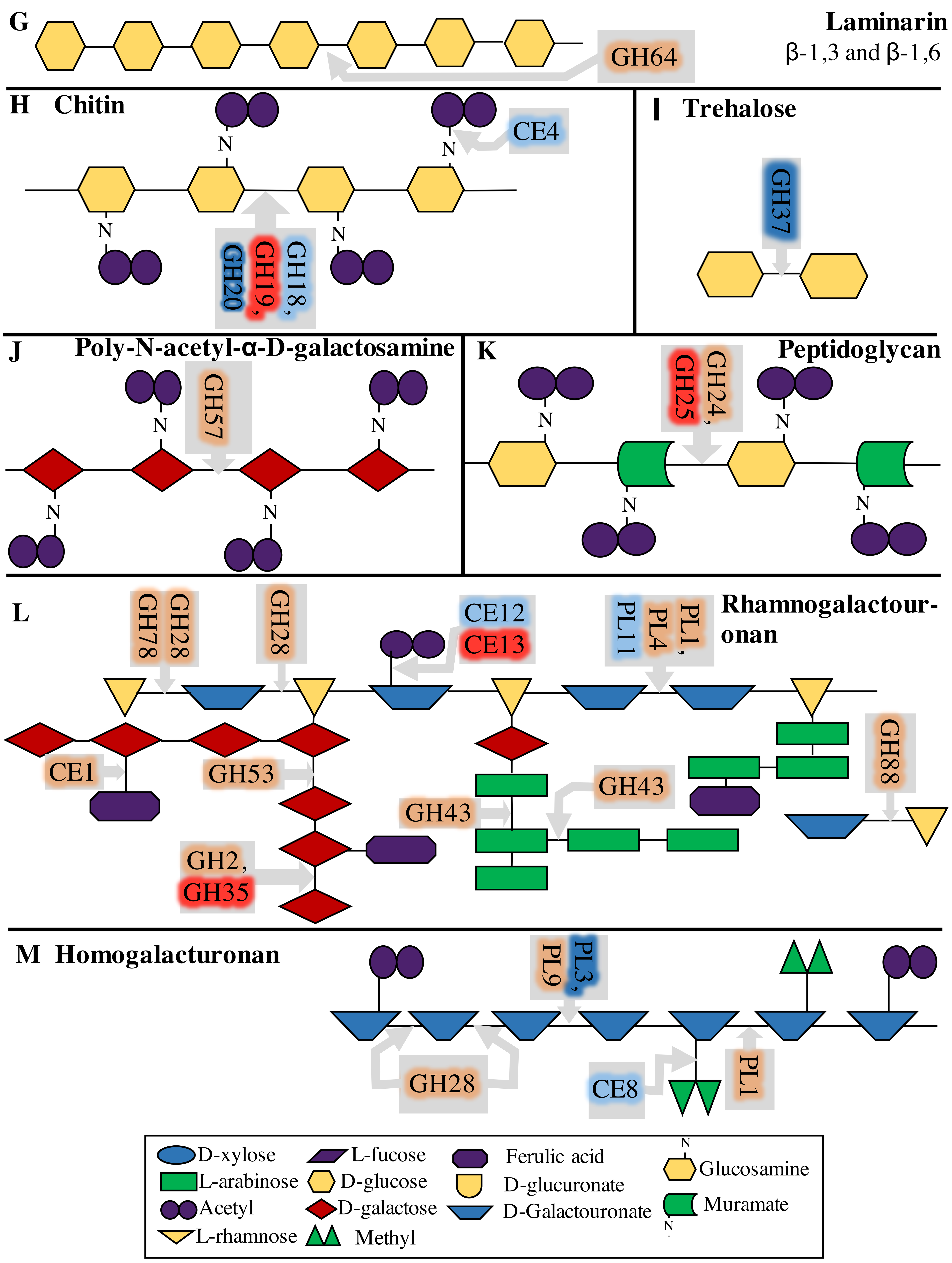
G
Laminarin
β-1,3 and β-1,6
GH64
H Chitin
CE4
GH18, GH19, GH20
I Trehalose
GH37
J Poly-N-acetyl-α-D-galactosamine
GH57
K
Peptidoglycan
GH24, GH25
L
Rhamnogalactour-onan
GH78
GH28
GH28
CE12
CE13
PL1, PL4, PL11
CE1
GH53
GH43
GH43
GH88
GH2, GH35
M Homogalacturonan
PL3, PL9
GH28
CE8
PL1
D-xylose
L-arabinose
Acetyl
L-rhamnose
L-fucose
D-glucose
D-galactose
Methyl
Ferulic acid
D-glucuronate
D-Galactouronate
Glucosamine
Muramate
